# The K2: Open-source simultaneous triple-color TIRF microscope for live-cell and single-molecule imaging

**DOI:** 10.1101/2022.12.19.521031

**Authors:** Christian Niederauer, Marco Seynen, Jan Zomerdijk, Marko Kamp, Kristina A. Ganzinger

## Abstract

Imaging the dynamics and interactions of biomolecules at the single-molecule level in live cells and reconstituted systems has generated unprecedented knowledge about the biomolecular processes underlying many cellular functions. To achieve the speed and sensitivity needed to detect and follow individual molecules, these experiments typically require custom-built microscopes or custom modifications of commercial systems. The costs of such single-molecule microscopes, their technical complexity and the lack of open-source documentation on how to build custom setups therefore limit the accessibility of single-molecule imaging techniques. To advance the adaptation of dynamic single-molecule imaging by a wider community, we present the “K2”: an open-source, simultaneous triple-color total internal reflection fluorescence (TIRF) microscope specifically designed for live-cell and single-molecule imaging. We explain our design considerations and provide step-by-step building instructions, parts list and full CAD models. The flexible design of this TIRF microscope allows users to customize it to their scientific and financial needs, or to re-use parts of our design to improve the capabilities of their existing setups without necessarily having to build a full copy of the K2 microscope.

## Introduction

Commercially available fluorescence microscopes for state-of-the-art single-molecule and super-resolution microscopy are often expensive to acquire and lack the flexibility to incorporate custom features beyond already available add-ons. In recent years, a number of open-source projects originated within the super-resolution fluorescence microscopy community, facilitating the design and construction of custom microscopes from the bottom-up. [1–7]. Additionally, stand-alone solutions for various sub-components of single-molecule microscopes have been published as open source hardware projects (*e.g.* focus stabilization [8–10], laser sources [11–14], objective heaters [15, 16] and beam-shaping devices [12, 17–19]). Our microscope design builds on previous open-source projects and our in-house engineering expertise, to provide a cost-effective and flexible solution for single-molecule fluorescence microscopy.

Here, we present a single-molecule total internal reflection fluorescence (TIRF) microscope that is capable of acquiring dynamic single-molecule data with four-color excitation and simultaneous triple-color detection. The microscope uses a single sCMOS camera and projects the fluorescence emission coming from the sample onto different camera regions for each fluorescence channel. Together with homogeneous illumination from a beamshaping device, this enables simultaneous three-color imaging with large fields of view (73 µmx73 µm) for each channel. The microscope further features objective heating, a plexiglass cover with an integrated LED array for brightfield illumination, and motorized switching of the angle of the excitation laser beam from TIRF to highly inclined and laminated optical sheet (HILO) and epifluorescence. Finally, the microscope also includes a flip-in lens for fluorescence recovery after photobleaching (FRAP) experiments and a focus stabilization system to correct axial sample drift in timelapse and multi-position measurements.

#### Box 1. Potential uses of the setup

**Box 1.**
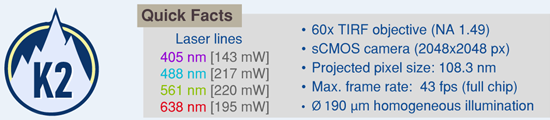
The K2: Open-source simultaneous triple-color TIRF microscope for live-cell and single-molecule imaging.

- (Single-molecule) TIRF microscopy with simultaneous detection of up to three colors
- Experiments with environmental control (temperature) or monitoring (temperature, humidity, sample drift)
- Video-rate recording with camera-triggered excitation (alternating or continuous)
- Timelapse- and multiposition-acquisitions with focus stabilization
- LED-based brightfield imaging
- Widefield epifluorescence imaging
- Fluorescence recovery after photobleaching (FRAP) experiments

Not demonstrated but possible:

- Single-molecule Förster resonance energy transfer (FRET) microscopy
- Multi-color super-resolution microscopy: DNA-PAINT, Photo-activated localization microscopy (PALM), stochastic optical reconstruction microscopy (STORM)

To ensure reliability and reproducibility of experiments, all imaging settings (*e.g.* laser power, exposure time, stage position, angle of incidence …) are carefully documented for every dataset. We also monitor and record environmental parameters (temperature, humidity) and the axial sample drift when using the focus stabilization system. This allows us to precisely control and replicate our experiments under identical imaging conditions, which is essential for generating comparable and reproducible data with TIRF microscopes [20].

### Setup description

The K2 single-molecule TIRF microscope is comprised of a central cube with the sample stage, sample holder and objective (see ***Figure 1*** and panel I in ***Figure 2***), as well as two enclosed boxes. The box on the right-hand side of the central cube houses the excitation path optics and the focus lock system (see ***Figure 1*** and panel II and III in ***Figure 2***). The box to the left contains the detection pathway, where emitted fluorescence light from the sample is split in up to three spectrally separated channels (see ***Figure 1*** and panel IV in ***Figure 2***). The excitation and detection pathways are enclosed in light-tight aluminum boxes to reduce background light levels, minimize laser safety issues, and prevent dust buildup. The entire setup is mounted on a vibration-damped optical table, with additional components (*e.g.* pre-assembled laser-combiner box, panel V in ***Figure 2***) and the setup control computer located nearby.

**Figure 1.**
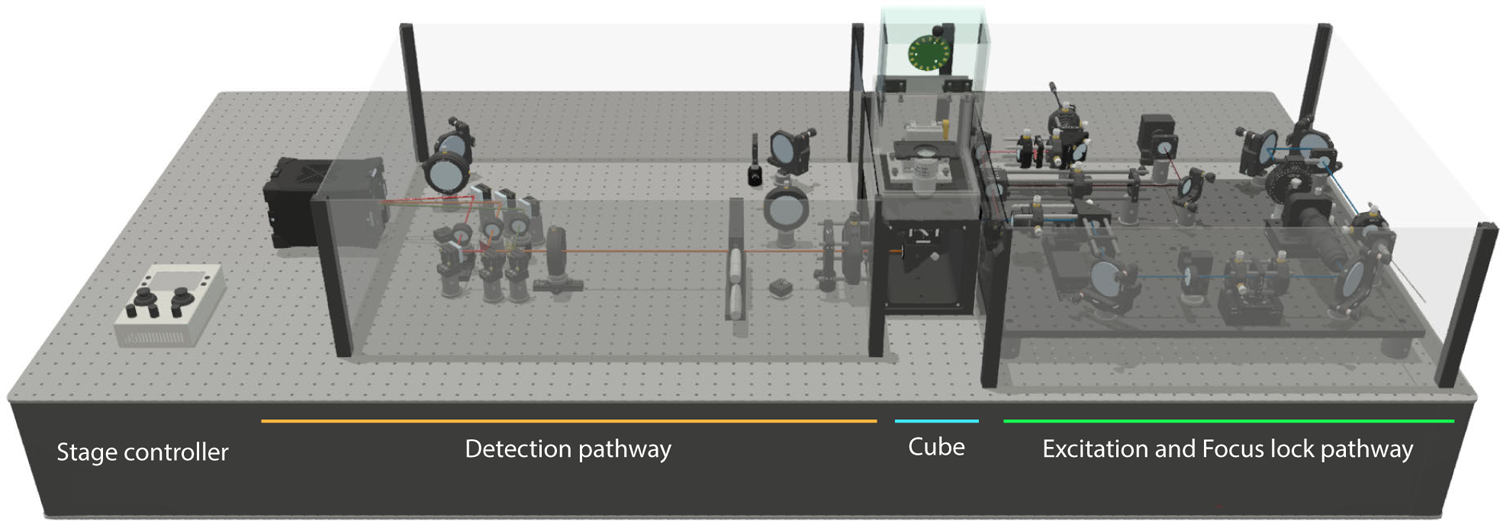
Render of the K2 open-source simultaneous triple-color TIRF microscope for live-cell and single-molecule imaging. For illustration purposes, the two enclosure boxes to the left and right of the central cube are rendered transparent and their lids are removed. The handheld sample stage joystick controller is placed on the optical bench only for illustration purposes and usually located on the computer desk.

**Figure 2.**
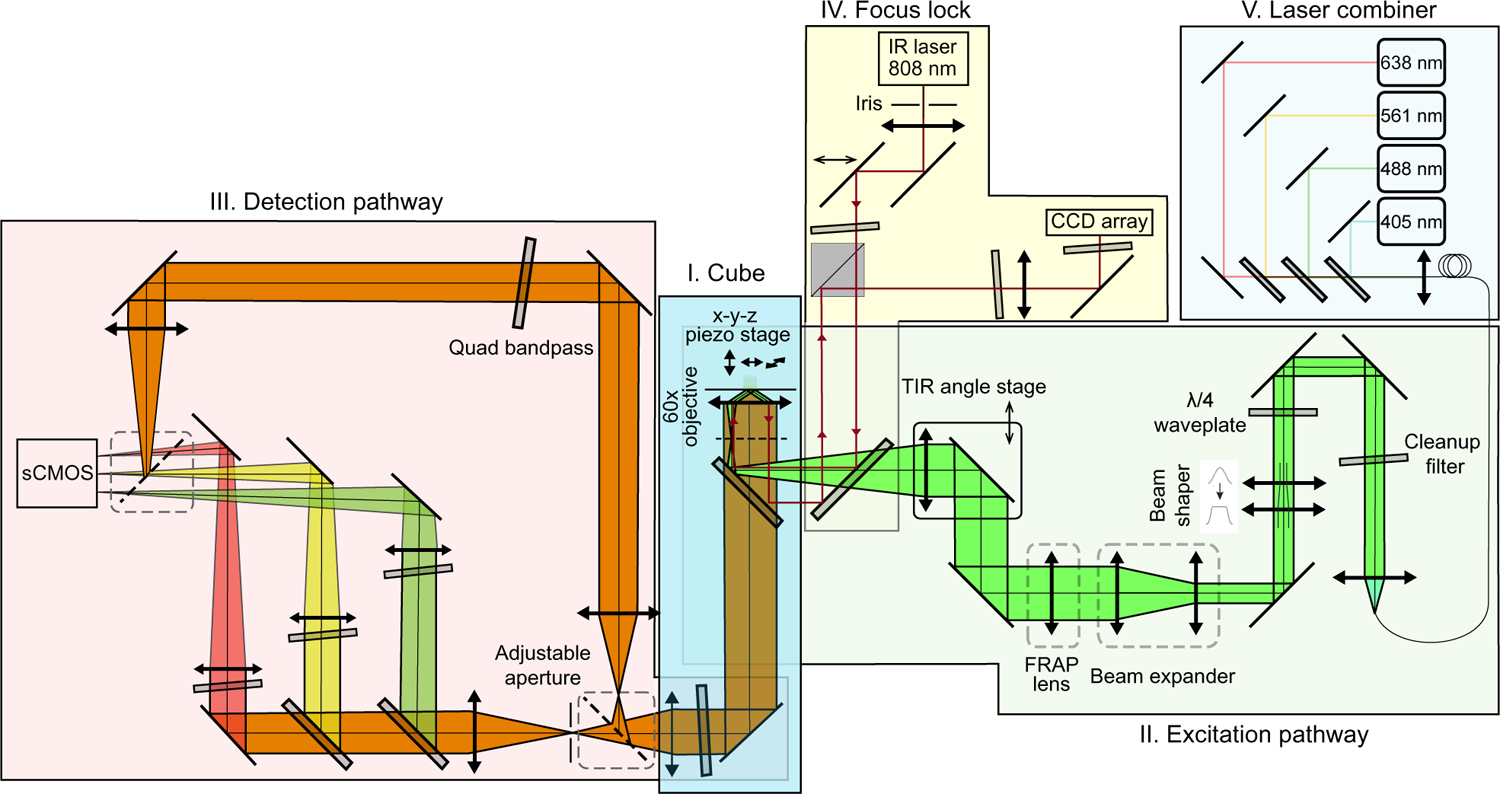
Optical pathway of the K2 microscope: I) The central cube houses the sample stage, objective, dichroic mirror for splitting excitation and emission light, and the tube lens. II) The excitation pathway launches a four-color laser beam with circular polarization and flat-top beam shape into the central cube. The angle of incidence is controlled using a motorized stage to switch between epifluorescence, HILO- and TIRF-imaging. A flip-in lens can be used to focus the beam at the sample plane for bleaching experiments. III) The detection pathway features a triple-color image splitter, projecting the field-of-view onto different regions of the sCMOS camera chip for simultaneous triple-color imaging. Two mirrors on magnetic mounts can be used to bypass the triple-color image splitter. IV) The focus stabilization pathway uses an infrared laser to detect and compensate for axial drift of the sample. V) A pre-assembled laser combiner box delivers the excitation laser beam via a single-mode fiber to the setup.

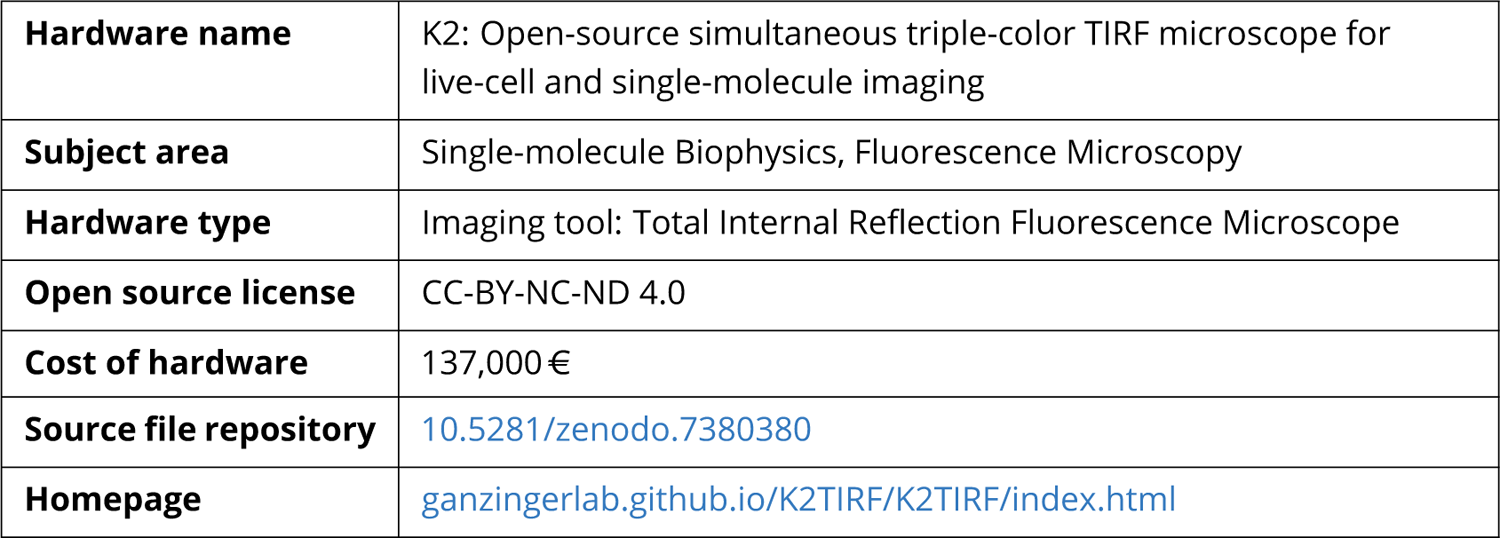

### Microscope body

The K2 single-molecule TIRF microscope is a custom-designed instrument that does not use a conventional microscope body (see ***Figure 3***). The central part is a solid, CNC-milled aluminum block, based on the miCube project [3]. The cube houses the objective, sample stage, dichroic mirror and tube lens. Unlike traditional commercial microscopes, the K2 does not include an objective turret or ocular, as they are not required for our experiments. This design results in a smaller footprint and excellent mechanical stability.

**Figure 3.**
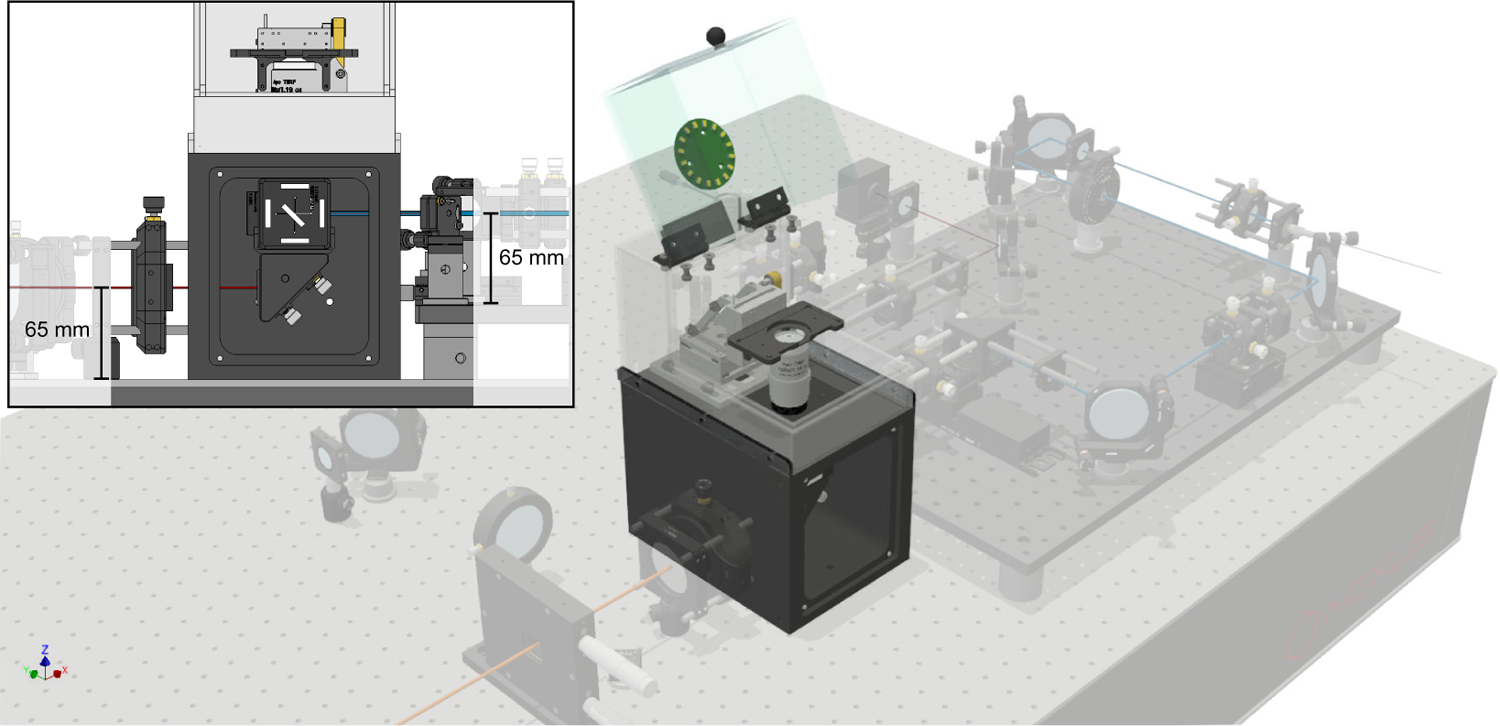
Central cube of the K2 microscope, showing the TIRF objective mounted on a thermal spacer, the sample stage with sample holder, and the dichroic mirror for splitting emitted fluorescence from excitation laser light inside the cube. The plexiglass cover encloses the objective and sample for added thermal stability and protection, and also provides a brightfield light source. Inset: The placement of the detection port on the central cube determines a beam height of 65 mm. The beam height of the excitation path is matched by placing the optical elements on an elevated breadboard, allowing the user to interchange alignment tools and pillar posts between the excitation and detection pathways.

A high numerical aperture oil-immersion objective (NA=1.49, CFI Apochromat TIRF, Nikon) with 60x magnification is mounted on a polyoxymethylene spacer to provide thermal isolation from the cube. A polyimide flexible heater, wrapped around the objective with Kapton foil, and a PT100 temperature sensor glued in place, enable precise temperature control via a proportional-integral-derivative (PID) controller (E5CC, Omron). This design is similar to other custom-made and commercial designs [15, 16, 21], and allows for active heating of the sample, which is thermally coupled to the objective via immersion oil. This enables live-cell imaging and *in vitro* experiments at physiological temperatures.

The sample stage of the K2 single-molecule TIRF microscope is equipped with a three-axis piezo stick-slip positioner (SLS-5252, Smaract) and a two-piece aluminum sample holder that accommodates square (18 mm to 26 mm side length), circular (25 mm to 37 mm diameter) and rectangular (62 mm to 80 mm length, up to 35 mm width) samples. Samples are held in place with magnets or an aluminum ring to minimize sample drift. The stage is controlled via the computer or a handheld joystick controller (see ***Figure 1***) and accurately positions samples with nanometer resolution and 30 mm travel range in all directions, allowing for high-throughput measurements of multi-well coverslides.

The excitation laser beam enters the central cube through a port on the right-hand side and is reflected upwards towards the objective by a dichroic mirror. Residual transmitted laser light is absorbed by a neutral density filter mounted on the dichroic mirror cube. Emitted fluorescence collected by the objective is transmitted through the dichroic mirror and reflected via a mirror towards the exit port on the left-hand side of the cube. A four-bandpass emission filter on the mirror mount blocks remaining laser excitation light from reaching the detection pathway. The dichroic mirror and the right-angle mirror are accessible through a door on the front of the cube, which is held in place with magnets.

The central cube is enclosed by a plexiglass cover that protects the sample and objective from dust buildup and improves temperature stability by minimizing convective air transport. A sensor for temperature and humidity is placed inside the plexiglass box and records the environmental parameters during experiments. The lid of the plexiglass cover includes a LED ring for brightfield imaging of cellular targets. For laser safety considerations, potential adopters of our design may consider using a non-transparent cover.

### Excitation pathway

The excitation pathway is mounted on an elevated breadboard (see ***Figure 4***), which helps to keep the length of posts short for stability. Spacers (39.7 mm height) were used to raise the breadboard (12.7 mm thickness) to the required level, resulting in final beam heights of 65 mm in both the excitation pathway and the detection pathway. In principle, the beam can be brought up to the required height also by using long enough posts or a periscope-assembly.

**Figure 4.**
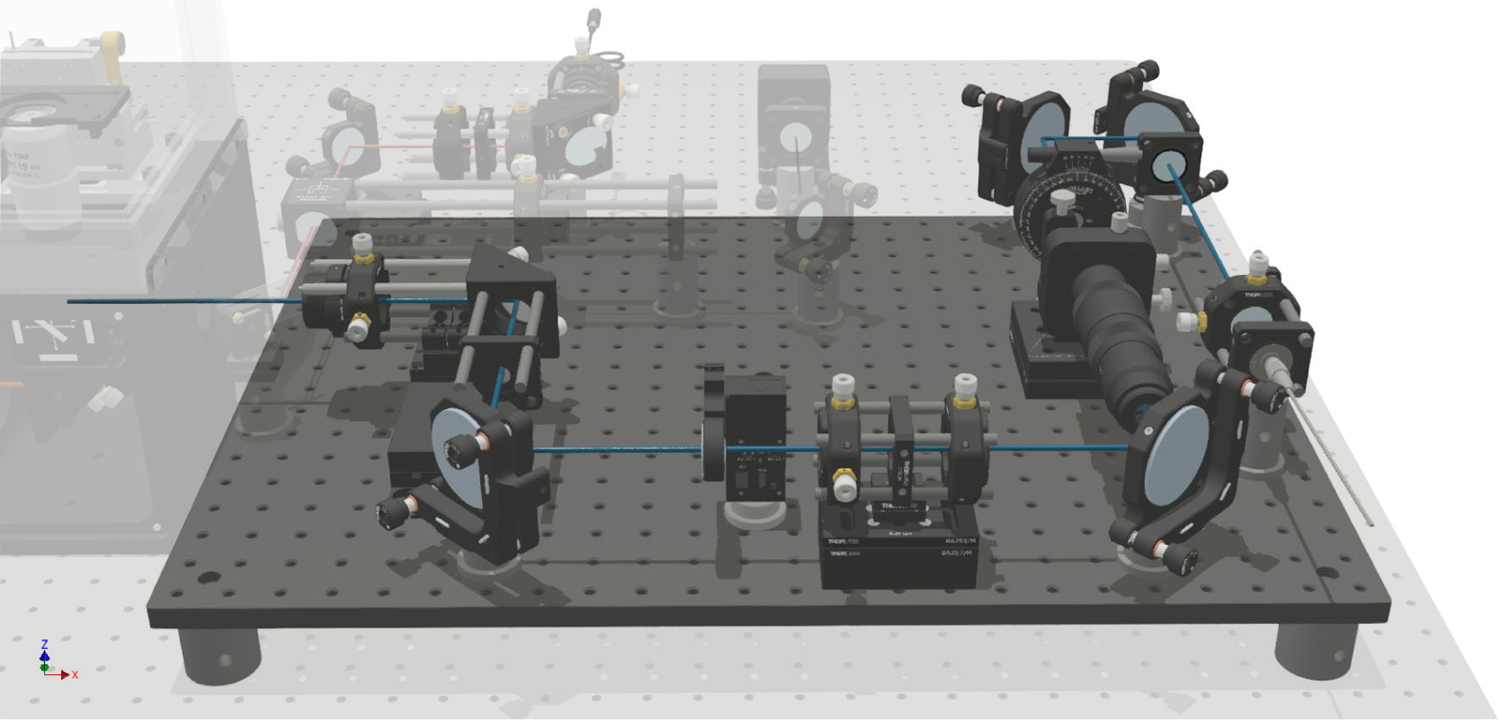
Excitation pathway of the TIRF microscope, featuring (in the order of placement along the optical path) the fiber collimator, laser clean-up filter, quarter-wavelength waveplate, refractive beamshaper, beam expander, flip-in FRAP lens and motorized stage with TIR focusing lens. These components work together to shape and deliver the excitation laser beam to the sample stage, enabling precise control of the angle of incidence for different imaging modes (*i.e.* epifluorescence, HILO, and TIRF). All parts are mounted on an optical breadboard to match the height of the central cube’s excitation port.

We use a commercial pre-assembled laser combiner with a single-mode fiber output for delivering four excitation wavelengths (C-FLEX laser combiner, Hübner Photonics; 405 nm 140 mW, 488 nm 200 mW, 561 nm 220 mW, 638 nm 195 mW) to the excitation pathway. Laser excitation is controlled by a custom-made laser trigger box, that allows alternating and simultaneous excitation synchronized to the camera exposure. Other approaches to laser triggering include the open-source NicoLase and SMILE projects [14, 22], as well as the commercial Triggerscope [23].

The TEM00 laser beam is re-collimated after exiting the fiber to a 1∕e^2^ beam diameter of 6 mm (@ 561 nm), directed through a quad-line bandpass for spectral clean-up, and an achromatic quarter-wavelength waveplate to ensure circular polarisation. The Gaussian laser beam profile is transformed into a flat-top profile using a refractive beam shaping device (piShaper 6_6_VIS, AdlOptica, comparable alternative: TopShape TSM25-10-D-B-6, asphericon) [17–19]. A (removable) telescope magnifies the laser beam by a factor of 2.5, resulting in a beam diameter of 15 mm. The flat-top profile of the laser beam begins to distort as it propagates over longer distances, so we designed the setup to keep the distance between the beam shaper and the objective as short as possible.

For FRAP experiments, an additional lens on a motorized flip-mount can be inserted into the optical path after the telescope, focusing the excitation laser into a 30 µm diameter spot in the imaging plane.

To enable switching between imaging modes (epifluorescence, HILO- and TIRF-imaging, see ***Figure 12***d), the lens focusing the laser beam onto the back focal plane of the objective is mounted on a motorized stage that allows the beam to be translated off-axis while maintaining its parallel alignment with the objective’s optical axis.

The lens that focuses the excitation beam onto the back focal plane of the objective is placed as close to the back focal plane as possible, which allows using a lens with a shorter focal length and therefore a bigger illuminated area at the sample plane. Using a telescope with a higher magnification to achieve an equally-sized illuminated area would result in clipping the laser beam, unless larger optics are used throughout the excitation path. To provide more flexibility in the arrangement of the optical elements, future versions of the central cube could be designed with a reduced distance between the objective port and excitation entry port. With the current design, our setup allows for a maximum homogeneously illuminated diameter of 190 µm.

### Detection pathway

The detection pathway is designed under the assumption that the microscope will mostly be used in a triple-color simultaneous imaging mode (see ***Figure 5***). Using a single camera to capture three color channels simultaneously is more cost-effective than using three separate cameras, and combining a 60x objective without additional magnification with a 2048×2048 px sCMOS chip allows us to achieve large field-of-views (73.65 µm x 73.65 µm per color channel) nevertheless. To achieve even larger field-of-views, the triple-color detection pathway can be bypassed by placing two mirrors on pre-installed magnetic mounts, resulting in a circular field of view of 190 µm diameter. A commercial alternative for image splitting is the OptoSplit (Cairn Research [24]).

**Figure 5.**
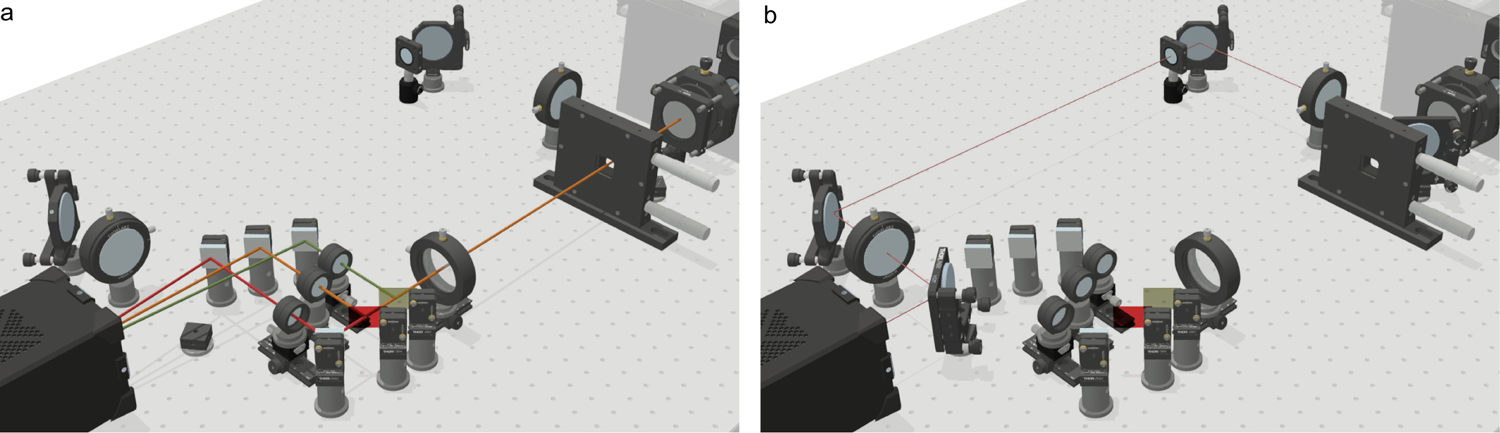
Detection pathway options of the K2 TIRF. **Left:** triple-color detection pathway with adjustable slit aperture and image splitter for simultaneous imaging of up to three color channels. **Right:** single-color full field-of-view path.

As described in the section about the central cube, fluorescence emission from the sample is collected by the objective, passes the quad-bandpass dichroic mirror in the cube and is reflected towards the detection pathway. A quad-line filter blocks residual excitation light from reaching the detection pathway. Emission fluorescence is directed through an infinity-corrected tube lens matching the objective, without further magnification (TTL200-A, Thorlabs). With the camera’s physical pixel size of 6.5 µm, this results in an effective pixel size of 6.5 µm∕60 = 108.3 nm.

In the triple-color imaging mode, the image of the fluorescent sample is cropped horizontally by a custom razorblade slit aperture in an intermediate image plane. This is necessary to avoid overlapping of the color channels when imaging them side-by-side on the camera chip (see ***Figure S29***). Splitting the fluorescence emission into the different spectral channels and filtering them is done in infinity space in a 4f-configuration. This reduces aberrations and allows more mechanical freedom, since the distance of the second to the first 4f-lens is not critical.

The first 4f-lens is positioned to have its focal plane aligned with the image plane formed by the tube lens. Then, the fluorescence emission is split into three colors by two low-pass dichroic mirrors, matched to the four-bandpass dichroic mirror in the cube, and one dielectric mirror. Furthermore, appropriate bandpass filters are placed in each color channel. Individual lenses mounted on rails and dovetail stages project the images of each color channel next to each other on the camera. The use of individual lenses allows precise co-alignment of the image planes and the correction of residual chromatic aberrations. A dielectric mirror is used in each color channel to project the spectrally-separated images onto the camera chip. Finally, three equally-sized square field-of-views are created by cropping the camera chip horizontally. By cropping in line with the scanning of the sCMOS chip, we are able to reduce the read-out time from 22.6 ms (full chip, 2048×2048) to 7.6 ms (central region, 2048×682). At a typical framerate of 25 fps, this approximately doubles the exposure time per frame (32.4 ms∕17.1 ms ≈ 1.89).

The optical path for the image splitter was designed to be as compact as possible to minimize vignetting, while at the same time allowing component accessibility for alignment. Additionally, projecting the image onto the camera at an angle introduces aberrations, therefore the individual focusing lenses and the mirrors are placed as close to each other as possible. As a consequence, this maximizes the distance between the last mirror and the camera and hence decreases the angles at which images are projected from the left and right color channel.

In the single-color imaging mode, the full 190 µm diameter illuminated field-of-view is imaged with spatially unseparated color channels. This is accomplished by bypassing the slit aperture and the image splitting pathway using two mirrors, that are placed manually on kinematic magnetic mounts. The first mirror is placed after the tube lens and the second mirror before the camera. Two lenses relay the image in a 4f-system, which conveniently also allows the placement of additional components like filters or phase masks in the fourier plane (see ***Figure 5***, ***Figure S28***).

### Focus stabilization pathway

In microscopy in general, but specifically in super-resolution and single-molecule experiments and timelapses, sample drift can significantly degrade the quality of the recorded data [25]. In live-cell and single-molecule tracking experiments, lateral drift of the sample usually does not interfere with the experiments as the typical drifting appears on much slower timescales than the molecular movement apparent in the sample (lateral sample drift in our setup: ≈ 1 nm min^−1^, diffusion of individual receptor proteins in cell membranes: ≈ 0.1 µm^2^ s^−1^ to 1 µm^2^ s^−1^).

Movement of the sample along the optical axis (axial drift), however, leads to a shift of the focal plane. Since in TIRF microscopy, fluorescence is only emitted from a thin layer close to the coverslip, this shift leads to a decreased signal-to-background ratio due to the defocus blurring of the point spread function, or even the loss of the object that the user initially focused on. Therefore, keeping the sample in focus is crucial, especially when performing long measurements and timelapses, or repeated acquisitions on different areas on a sample.

One way to measure and control the distance of the sample from the objective is to use the back-reflection of the excitation laser, with the drawback that measuring and updating the *Z*-position can only be done during the excitation cycle [8]. A dedicated infrared laser is a more versatile option, as it does not excite commonly used fluorescent proteins and dyes, allowing for continuous measurement and control of the sample’s axial position without interfering with fluorescence measurements.

The principle of the focus stabilization is depicted in ***Figure 6***a. First, the sample is brought into focus manually and the position of the back-reflected beam is stored. Axial drift Δ*Z* of the sample leads to a lateral displacement Δ*d* of the totally internally reflected infrared laser beam. The back-reflected laser beam is relayed through additional optics, resulting in a displacement Δ*d*^′’^, that is determined by comparing the center peaks of the previously saved reference and the currently measured laser profiles on a line-array sensor. To mechanically compensate for the axial drift, an active control mechanism reacts to the lateral displacement and moves the sample stage to keep the position of the back-reflected beam stable.

**Figure 6.**
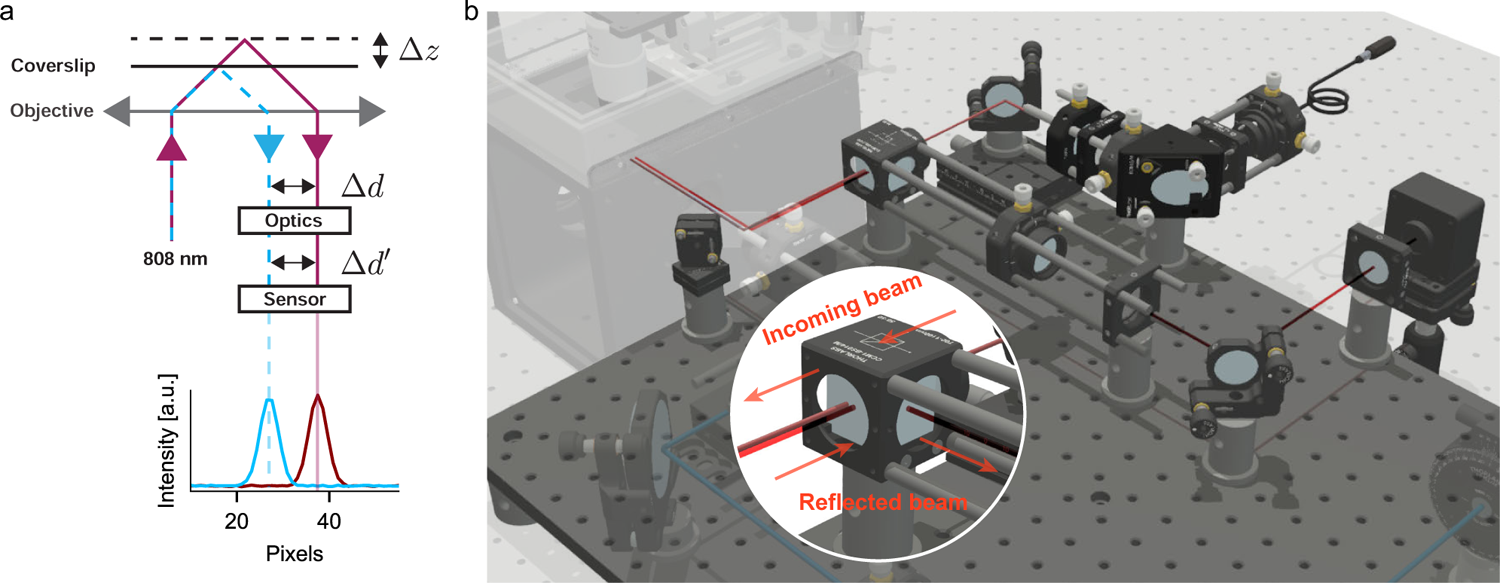
**a**) Schematic of the qgFocus lock: an infrared laser beam is focused off-center at the back focal plane of the objective and is totally internally reflected at a coverslip. Upon an axial movement Δ*Z* of the coverslip, the reflection of the beam is shifted laterally by a distance Δ*d*, which is translated into a distance Δ*d*^′^ through downstream optics. A sensor registers the intensity profile of the back reflected beam and determines its center by a Gaussian fit. A control loop acting on the sample stage moves the sample such that the displacement of the currently measured intensity profile and of the referenced intensity profile is approaching zero. **b**) Render of the focus stabilization pathway of the K2 TIRF, mounted alongside the excitation optics on an elevated breadboard. Inset: Separation of incoming infrared laser beam and back-reflected beam by a beamsplitter cube.

Our focus lock implementation (see ***Figure 6***b), qgFocus (short for **q**uite **g**ood Focus), is similar to other custom-built and commercial designs such as pgFocus [10] or CRISP [9]: A collimated infrared laser is coupled into the excitation path by a short pass dichroic mirror and attenuated with a neutral density filter so that it does not interfere with the fluorescence signals. A manual micrometer stage is used to shift the infrared laser beam off-axis, bringing it into total internal reflection. The infrared laser is then focused at the back focal plane, reflected at the glass-sample interface, and travels back towards a beamsplitter. The back-reflected laser beam is diverted by the beamsplitter towards a bandpass filter and finally focused onto a linescan sensor (TSL1401, Parallax). The intensity curve is registered and fitted with a Gaussian function. A control loop that locks onto the saved in-focus position of the Gaussian function center then moves the piezo-driven sample stage in the axial direction to maintain focus. In our system, manual sample travel in *x-y* during focus-controlled experiments is limited to 100 µm s^−1^ to avoid jumps in the back-reflected beam position that cannot be followed by the control loop.

### Scripts

Our microscope control software includes a script interface that allows users to control the setup and perform various tasks using C++ commands. Examples of scripts that can be run using the interface are available at https://github.com/GanzingerLab/K2TIRF/tree/master/K2TIRF/scripts. The available examples include a simple *x-y* raster scanning script, a laser power calibration method that we used to calibrate the laser percentage settings (see ***Figure S50***) and a script that continuously moves the sample stage up and down, which we used during the alignment process (see ***Figure S26*** and ***Figure SM1***).

### Design files

A virtual “copy” of the setup with all optical components has been uploaded to the public repository Zenodo (10.5281/zenodo.7380380). Please note that while we tried to reproduce the physical positions and distances of the optical components as faithfully as possible, this CAD model mainly serves illustration purposes and should not be used as an alignment guide.

For all custom-made mechanical components, CAD files and technical drawings are provided, following best practices for sharing reproducible microscope hardware [26]. For all custom-made electronic components, PCB layouts and electronic design files are provided. A ready-to-view model is uploaded to Autodesk Drive (https://autode.sk/3vdsOu0). The source code of the software for the operation of the setup is uploaded to the Zenodo repository. Python code for processing and analyzing single-molecule imaging data is available at the Ganzinger group Github (https://github.com/ GanzingerLab)

### Design Files Summary

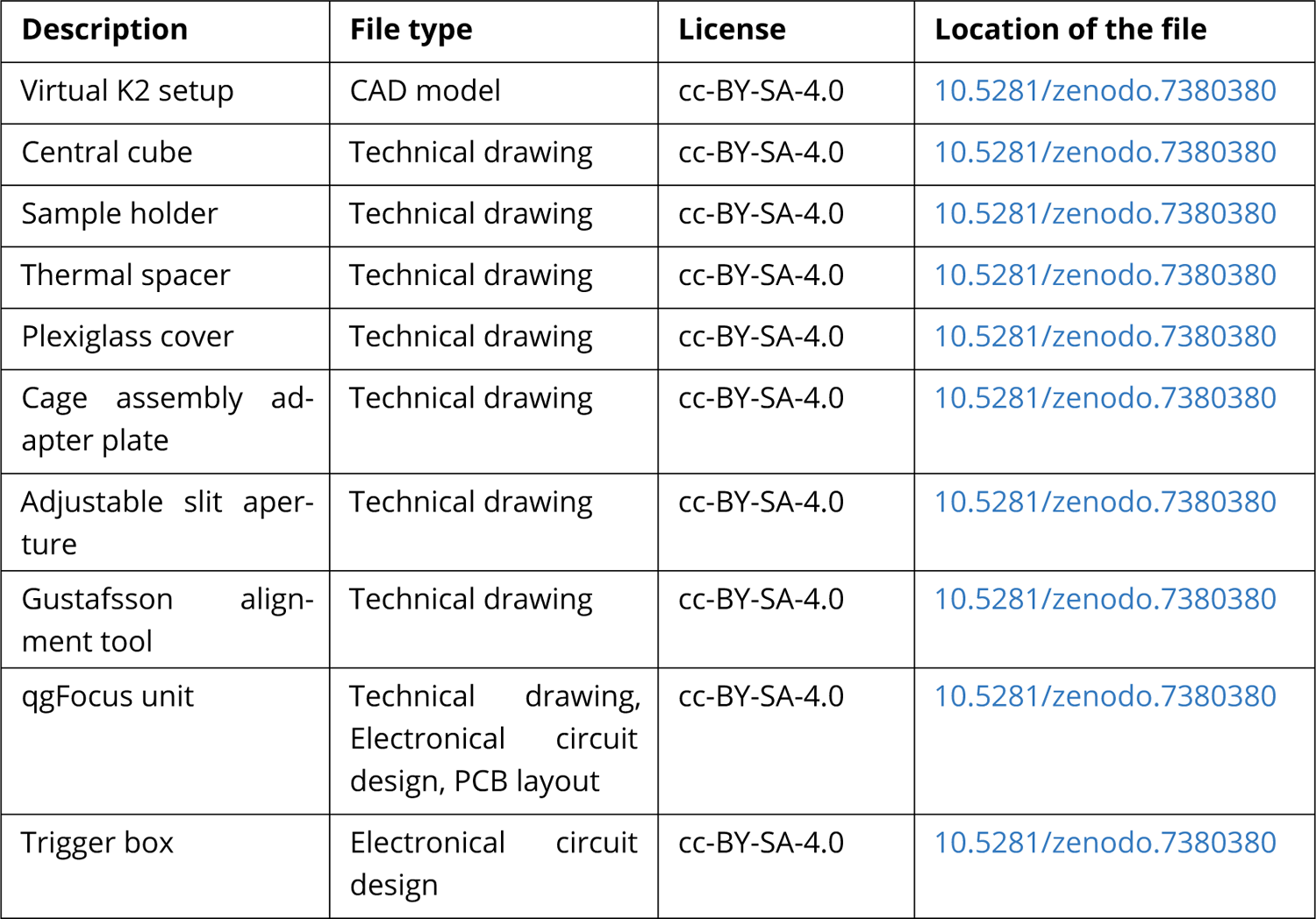

| Description | File type | License | Location of the file |
| --- | --- | --- | --- |
| Virtual K2 setup | CAD model | cc-BY-SA-4.0 | [10.5281/zenodo.7380380](https://zenodo.org/record/7380380/files/Virtual_K2_setup.zip) |
| Central cube | Technical drawing | cc-BY-SA-4.0 | [10.5281/zenodo.7380380](https://zenodo.org/record/7380380/files/Central_cube.dwg) |
| Sample holder | Technical drawing | cc-BY-SA-4.0 | [10.5281/zenodo.7380380](https://zenodo.org/record/7380380/files/Sample_holder.dwg) |
| Thermal spacer | Technical drawing | cc-BY-SA-4.0 | [10.5281/zenodo.7380380](https://zenodo.org/record/7380380/files/Thermal_spacer.dwg) |
| Plexiglass cover | Technical drawing | cc-BY-SA-4.0 | [10.5281/zenodo.7380380](https://zenodo.org/record/7380380/files/Plexiglass_cover.dwg) |
| Cage assembly adapter plate | Technical drawing | cc-BY-SA-4.0 | [10.5281/zenodo.7380380](https://zenodo.org/record/7380380/files/Cage_assembly_adapter_plate.dwg) |
| Adjustable slit aperture | Technical drawing | cc-BY-SA-4.0 | [10.5281/zenodo.7380380](https://zenodo.org/record/7380380/files/Adjustable_slit_aperture.dwg) |
| Gustafsson alignment tool | Technical drawing | cc-BY-SA-4.0 | [10.5281/zenodo.7380380](https://zenodo.org/record/7380380/files/Gustafsson_alignment_tool.dwg) |
| qgFocus unit | Technical drawing, Electronical circuit design, PCB layout | cc-BY-SA-4.0 | [10.5281/zenodo.7380380](https://zenodo.org/record/7380380/files/qgFocus_unit.zip) |
| Trigger box | Electronical circuit design | cc-BY-SA-4.0 | [10.5281/zenodo.7380380](https://zenodo.org/record/7380380/files/Trigger_box.zip) |

#### Bill of materials

The up-to-date bill of materials for the K2 TIRF project is available on the open-access Github page (https://ganzingerlab.github.io/K2TIRF/K2TIRF/component_table.html), providing detailed information on the components and materials needed to build the microscope.

#### Build instructions

For an overview of the layout of the optical components, see ***Figure 1*** (CAD-model render), ***Figure 2*** (schematic of the optical pathway) and ***Figure 7*** (picture of the setup). The setup requires approximately 190 cm by 90 cm of optical bench space, as well as a similar area (*e.g.* above the optical bench) for the laser combiner, objective heating PID controller and laser trigger box, as well as space for the computer and screens.

**Figure 7.**
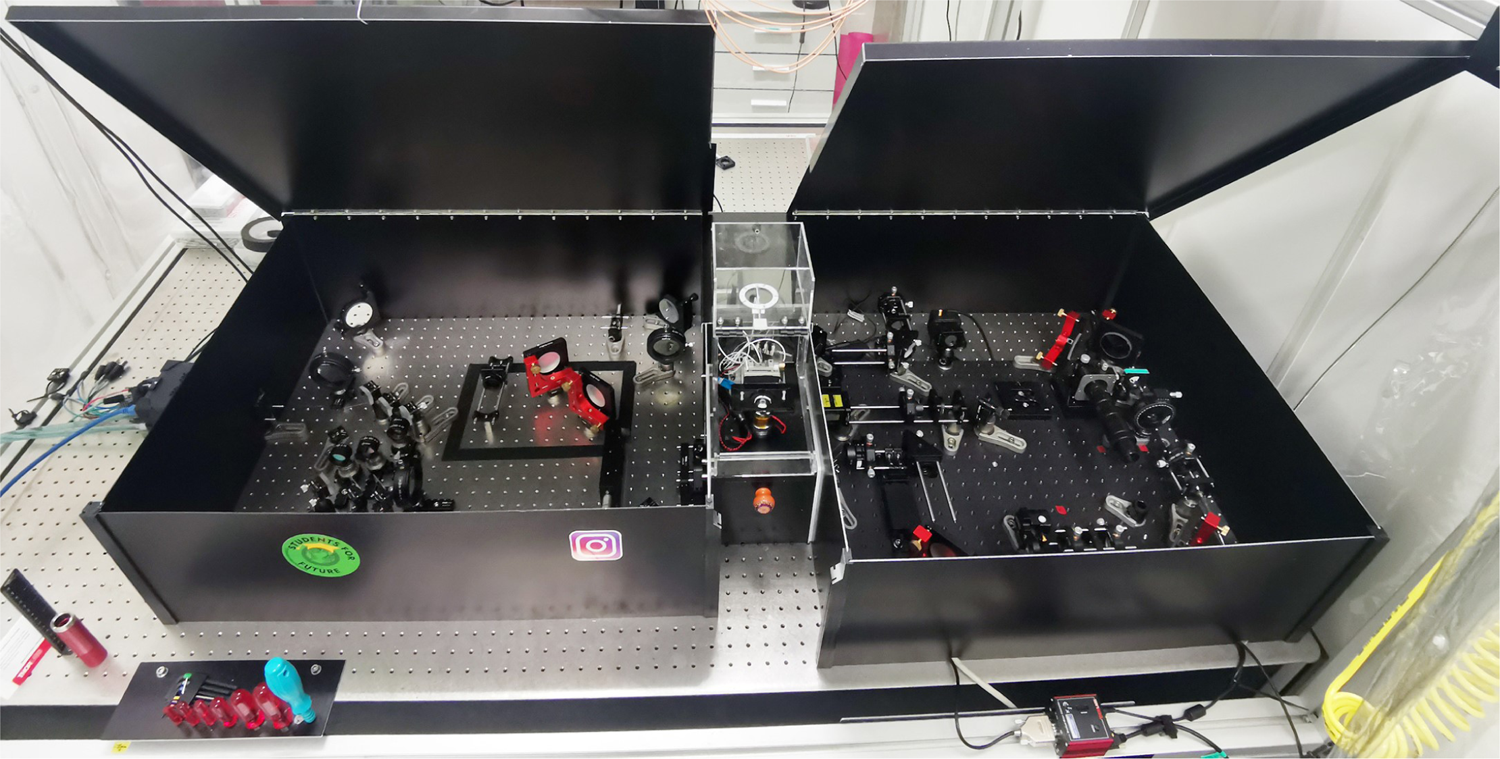
Picture of the K2 TIRF microscope. From right to left, there are the excitation and focus lock pathways on an elevated breadboard, the central cube with the objective, sample stage and a plexiglass cover, and the triple- and single-color detection pathways with the camera on the left. The excitation and detection pathways are enclosed in light-tight aluminium boxes to reduce background light levels, minimize laser safety issues, and to prevent dust from building up.

When setting up the microscope, the most convenient approach is to start with building and aligning the excitation pathway and use the laser source for alignment. However, if this step needs to be deferred and another laser source is available (*f.i.* the Gustafsson alignment tool), it is possible to begin with the central cube and the detection pathway, since the excitation and detection pathways can largely be assembled and aligned independently of each other. In addition to commonly used alignment tools such as viewing disks, reticles, irises, and calibration slides, we use a cage system mounted on the objective port to simplify the alignment of the excitation path, and the Gustafsson alignment tool for the detection path alignment [27]. The Gustafsson alignment tool is a small diode laser aligned in a mount that is screwed into the objective thread. The wavelength should be situated within one of the fluorescence emission bands, *e.g.* in the GFP-like emission band between 500 nm and 550 nm. This allows the alignment laser to pass through the emission dichroic mirror, and, if alignment is performed in the correct order, the laser beam can be used to align the entire detection pathway. General approaches and recommendations for the alignment of optical parts are described in [28].

Detailed, step-by-step build instructions are provided in the supplementary information.

### Operation instructions

#### K2 software

A software package for operating the setup was written in C#. It features a graphical user interface (GUI) that provides a simple and intuitive way to control and customize the microscope setup for different experiments. The GUI includes a number of panels for changing settings and directly controlling the various components of the microscope, such as the camera, stage, lasers, and focus lock. ***Figure 8*** and ***Figure 9*** show some of the most important panels for controlling these components. The software also allows users to save experiment routines as templates, with a wide range of custom settings and options.

**Figure 8.**
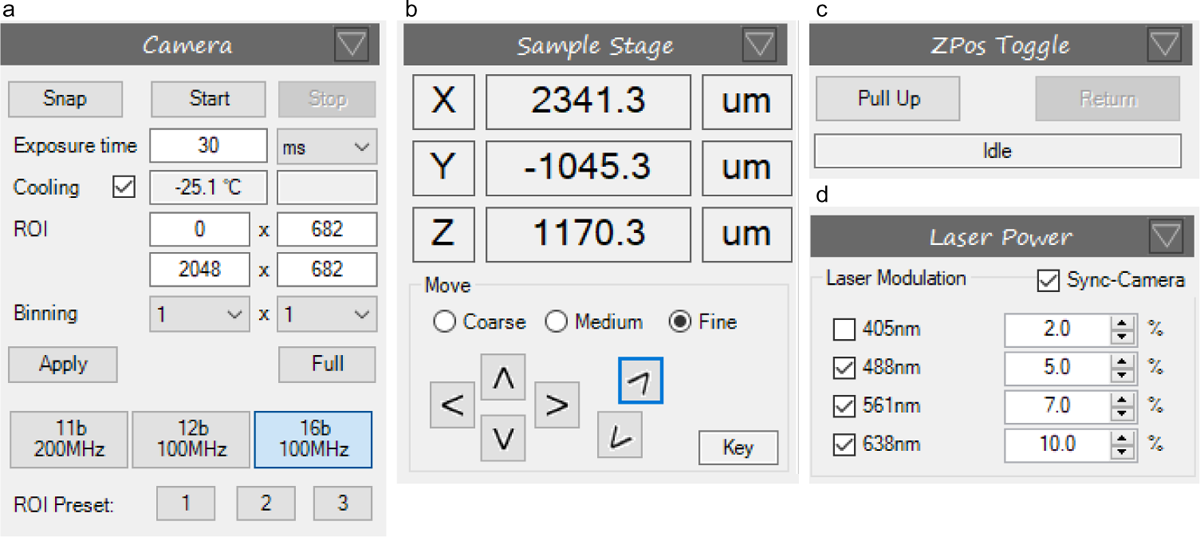
Main control panels of the K2 software. **a**) Camera: In this panel, the exposure time, manual/preset regions of interest (*ROI*), binning and bit-depth of the camera are configured, as well as live-viewing of the camera image (Start/Stop) and snapping a single frame (Snap). **b**) Sample Stage: Piezo stage position readings, and arrow keys (operated via cursor click or with arrow keys on keyboard), complementary to the physical joystick, with three user-defined step sizes. **c**) Zpos Toggle: Toggle for pulling up the sample away from the objective, and returning the sample to the initial *Z*position (*Pull Up*, *Return*). **d**) Laser powers: Laser power settings used during camera live-view, calibrated to achieve a linear percent-to-power relation. The synchronization of the lasers to the camera trigger can be switched off.

**Figure 9.**
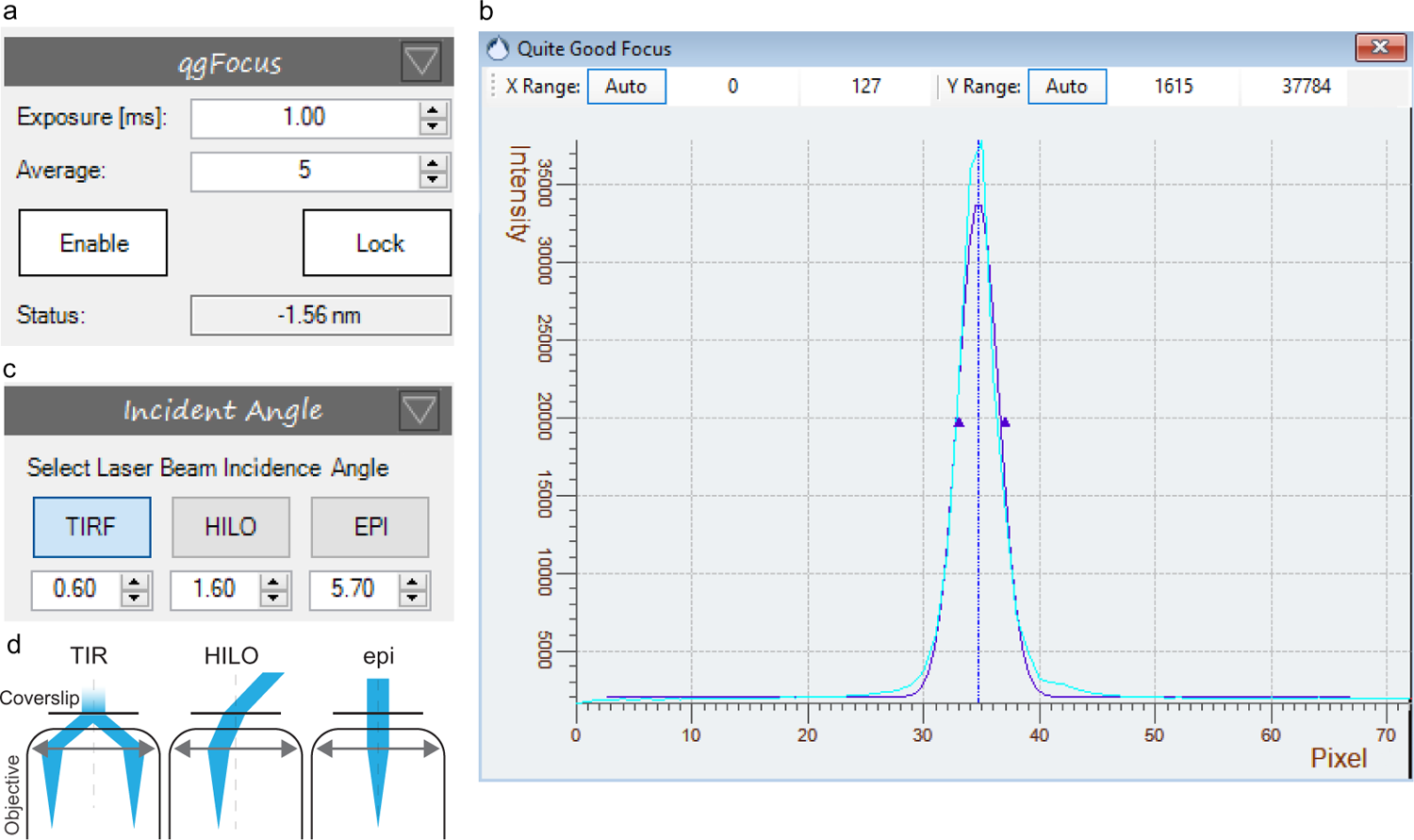
**a**) qgFocus control panel to switch on the focus lock sensor (*Enable*), to lock onto a focus position (*Lock*) and to set the exposure time and number of averaging cycles. The *Status* reports on the current distance of the stage with respect to its initial *Z*-position. **b**) qgFocus sensor output displaying the measured back-reflected laser intensity profile on qgFocus sensor (cyan line) and a Gaussian fit to the intensity profile (blue line). **c-d**) TIR stage panel: Stage positions for TIRF, HILO and epifluorescence imaging are set and can be called during the experiment to change the angle of incidence of the excitation laser beam.

**Figure 10.**
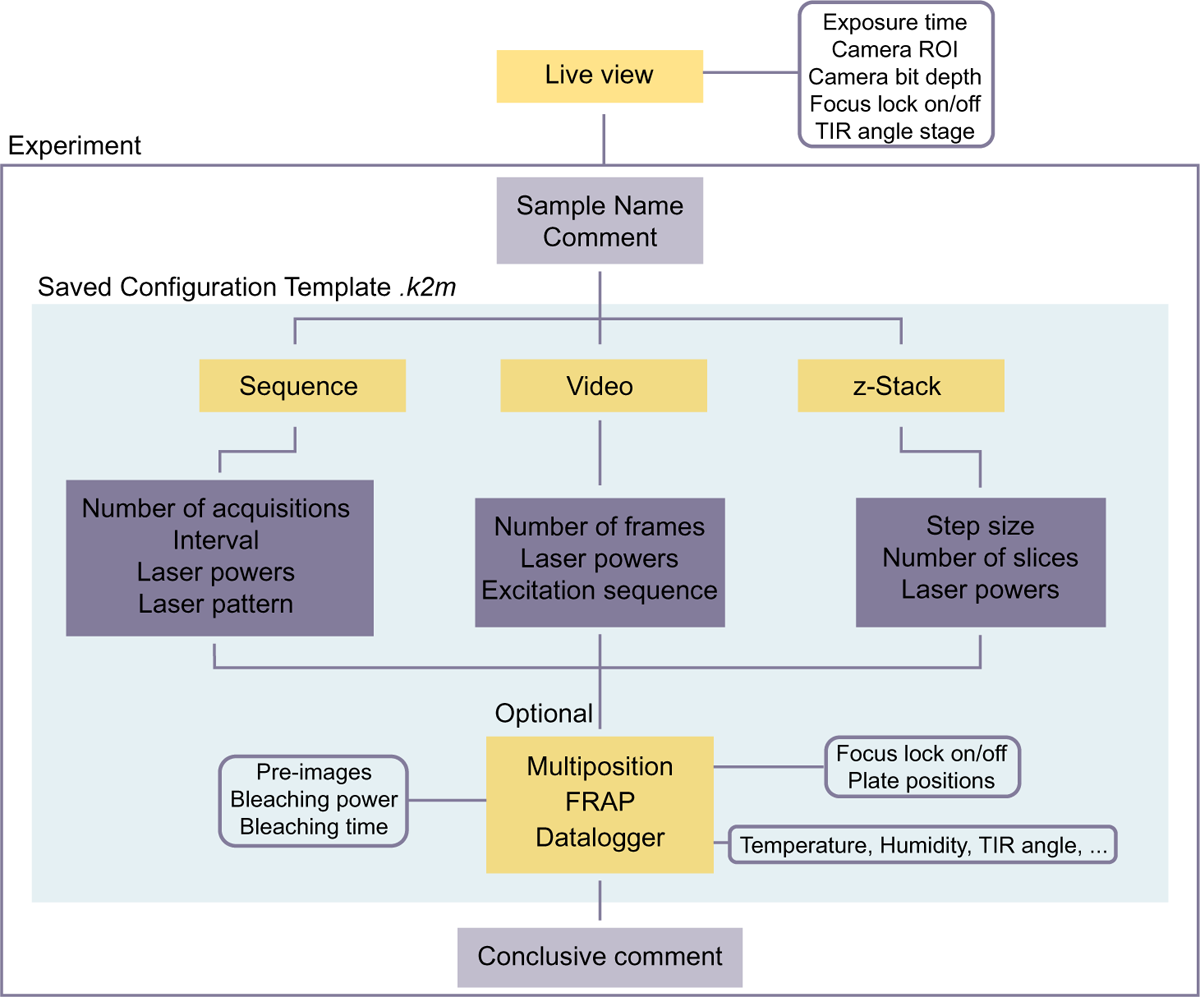
Experiments configuration diagram: Experimental settings are optimized during live-view. An acquisition is then performed by configuring, or reusing a experiment configuration template and adding details about the sample and experimental conditions. For the acquisition itself, three options (*Sequence*, *Video* and *z-Stack*) are available and for each option, the necessary settings are configured. Optionally, the experiment can be performed at different sample positions (*Multiposition*), as a *FRAP* experiment, and with additional recording of environmental parameters. After the experiment is performed, a comment may be added to the file.

The source code has been uploaded to the public repository Zenodo (10.5281/zenodo.7380380), excluding any third party software (which we cannot distribute) and excluding internal libraries which are shared among other projects in the institute. Thus, this release will not compile into the final product but it can serve as an inspiration for other development efforts. For all commercial components (sample stage, camera, laser source, flip-in lens, TIR angle stage) LabView virtual instruments, python libraries, and/or plugins for micromanager [29] exist and can be used alternatively for the setup control.

In the following, the operation of the setup is described using the K2 software.

#### Camera

The camera panel (***Figure 8***a) provides access to all relevant camera settings, such as the exposure time, region of interest, pixel binning, and bit-depth. The Start/Stop buttons can be used to activate or stop the camera live-view streaming (Figure 15), and the Snap button takes a single frame snapshot.

#### Live-view and RGB viewing mode

The camera image is streamed in the live-view mode, allowing the user to position the sample and to search for a suitable region of interest, as well as to determine appropriate imaging conditions, such as laser power and exposure time. The contrast and colormap settings of the streamed image are set by the user (***Figure S46***). The K2 software also features a RGB viewing mode, that super-positions the triple-color separated camera channels as a red-green-blue overlay in the live-view (***Figure S47***). This allows to quickly assess colocalization in different channels and therefore speeds up screening for an interesting cell or region of interest on the sample, and is a convenient tool during the alignment of the triple-color image splitter (see ***Figure S26***).

#### Sample stage

The piezo stick-slip stage is steered both with the software (either by clicking on the sample stage panel arrows, or using arrow keys on the keyboard) and by a handheld joystick controller. To ensure safe and efficient sample handling, the user can adjust the step sizes *Coarse*, *Medium* and *Fine* and define position limits (***Figure S48***).

A *Z*-position toggle is available that improves the experimental workflow and reduces the amount of immersion oil needed for high-throughput measurements. It works by moving the sample upwards away from the objective, with *x* and *y* positions unchanged. The sample is first moved slowly (at 0.8 mm s^−1^ for 3 mm), to avoid introducing air bubbles into the immersion oil and to give the fluid time to draw back to the objective (see ***Figure SM2***). The sample can then be exchanged or moved horizontally, and returns to the previous *Z*-position, again approaching the last millimeters towards the objective slowly. This considerably improves the handling when several samples are imaged and need to be exchanged often. When using multi-well samples, or when imaging many different regions on a single sample, the *Z*-position toggle is used for removing the sample from the immersion oil, repositioning in *x-y*, and moving it back to the original *Z*-position (see ***Figure SM3***). This reduces the amount of immersion oil that has to be reapplied and thus allows for high-throughput measurements with minimal user interaction. The exact values for the distance and speed limits are user-defined in the K2 software backend settings (***Figure S49***).

### Laser powers

The laser powers panel controls laser on/off state and excitation power, and features the option to deselect synchronization of the excitation to the camera exposure. A linear relation between the power percentage set by the user and the actually delivered laser power is established in an initial calibration step (see ***Figure S50***).

#### qgFocus

In the qgFocus panel, the focus lock sensor’s exposure time and number of averaging cycles are set (***Figure 9***). Within an axial range of about 5 µm around the samples’ in-focus position, the back-reflected infrared laser is registered as a sharp peak on the qgFocus sensor and fit with a Gaussian function. For locking the focus position, the center position of the Gaussian fit is saved and used as the setpoint for a control loop.

*Action point:* During the initial usage of the setup, determine the control loop variables to ensure that the focus position stabilizes within a few seconds after moving the sample in *x-y* (used values: proportional gain 0.7, integral time 2 s, integral limit 1 × 10^10^ s).

Once the focus lock is engaged, the qgFocus panel displays the extent to which the stage has been shifted to maintain the sample in focus. The sample stage movement is restricted to a speed of 100 µm s^−1^ to allow the qgFocus to follow the sample’s *Z*-position continuously. This speed limit can be defined by the user in the K2 software backend settings (***Figure S51***).

#### Angle of incidence

The motorized TIR stage sets the angle of incidence of the excitation laser beam, allowing the user to switch between epifluorescence, HILO and TIRF imaging modes by calling user-defined positions on the Angle of Incidence panel (see ***Figure 9***).

*Action point:* When operating the setup the first time, the TIR stage positions corresponding to the different imaging modes are determined with a fluorescent sample (*e.g.* dye in solution) mounted: Starting from the epifluorescence position after setup alignment, the TIR stage is initially moved 4.5 mm, and then in steps of 0.1 mm. The transition from epifluorescence to TIRF imaging is easy to recognize (see [30] for a detailed description), as interference patterns caused by specs of dust scattering the coherent excitation laser beam, become visible. In sparse single-molecule type of samples, these interference patterns are not visible, but the transition is marked by a sharp increase in the signal-to-background ratio [5]. HILO imaging, where the laser beam penetrates the sample at an angle instead of being totally internally reflected, is typically performed close to the TIR setting (*i.e.* between 4 mm to 5 mm from the epifluorescence position).

### Pre-configuring experiments

Imaging conditions and sequences may be pre-configured as saveable experiment configuration file (*\*.k2m*) and can be reused as templates. The user can choose from three general modes: a) Sequence acquisitions, where a pre-programmed laser excitation sequence is looped through every timepoint - for a user-defined number of times, allowing for both alternating laser excitation style experiments and simple timelapses. b) Video acquisitions at the maximum camera framerate, with one or several lasers simultaneously activated. c) *Z*-stacks, where images are taken at pre-defined axial positions, with one or several lasers switched on.

Each of these modes can be combined with a FRAP experiment at the beginning of the recording, or a multiposition acquisition. During a multiposition acquisition, the software performs the pre-configured acquisition at different, pre-defined stage positions.

The experiment configuration, sample name and optional comments are saved as a *\*.txt* file along with the microscopy data (*\*.raw* and *\*.yaml*.)

### Sequence acquisition

For certain experiments (*e.g.* FRET), fluorophores need to be (alternatingly) excited by a single excitation wavelength at a time. This can be done by performing a Sequence Acquisition (see ***Figure S55***). Any excitation sequence, with one or several lasers excited during a given camera frame, can be pre-programmed, and is looped through for every timepoint throughout the duration of the experiment. The interval between each timepoint can be set, which also allows for timelapse recordings (see ***Figure S56***).

### Video acquisition

The video acquisition mode allows the simultaneous imaging of multiple color channels, *e.g.* for multi-color single-molecule tracking experiments, at the maximum camera framerate without intervals between acquired frames. For every frame, the same pre-defined set of lasers at a certain power percentage will be used to excite the sample (see ***Figure S57***). Additionally, there is the option to use in-frame alternating laser excitation. For simultaneous imaging of blue excitable dyes (*e.g.* ATTO488) and cyanine dyes (*e.g.* ATTO643), this was demonstrated to decrease the bleaching of red dyes induced by the 488 nm laser drastically [31]. The duration of the pulses is set to be at least ten times smaller than the total frame exposure time, and the triggering scheme is such that the 488 nm laser is pulsing anti-cyclic with respect to the others (see ***Figure S58***).

### zStack

Although TIRF microscopy implies collecting fluorescence from the sample surface plane, most TIRF setups allow changing the angle of incidence to HILO or epifluorescence imaging. In these imaging modes, a *Z*-stack can deliver additional information along the axial dimension. The K2 software requires a starting *Z*-position, the number of slices and *Z*-step per slice and will record one frame per slice with a given laser power and exposure time. During the acquisition, the focus lock is disengaged and the sample is moved back to the original *Z*-position after the last slice.

### FRAP

Fluorescence recovery after photobleaching (FRAP) is a method for studying diffusion dynamics by bleaching the sample locally and observing the recovery of fluorescence by diffusive exchange of unbleached fluorophores into the bleached area. A FRAP experiment in the K2 setup is performed by collecting a number of pre-bleaching frames (either in Sequence or Video-mode), followed by a bleaching phase where the flip-in lens is flipped into the excitation path, the power of one or several lasers is increased. After the bleaching phase, the recovery is recorded using the Sequence- or Video-mode (see ***Figure SM4***). Between bleaching and the recovery phase, a short delay is introduced to account for the finite flipping time of the flip-in lens (flip-in: 0.5 s, flip-out: 1 s). The area and the position of the bleached spot is determined by the positioning and focal length of the flip-in lens.

### Multiposition acquisition

Coordinates (*x*,*y* and *Z*) of stage positions can be generated (*e.g.* for a raster scanning of the sample, see ***Figure S52***), saved in a list and called manually, or automated during an experiment where a pre-designed (sequence, video, and/or FRAP) acquisition is performed at each position of the coordinate list (see ***Figure SM3***). If the focus lock is engaged, the stored *Z*-position is ignored.

#### Starting up the setup

Before turning on the lasers, verify that any removable optical component (beamshaper, beam expander, mirrors for bypassing the triple-color pathway) and filters are placed correctly depending on the experimental requirements. Make sure that the flip-in lens is flipped out, the TIR stage is in a safe position, the objective correction collar is in the correct position, and that the objective was cleaned properly by the last user. If heating is required, make sure to turn it on about 30 min before the experiment, to allow the temperature to stabilize (see ***Figure 14***).

Switch on the required power supplies (lasers, camera, camera cooling, motorized stages and mounts, computer) and mount the sample. Open the setup control software and initialize the motorized stages. Switch on the appropriate lasers at a low power and start the qgFocus and the live-view of the camera. Bring the sample into focus using the sample stage joystick controller. As described before, during the initial operation, the qgFocus control loop settings and the TIR stage positions for epifluorescence, HILO and TIRF imaging should be determined to ensure optimal imaging performance.

### Validation and characterization

#### Multicolor channel alignment

Chromatic dispersion and misalignment of optical elements in the detection pathway lead to distortions across the different color channels, complicating the analysis and interpretation of multicolor microscopy data. Especially for (single-molecule) colocalization experiments, different color channels need to be precisely co-aligned. The physical alignment of the color-splitting optical path is done while imaging a calibration slide and superimposing the three color channels, as described in the supplementary build instructions (see ***Figure S26***). In doing so, we achieve an average spatial variation across different color channels below 1 µm. The remaining spatial offset between the channels is then corrected computationally: first, accumulated bead positions from raster scanning the calibration slide are assigned throughout the corresponding channels (see ***Figure 11***a-b). A spatial correction map is then calculated (***Figure 11***c) using a recently introduced method based on Zernike polynomial gradients [32]. Using this correction map, mean residual spatial variations are below 30 nm (***Figure 11***c-d).

**Figure 11.**
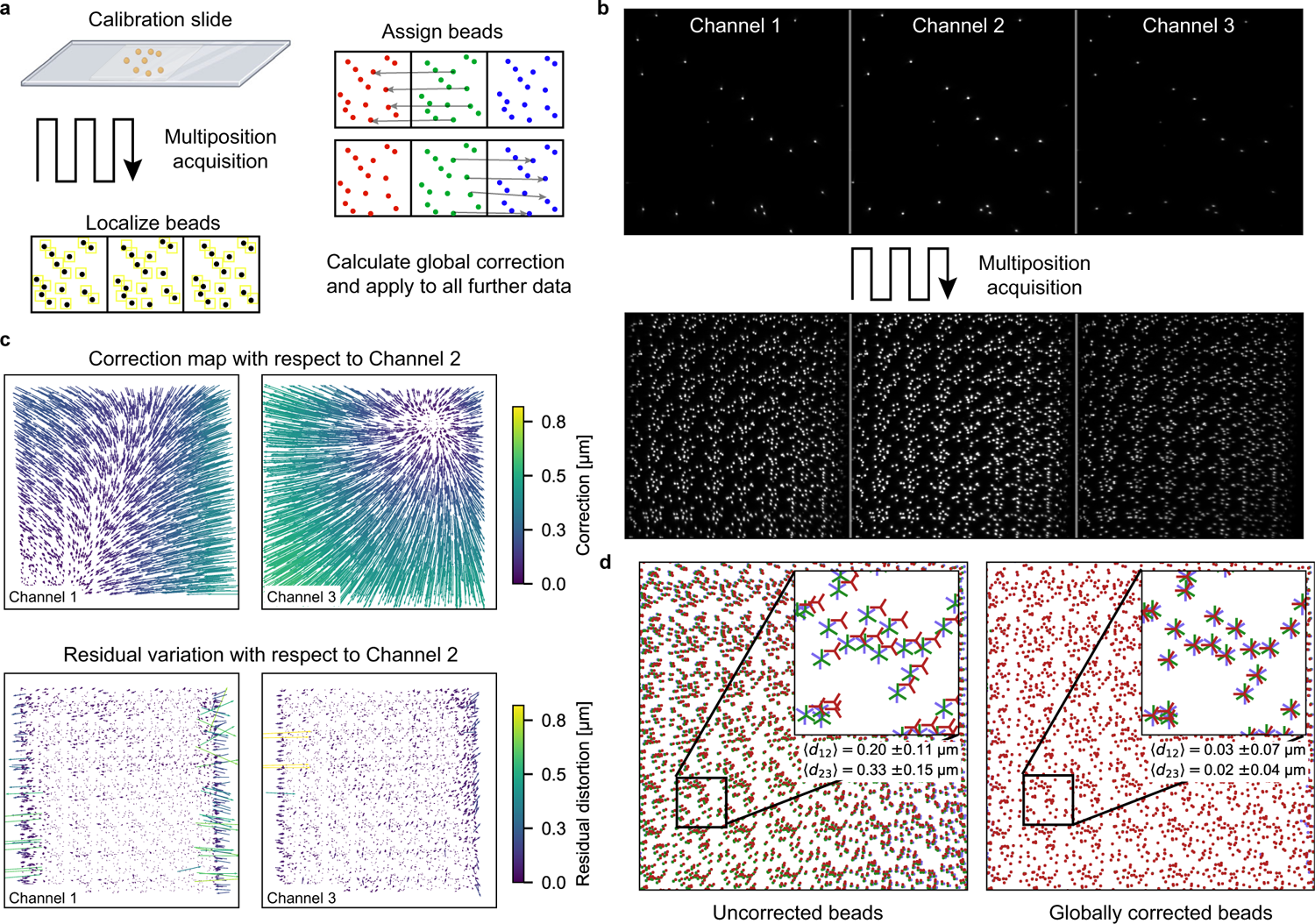
Multicolor channel alignment a)Multicolor-labeled Tetraspeck beads are deposited sparsely on a coverslide and imaged at multiple positions to collect a large number of *x* − *y* sample points. Identical beads are identified in the three color channels and localized frame-by-frame. Variations of bead *x*− *y* coordinates relative to the center channel are calculated pairwise, and used to compute a global correction map. This correction map is then used to correct single-molecule localizations acquired in further experiments. **b)** Upper row: Image of Tetraspeck beads in the three color channels, for a given sample position. Lower row: Maximum projection of all bead images acquired during a multiposition raster-scan. **c)** Upper row: Correction maps of left and right channel, with respect to the middle channel. Lower row: Residual spatial variation remaining after the correction is applied to the calibration data. Arrows indicate the direction of the necessary correction, with their length and color representing the amount of correction needed. **d)** Superimposed *x* − *y* positions of Tetraspeck beads with uncorrected coordinates (left) and corrected coordinates (right), with average spatial variations ⟨*d*_12_⟩ and ⟨*d*_23_⟩ of identical beads in the left and right channel, with respect to the middle channel.

#### Sample drift and focus stabilization

Minimizing sample drift is crucial in (TIRF) microscopy to ensure optimal signal-to-background ratios and the reproducibility of measurements. Therefore, we evaluated the axial and lateral drift of our K2 TIRF microscope, with and without the qgFocus focus stabilization, during a three hour experiment. Fluorescent beads (TetraSpeck Microspheres, 0.2 µm) mounted on the surface of a microscope slide were used as fiducials. The axial drift during experiments without focus lock (***Figure 12***a) is observable as a slight blurring of the beads, which is not the case for experiments with enabled focus lock. The axial sample drift was quantified by converting the lateral displacement of the back-reflected laser on the sensor into an axial distance. The displacement-to-pixel was determined in an initial calibration experiment (***Figure S51***, 1 px =*^* 561 nm). ***Figure 12***b shows the axial drift throughout the three hour experiment exceeded 400 nm without focus lock, whereas virtually no residual drift was detectable when the focus lock was enabled (*<* 10 nm). Lateral drift was quantified by tracking the fiducial beads and calculating their euclidean distance from their original position at the start of the experiment. We found the lateral drift to be similar with and without focus lock enabled, which is expected since the focus lock only compensates the axial drift. In comparison to other setups [33–35], the lateral drift is extremely low (≈ 200 nm in three hours), which is likely due to using a closed-loop piezo stage not only for the *Z*-axis, but also for *x*- and *y*-axes. Most commercial and home-built setups use servomotors for the *x*- and *y*-axes, with at least one order of magnitude worse resolution and stability [33, 36, 37].

**Figure 12.**
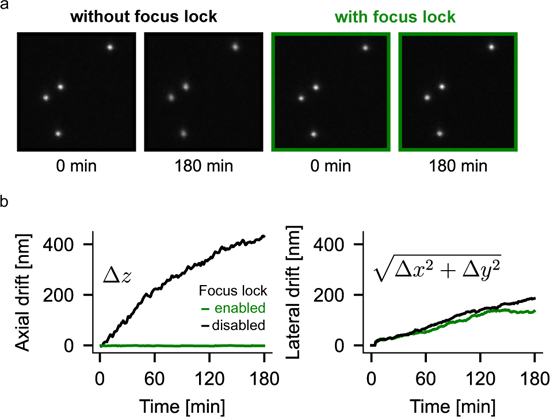
**a**) Still frames of a sample of fluorescent beads, imaged with and without qgFocus. For the experiment without focus lock, there is a slight blurring of beads observable over time, whereas the beads with the focus lock enabled stay in focus throughout the entire three hours experiment. For both experiments, the same sample was used, but before each acquisition, the sample was withdrawn from the immersion oil and re-positioned, to avoid biases due to non-identical settling times of the mounted sample. One frame was captured every 30 s for three hours. Scale bar is 5 µm. **b**) Representative axial and lateral sample drift during a three hour experiment with and without enabled focus lock. Left: Axial drift, Right: Lateral drift of tracked beads, as given by the euclidean distance of their current position versus their position in the first frame.

#### Large field-of-view and illumination homogeneity

One of the key features of the K2 TIRF microscope is its large field-of-view that is flat and homogeneously illuminated, which facilitates conducting statistically sound and quantitative single-molecule fluorescence experiments. Using a refractive beamshaper and appropriate optics, we are able to evenly illuminate a large field of view up to 190 µm in diameter, providing a high signal-to-background ratio across the entire field of view. This is especially important for dynamic live-cell or single-molecule tracking experiments (*e.g.* cellular activation essays, kinetic studies of molecular interactions), because events may be non-recurrent and capturing as much data from a single field of view as possible with little to no spatial variation in imaging quality is essential (***Figure 13***c).

**Figure 13.**
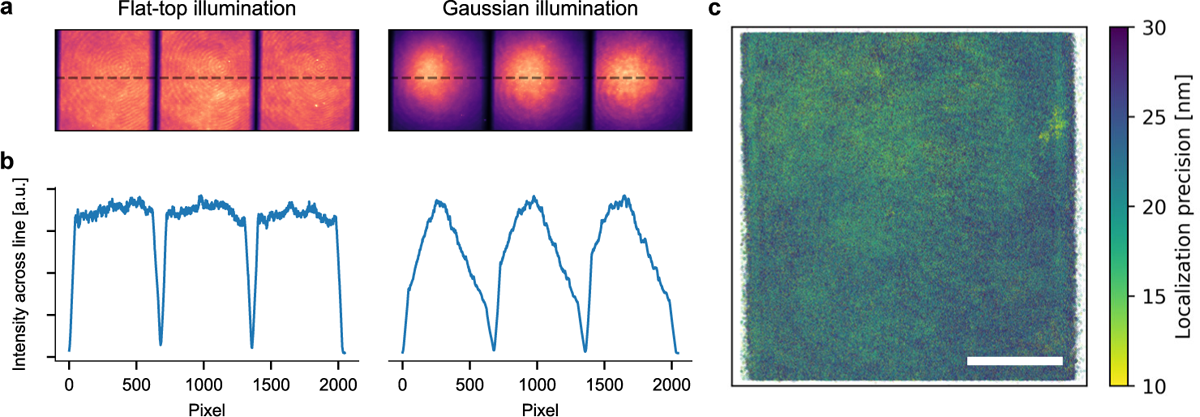
Triple-color TIRF imaging of a multicolor fluorescently labeled bilayer, excited by a spatially homogeneous excitation field (**a**) versus a Gaussian beam (**b**). Laser intensities were adjusted for each color channel to produce similar peak intensities. Fields of view per color channel are 682×682 pixels, or 73.656 µm x 73.656 µm. **c)** Localization precision map computed from a total of 3,300,599 individual localizations of freely diffusing Cy3B-labeled reconstituted proteins on a supported lipid bilayer. The homogeneous excitation field allows precise localization of single-molecules across the entire field-of-view. Scale bar: 20 µm.

In addition, the large 2048×2048 px camera chip allows the imaging up to three color channels simultaneously side-by-side without the need to invest in multiple, synchronized cameras. Simultaneous capture of multi-color emission is important for accurately tracking fast dynamic processes of multiple species of fluorescently labeled molecules in cellular and artificial environments. For triple-color simultaneous imaging, the camera is cropped to produce three equal-sized fields of view of 682×682 pixels, or 73.656 µm x 73.656 µm (see ***Figure 13***).

Furthermore, the homogeneous illumination of the field of view of the K2 microscope not only provides a high signal-to-background ratio across the entire field of view, but also enables seamless stitching of images acquired by raster-scanning the sample. This is because the lack of vignetting at the edges of the field of view allows the individual images to be easily combined into a single, large image without loss of quality (see ***Figure S54***). Furthermore, the focus stabilization system is able to compensate for coverslip tilt and inhomogeneities that can degrade the image quality during raster-scanning. This allows us to perform fully automated raster-scanned image acquisitions, without the need for manual intervention to maintain focus (see ***Figure SM3***).

### Objective heating stability and response time

We use an objective heater to maintain a stable, physiological temperature of the sample during extended live-cell imaging at 37 ^◦^C. The heater is controlled by a feedback loop that uses a temperature sensor at the objective to control the flow of current through a polyimide heater wrapped around the objective. Thermal decoupling of the objective and a plexiglass enclosure around the central cube help to maintain a constant temperature. The offset of objective temperature and the temperature directly at the sample was determined using a second sensor immersed in a test sample. At a room temperature of around 21 ^◦^C, the system stabilizes at an objective temperature of 42 ^◦^C (corresponding to 37 ^◦^C at the sample) after about 20 min (***Figure 14***). Since there is no active cooling, it takes about 200 min for the system to return to room temperature. The thermal isolation and the mass of the central cube ensure minimal heating of the rest of the setup, which is important for maintaining alignment. In fact, the environmental sensor on the central cube showed an increase in temperature of less than 1 ^◦^C when the objective was heated to 42 ^◦^C (***Figure S53***). Overall, the objective heater is an effective and low-cost solution for maintaining a stable, physiological temperature of the sample during live-cell imaging.

**Figure 14.**
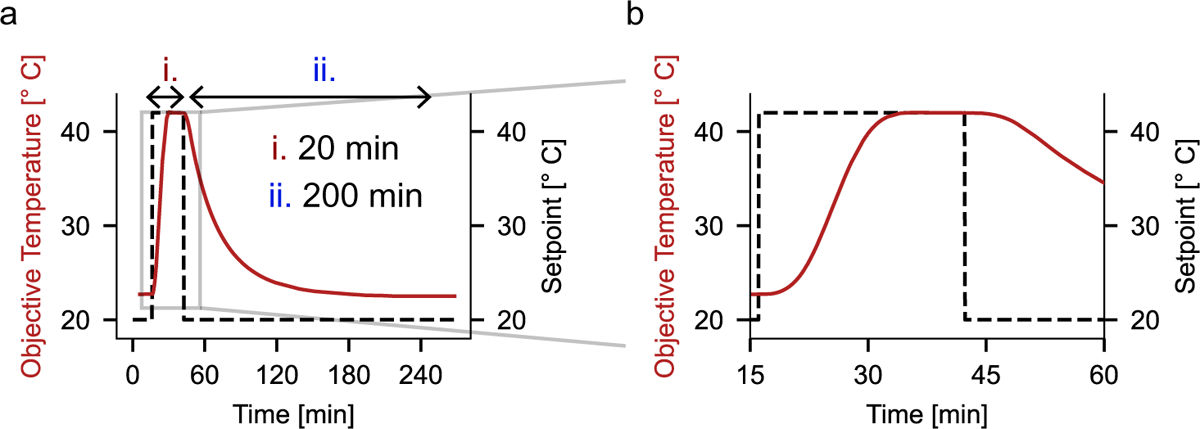
**a)** Heating phase, stabilized temperature and cooling phase during an objective heating cycle with a setpoint temperature of 42^◦^, corresponding to 37^◦^ sample temperature. **b)** Close-up showing stabilized temperature about 20 min after activating the objective heating.

**Figure 15.**
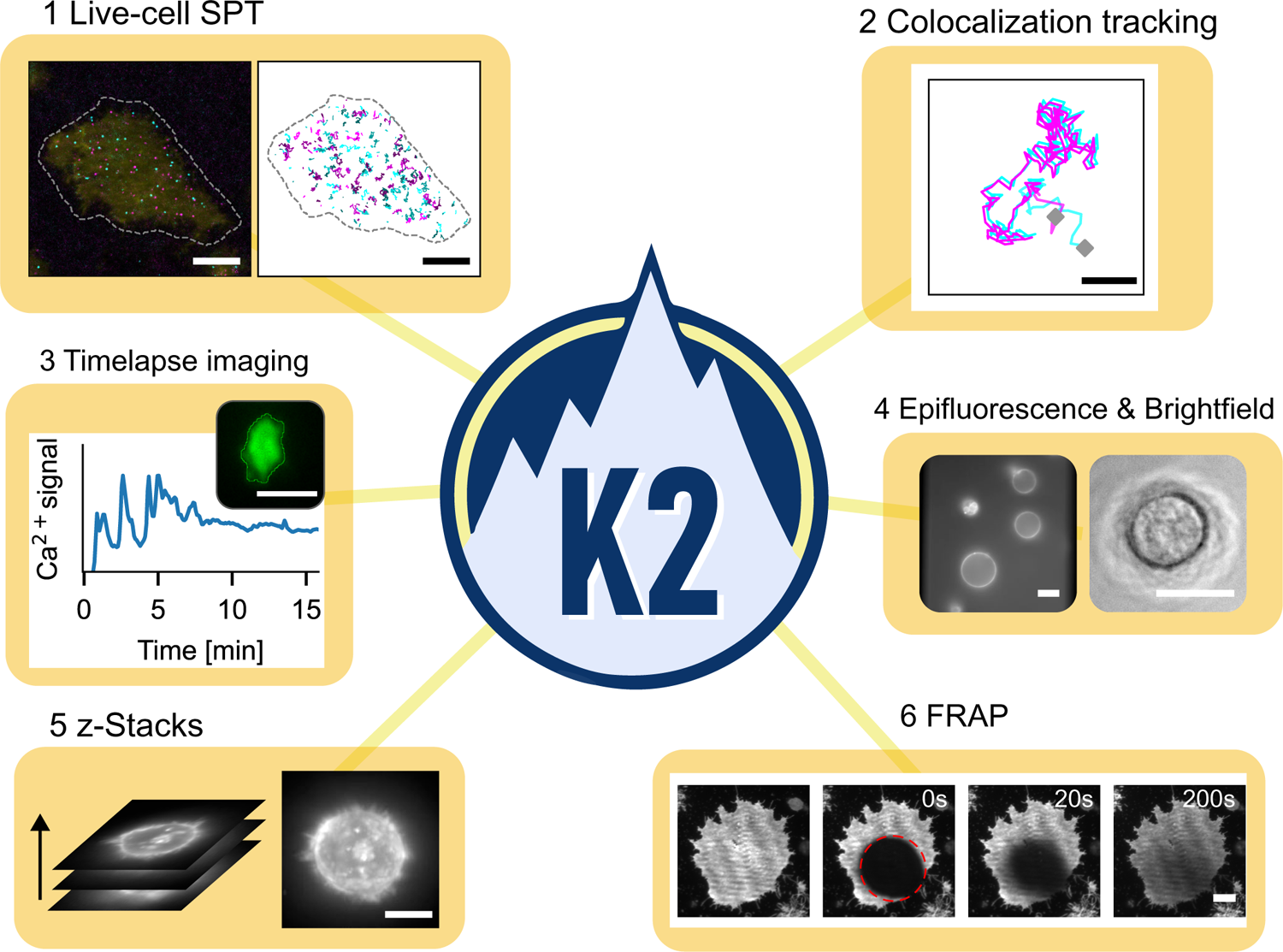
Examples of the experiments that can be performed with the K2: (1) Dual-color single particle tracking on live Jurkat T-cells with a third channel used as a transfection reporter and (2) dual-color single-molecule tracking of reconstituted proteins on a supported lipid bilayer. Tracking the movement of two (or more) different fluorescently labeled molecules provides information about the interactions between different molecules by colocalization analysis; (3) Live-cell timelapse imaging to measure immune cell activation: Jurkat T-cell outlines are tracked using cellpose [38] and TrackMate [39] and the mean intensity of the cell-permeable green fluorescent Ca^2+^ reporter Fluo4 is measured. (4) Epifluorescence imaging of giant unilamellar vesicles (left) and brightfield imaging of an adherent Jurkat T-cell (right). (5) Maximum projection of a *Z*-stack of a Jurkat T-cell in suspension, imaged in epifluorescence mode. (6) Timeseries of a fluorescence recovery after photobleaching measurements on a COS7 fibroblast cell: A reference image before the photobleaching step (left), directly after a bleaching pulse in the red circle (0 s, and the subsequent recovery after 20 s and 200 s, showing the mobility of the fluorescently labeled membrane. All scale bars: 10 µm

### Experimental applications: from live-cell imaging to single-molecule tracking

The K2 TIRF microscope can be used for a wide range of live-cell and single-molecule experiments. ***Figure 15*** showcases several such applications: dual-color single-molecule tracking of membrane proteins on live cells, with the third channel being used for detecting a transfection reporter (1) and dual-color single-molecule tracking of reconstituted membrane proteins transiently forming dimers on a supported lipid bilayer (2), as well as timelapse imaging of immune cell activation (3). Epifluorescence imaging of giant unilamellar vesicles is shown in (4), alongside bright-field imaging of an immune cell using the circular LED array. Combining epifluorescence imaging together with the *Z*-stack function allows to capture 3D volumes of cells in suspension (5). The K2 TIRF microscope is also capable of performing fluorescence recovery after photobleaching (FRAP) experiments, here shown on a COS7 cell labeled with a fluorescent membrane marker (6).

In summary, the K2 TIRF microscope is a versatile instrument that allows the study of cellular and molecular dynamics in up to three fluorescence channels with an additional brightfield option. A large and homogeneously illuminated field-of-view and the highly-automated sample positioning and raster-scanning options enable the acquisition of high-quality microscopy data with comparatively high throughput and little demand on user input. Furthermore, imaging experiments can be performed for extended durations (hours) due to the setup’s excellent stability, active focus stabilization and objective heating for live-cell compatibility. The extensive open-source documentation should allow researchers to easily recreate the setup, or parts of it, and adapt it to their specific requirements.

## Acknowledgment

We acknowledge support from AMOLF mechanical, software and electronical engineering departments, as well as from the precision manufacturing department. In particular, we would like to thank Marco Konijnenburg, Dico Kruining, Bob Krijger, Henk-Jan Boluijt, Hinco Schoenmaker, Niels Winkelaar and Jan van der Linden for their various contributions. We would like to thank Chi Nguyen for providing timelapse movies of Jurkat T-cell activation acquired on the K2 TIRF setup, and Nebojša Jukić for proofreading the manuscript. We thank Dr. Philipp Blumhardt for expert advice on TIRF setup construction. This preprint was created using the LaPreprint template (https://github.com/roaldarbol/lapreprint) by Mikkel Roald-Arbøl.

## Supporting Information

### Detailed build instructions

A general description of the design of the setup is given in the main text. In the following, step-by-step build and alignment instructions are provided. Information about individual components is available at the K2 TIRF Github page (https://ganzingerlab.github.io/K2TIRF/K2TIRF/component_table.html).

### Excitation path

**Figure S1.**
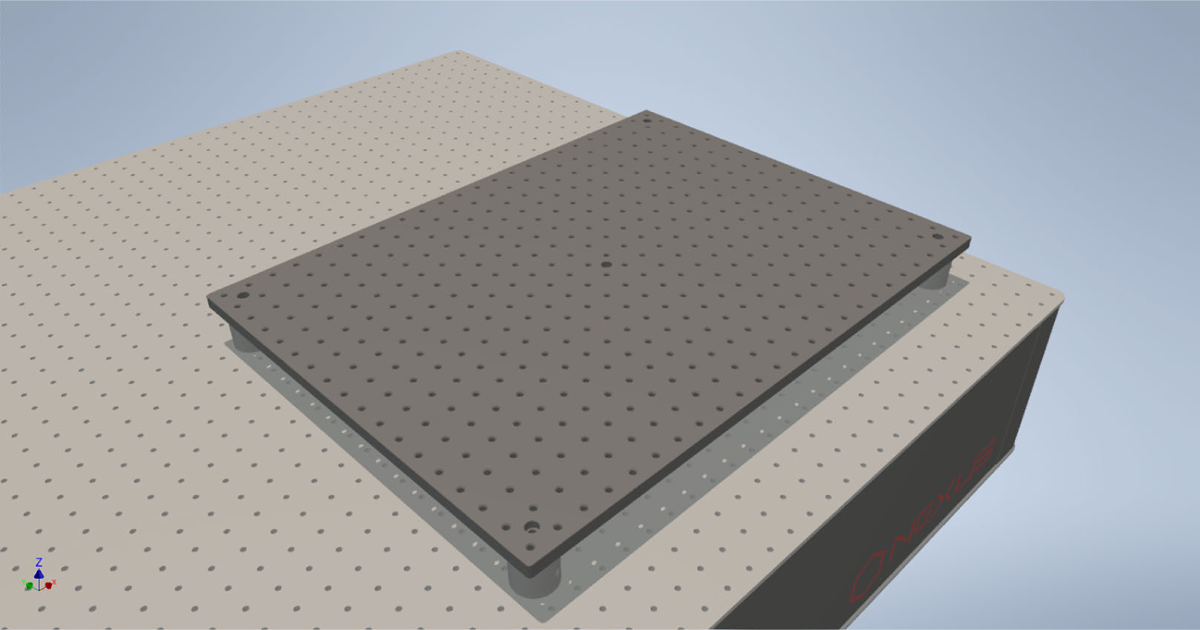
Mounting the elevated breadboard Start by freeing up optical bench space to accommodate the whole setup, including the light-protective enclosure. This requires about 190 cm by 90 cm. First, the elevated breadboard (12.7 mm thickness) for the excitation optics is mounted. To bring the excitation beam to the required height of the central cube’s entry port (117.4 mm), we use 30 mm posts and 10 mm spacers which were shortened by 0.3 mm. This brings the upper surface of the breadboard to a height of 52.4 mm above the optical table, resulting in a beam height to 65 mm above the elevated breadboard. This beam height is matched to that of the detection path, which itself is determined by the center of the emission port of the central cube (65 mm above the optical bench). Align the breadboard on the optical table with the rows of holes in the breadboard running parallel to the rows of holes in the optical table. To ensure parallel alignment, mount two pairs of irises along one row of holes on the breadboard and the optical bench at the maximal distance, and verify that a collimated laser (*e.g.* the laser from the Gustafsson alignment tool) passes through all irises. Please note that the breadboard is not magnetic and extra care has to be taken with unmounted optical parts on the surface.

**Figure S2.**
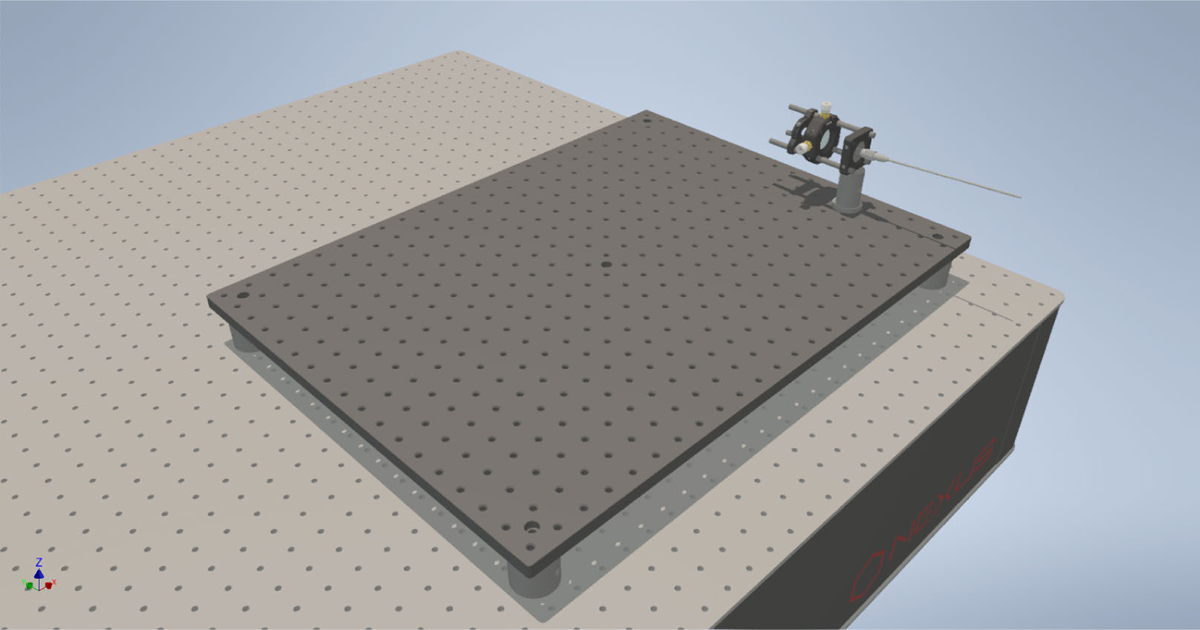
Collimating the fiber output Set up the laser box and its accessories (see Figure 2, V.1-V.5) as described by the manufacturer, or use a comparable system (*e.g*. NicoLase [22] or SMILE [14]). Then, pre-assemble the collimator cage: screw the fiber mount into the cage plate, slide in cage rods and the translating lens mount with the collimation lens. Slide on a cage iris just after the translating lens mount. Mount the collimator cage on the elevated breadboard and connect the fiber. Turn on the laser and close the iris almost entirely. Adjust the lens alignment so that the laser beam is centered around the opening of the iris, and ensure that the cage is oriented with the exiting laser beam following one row of breadboard holes. After centering the lens, the collimator lens needs to be aligned axially to collimate the laser beam. It should be placed approximately one focal length away from the fiber exit. To determine the correct position, a viewing card with a ruler may be used to view the beam close to the lens and in increasing distances. The diameter of the laser beam will not change noticeably during propagation if the beam is well collimated. If available, a shear interferometer should be used to ensure collimation as precisely as possible. If the use of a beamshaper is planned, the collimation lens local length should be chosen to achieve the required output beam diameter (pishaper: 5.9 mm-6 mm). The output diameter after the collimation lens can be confirmed with a beam profiler or any camera with a physically big enough chip. In that case, make sure to use very low laser power and neutral density filters with sufficient blocking power at the used laser wavelengths to prevent damage to the camera chip.

**Figure S3.**
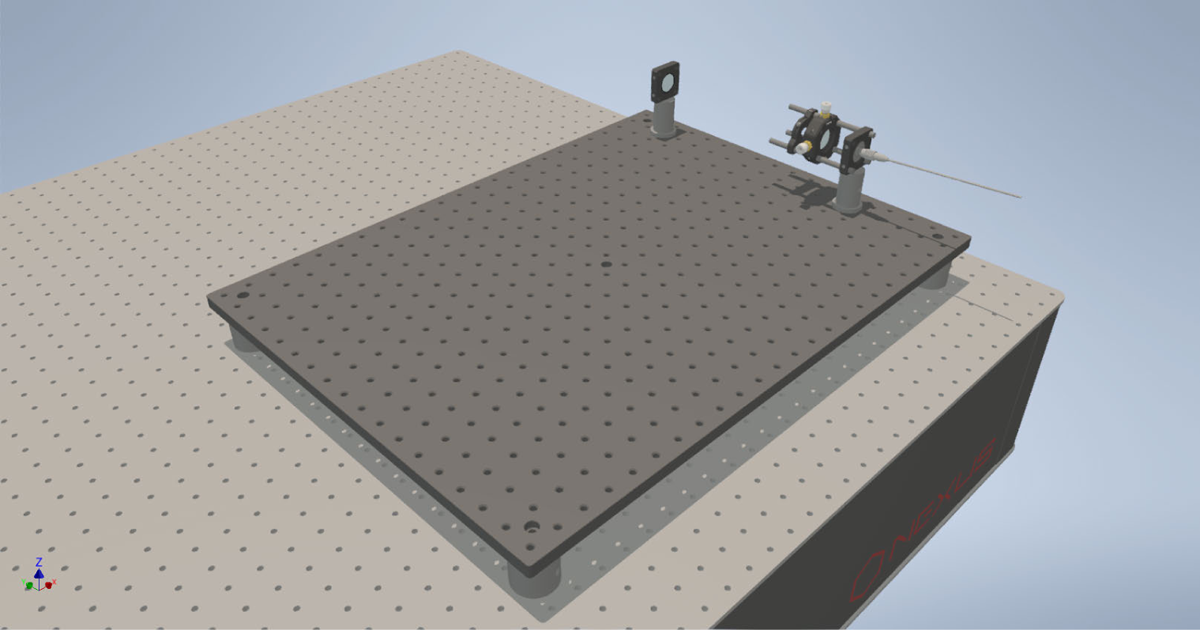
Installing the laser clean-up filter The quad-line laser clean-up filter is placed in a lens mount or cage plate and mounted on the breadboard, centered on the optical axis.

**Figure S4.**
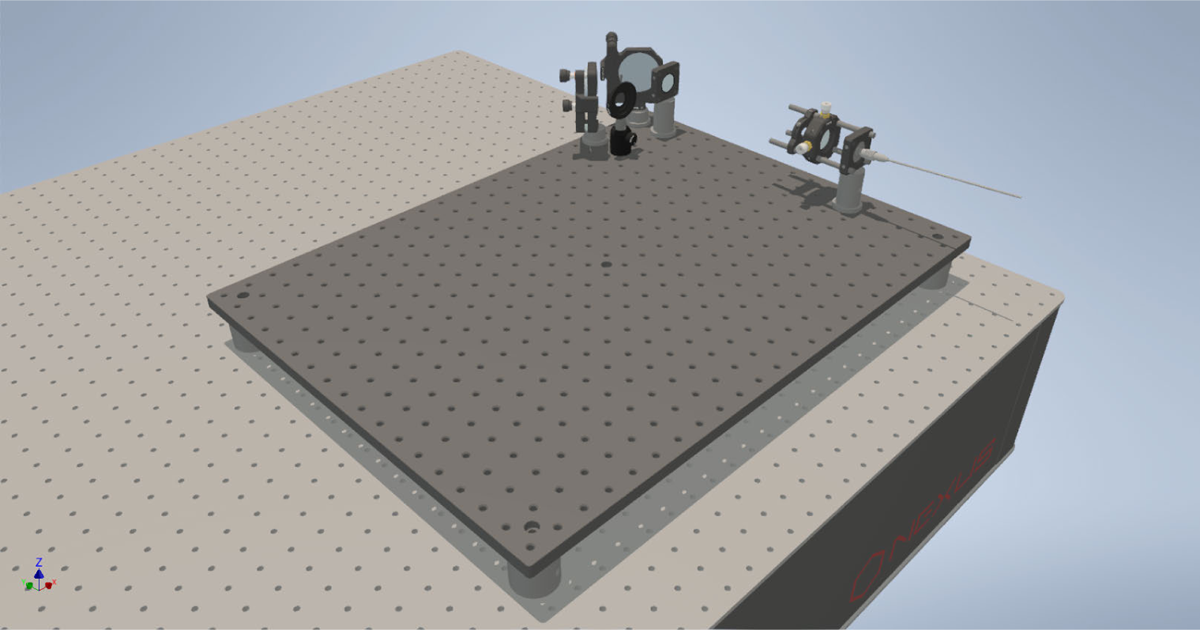
Folding mirrors and alignment irises Two folding mirrors are used for decoupling the alignment of the optical fiber with the rest of the excitation pathway. Having these two mirrors in place can drastically reduce realignment procedures afterwards, for example when the fiber is replaced or cleaned, or the collimation lens is adjusted. As long as the laser beam is collimated, the downstream alignment can always be restored by beamwalking the fiber output through two alignment irises, using the two folding mirrors. Place the mirrors roughly with their centers co-aligning with the optical axis, and facing each other at an angle of 90^◦^. Prepare two irises with their centers at 65 mm, *e.g.* by milling posts to the required length. This matches the height of the entrance of the cube above the elevated breadboard. Place two of these irises in post holders and screw them directly into the breadboard, one close to the second folding mirror, and the second iris towards the edge of the breadboard, leaving space for another folding mirror. Beam walk using the two folding mirrors until the beam passes both irises.

**Figure S5.**
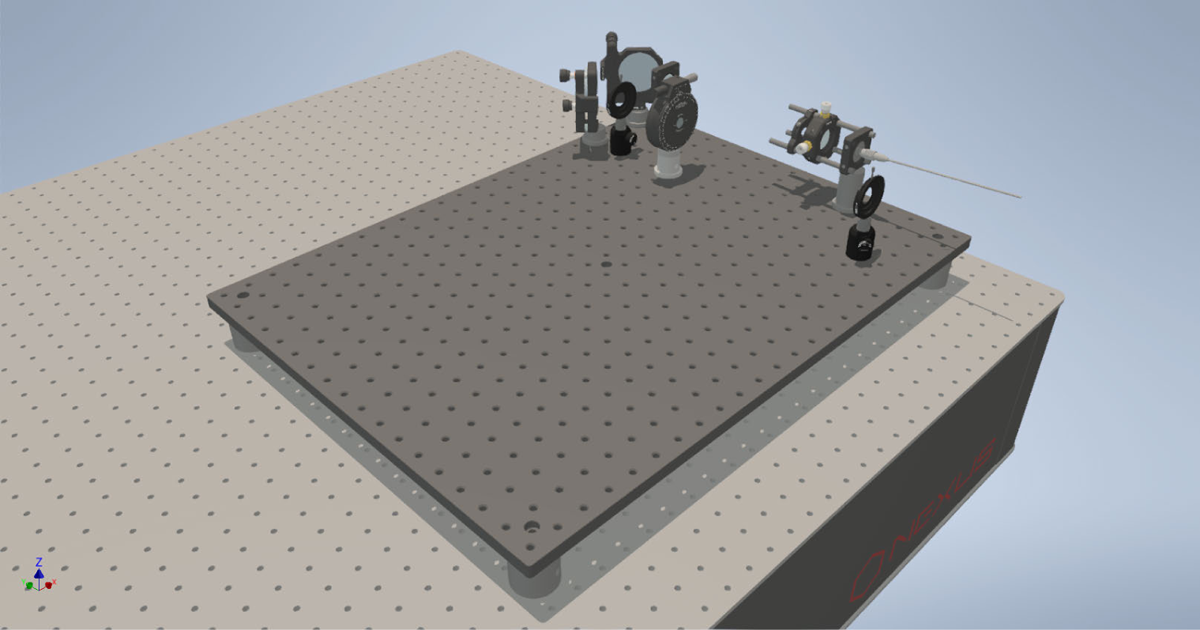
Installing the quarter-wave plate Start by determining the initial polarization of the laser beam exiting the single-mode fiber. Place a polarizer (*e.g.* LPVIS100, Thorlabs) and rotate it until maximum extinction is achieved. Then mount the achromatic quarter-wave plate in its rotation mount, and place it after the polarizer. Turn it such that the maximum extinction is maintained. From there, rotate the quarter-wave plate by 45^◦^. Center the mount and use the back-reflection to correct for tilts. Verify the circular polarization by rotating the polarizer. The intensity of the transmitted light should remain the same (*i.e.* half of the initial intensity).

**Figure S6.**
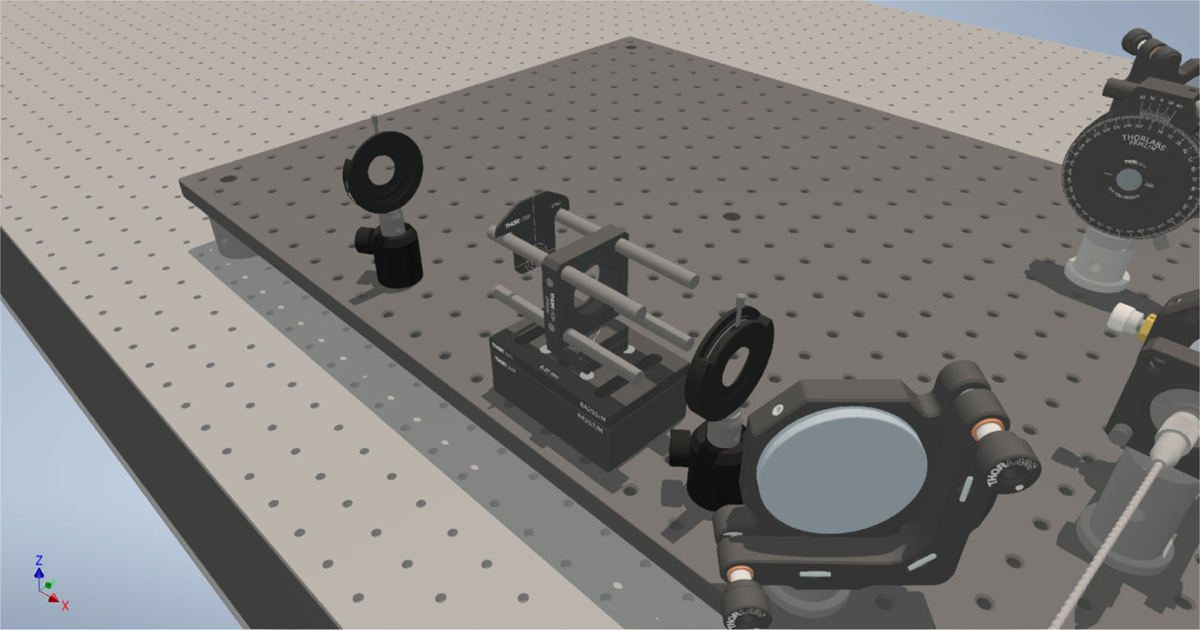
Installing the beam expander telescope Place another mirror and align it by re-purposing the two irises. The laser beam should follow a row of breadboard holes at unchanged height. Mount the spacers for the beam expander, screw the bottom part of the kinematic base in the center hole of the top spacer, and the top part on a cage plate. Take care to mount the base square with the cage plate. Slide four cage rods into a cage plate and fasten them. The center of the cage plate is now raised 65 mm above the breadboard. A cage alignment target is used to ensure that the laser beam runs parallel to the cage. If there are big deviations, correct the orientation of the kinematic mount and make sure they are both square with respect to the spacers and cage assembly. Smaller deviations can be corrected for by carefully rotating the spacers with slightly unscrewed mounting screws.

**Figure S7.**
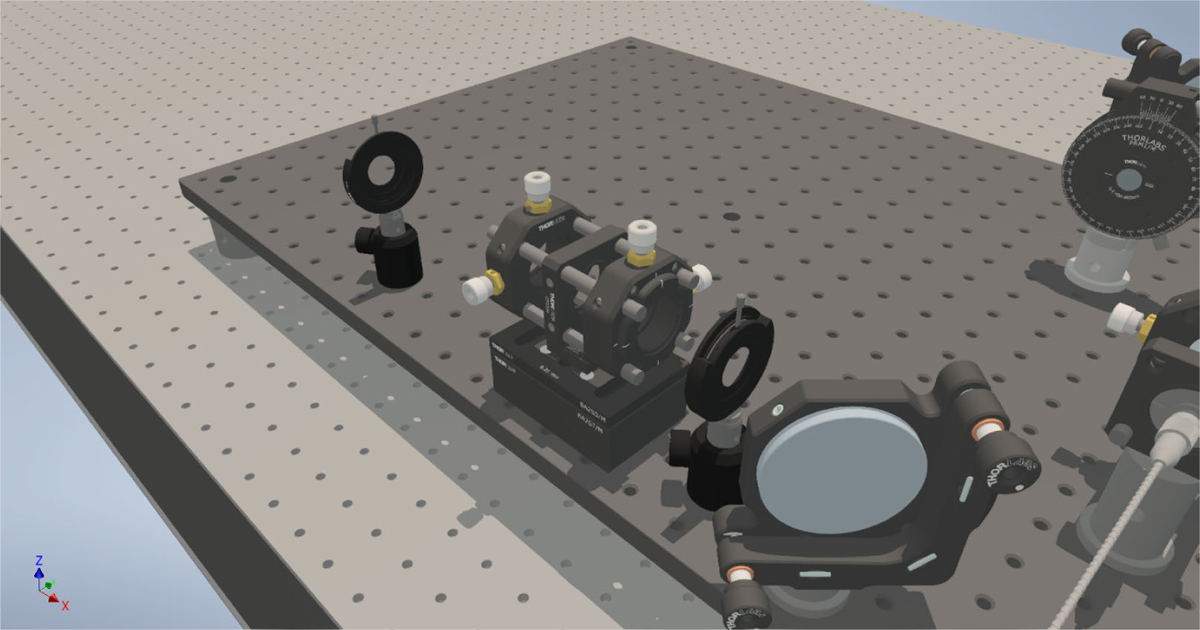
Alignment of the beam expander telescope Before placing the first lens of the telescope, mark the laser spot on a surface further downstream the optical path, or use an iris. Mount the negative focal length lens in the telescope and align the lens by centering the (convergent) back-reflection on the iris that was placed in front of the cage assembly. Additionally, this iris helps decreasing the laser beam diameter, so that the beam passing through the lens is only slightly divergent and can be used to align the lens so that the beam coincides with the marked spot. Install the second lens at a distance equal to the sum of the two focal lengths (100 mm − 40 mm = 60 mm) and align it, so that the beam hits the previously marked spot. The back-reflection can be viewed on a cage alignment plate with a pre-drilled hole. To check the collimation, compare beam diameters at different distances, and move the second lens on axis until beam diameters are equal. Another option is to place an iris near the lens, that clips the beam and use a mirror as far away as possible to reflect straight back at the iris. If collimated, the returning beam should fit exactly through the iris. Care should be taken that there is not a focus on the mirror. Ideally, if available, use a shear interferometer to confirm collimation. After alignment, make sure to fasten all screws of the telescope and the spacers tightly.

**Figure S8.**
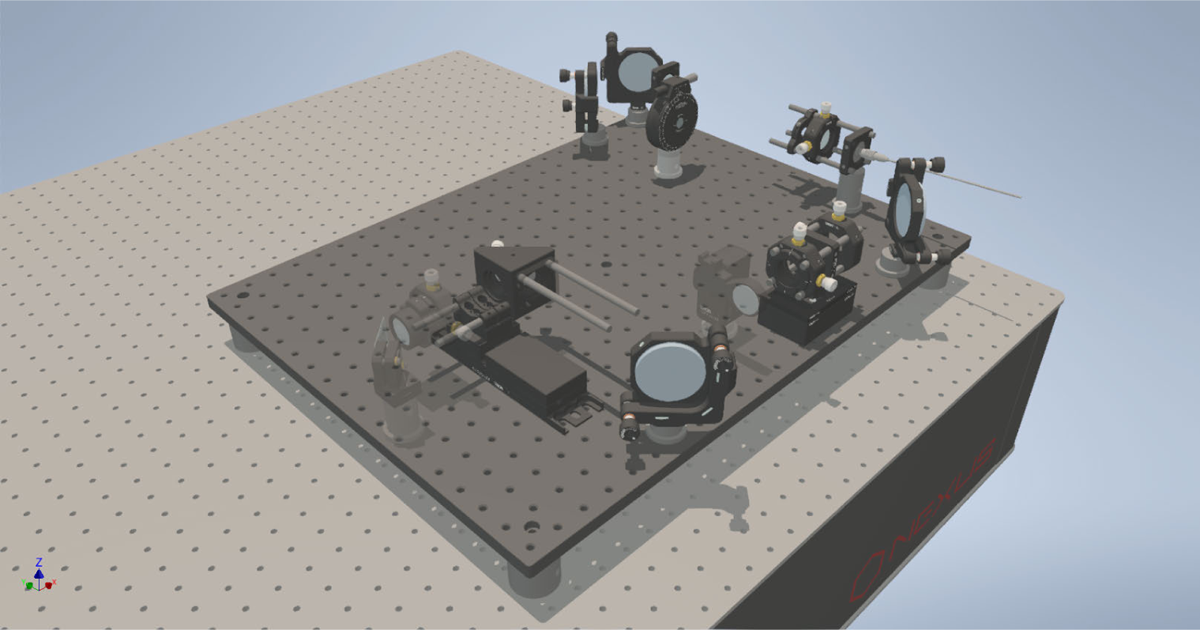
Mounting TIR stage and assembling cage system After the beam expander, leave space for the flip-in lens mount and place another mirror to guide the laser beam towards the motorized stage, which holds a cage with an right-angle mirror mount and the lens focusing the laser beam onto the back-focal plane of the objective. This stage controls the incident angle of the excitation beam onto the microscope glass slide, directing the laser beam into total internal reflection (TIR), and is therefore called TIR stage (see Figure 9). First, mount the cage system brackets on the stage and the stage itself on the adapter plates. Make sure to keep 100 mm of space for the TIR lens and the dichroic mirror that couples the infrared laser of the focus lock system. Connect the stage to the controller, which is itself connected to the computer. Place a mirror in the right-angle mirror mount and assemble the cage system.

**Figure S9.**
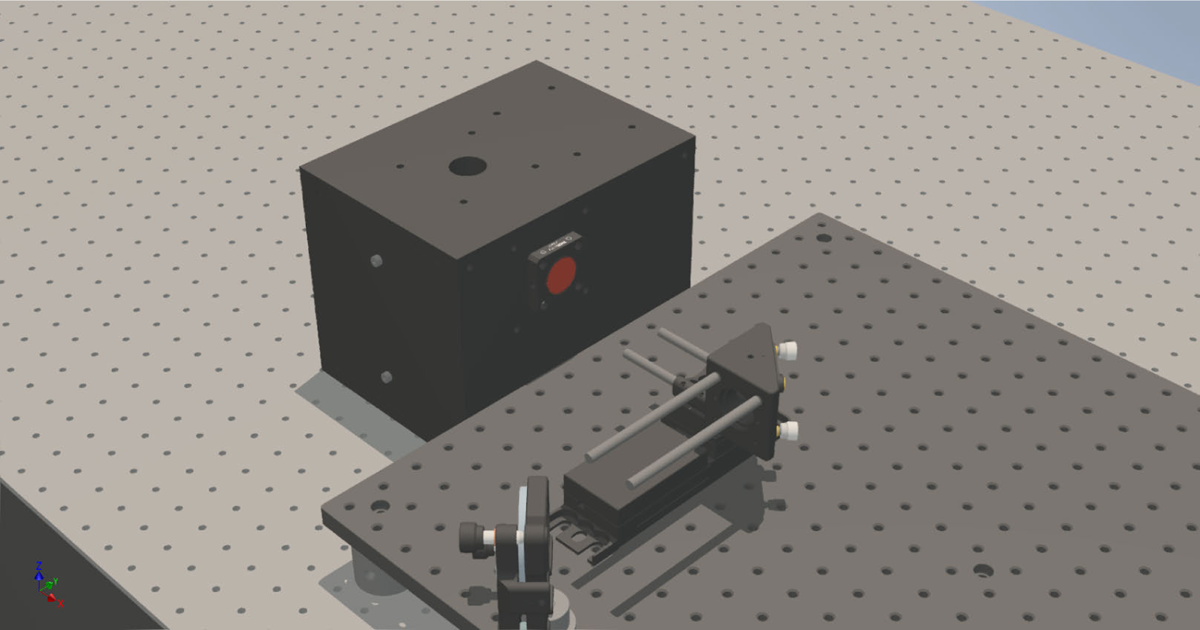
Mounting the central cube Place the central cube on the optical table, with a 10.5 mm distance from the elevated breadboard to account for the aluminum enclosures, and with the excitation entry port roughly aligned to the cage system on the motorized stage in its default position. Mount a cage plate with a fluorescent alignment disk in the excitation entry port of the cube, and use the motorized stage to move the cage assembly until the laser beam is roughly centered on the entry port. Putting the stage as close to its zero position when the beam is centered is a deliberate choice since it reduces the chances of the laser beam pointing towards the user when the stage is moved in the wrong direction during an illumination mode switch. Fasten the central cube and the stage securely.

**Figure S10.**
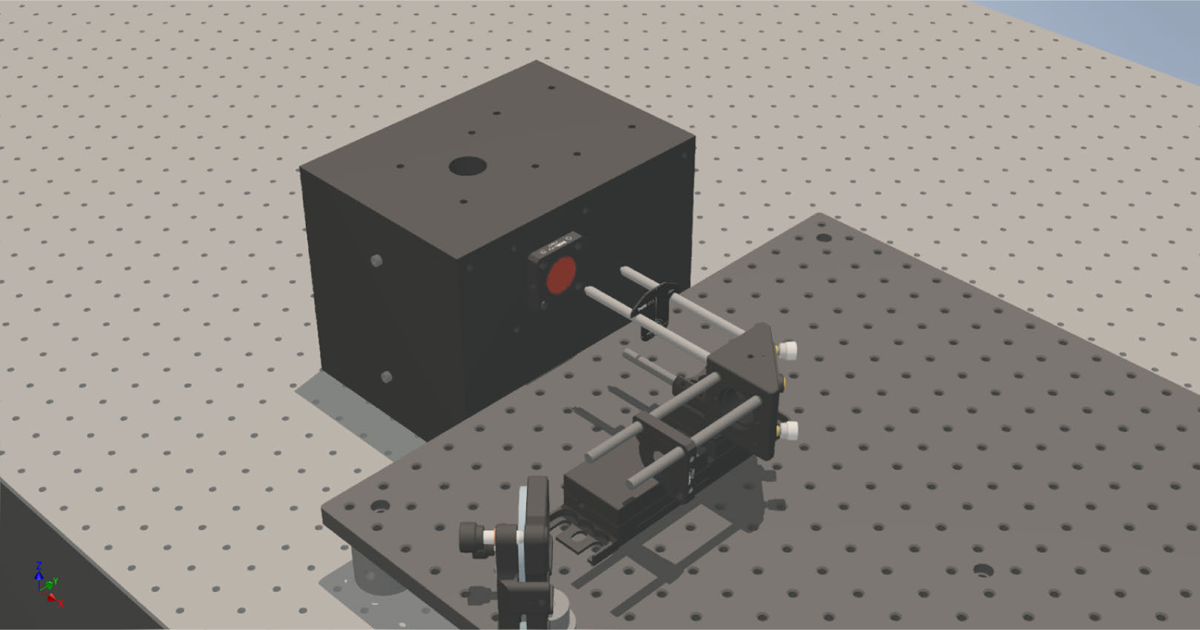
Launching the laser beam into the central cube Mount a cage alignment plate or a sliding cage iris onto the cage rods pointing towards the previous mirror in the excitation path. Use both the position and the angle of that mirror to launch the laser beam centered and parallel to the cage axis. The position of the laser beam should not change when the iris or alignment plate is slid along the cage rods, and there should be no cropping of the laser beam when it enters the right-angle mirror mount. Place a cage alignment plate on the cage rods pointing towards the cube and ensure that the right-angle mirror launches the laser beam centered and parallel to the cage: First, slide the alignment plate close to the right-angle mirror mount and verify that the laser beam stays centered when moving the motorized stage. Then slide the reticle towards the central cube and correct the angular misalignment using the right-angle mirror mount. Finally, use the motorized stage to center the beam again on the fluorescent alignment disk mounted on the central cube’s entry port.

**Figure S11.**
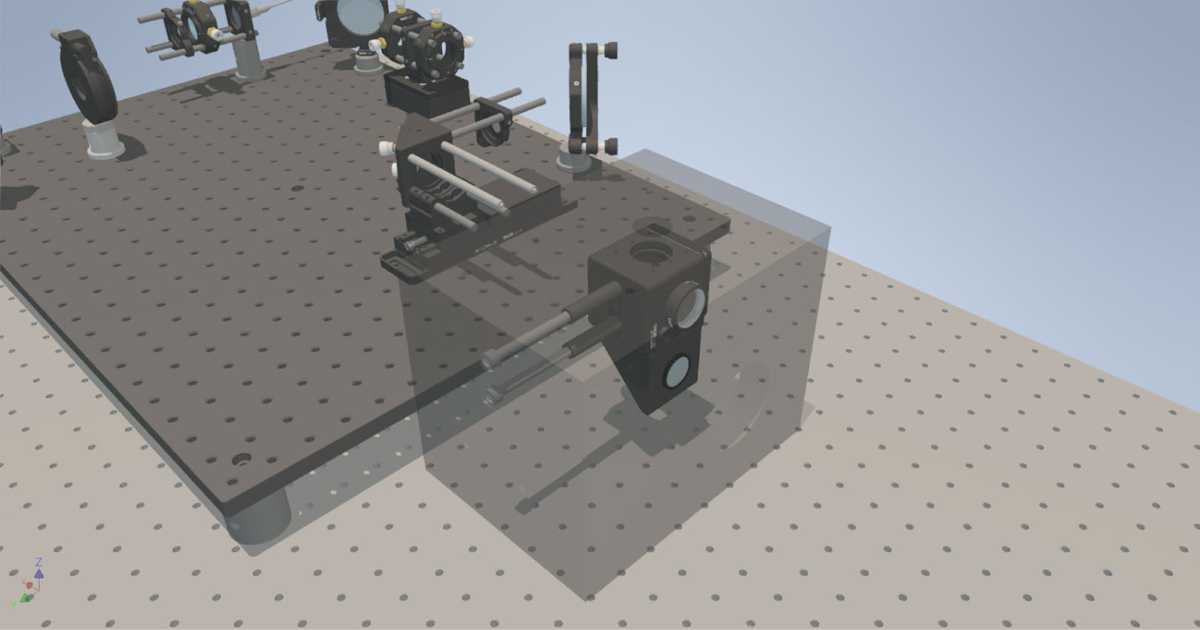
Assembly of the central cube interior parts *Warning: The following steps need to be done very carefully with the appropriate laser goggles, since the collimated laser beam can potentially hit and harm the eye.* Place the dichroic mirror inside the dichroic mirror cube, mount the connecting bracket and the right-angle mirror mount with the mirror placed inside. In the dichroic mirror cube, mount the neutral density filter that absorbs residual, undesired transmission of laser light through the dichroic mirror. Mount the quad-band filter on the right-angle mirror mount, to prevent residual excitation laser light in the detection path originating from back-reflections at the sample. Install the assembly in the cube using the 40 mm threaded spacers.

**Figure S12.**
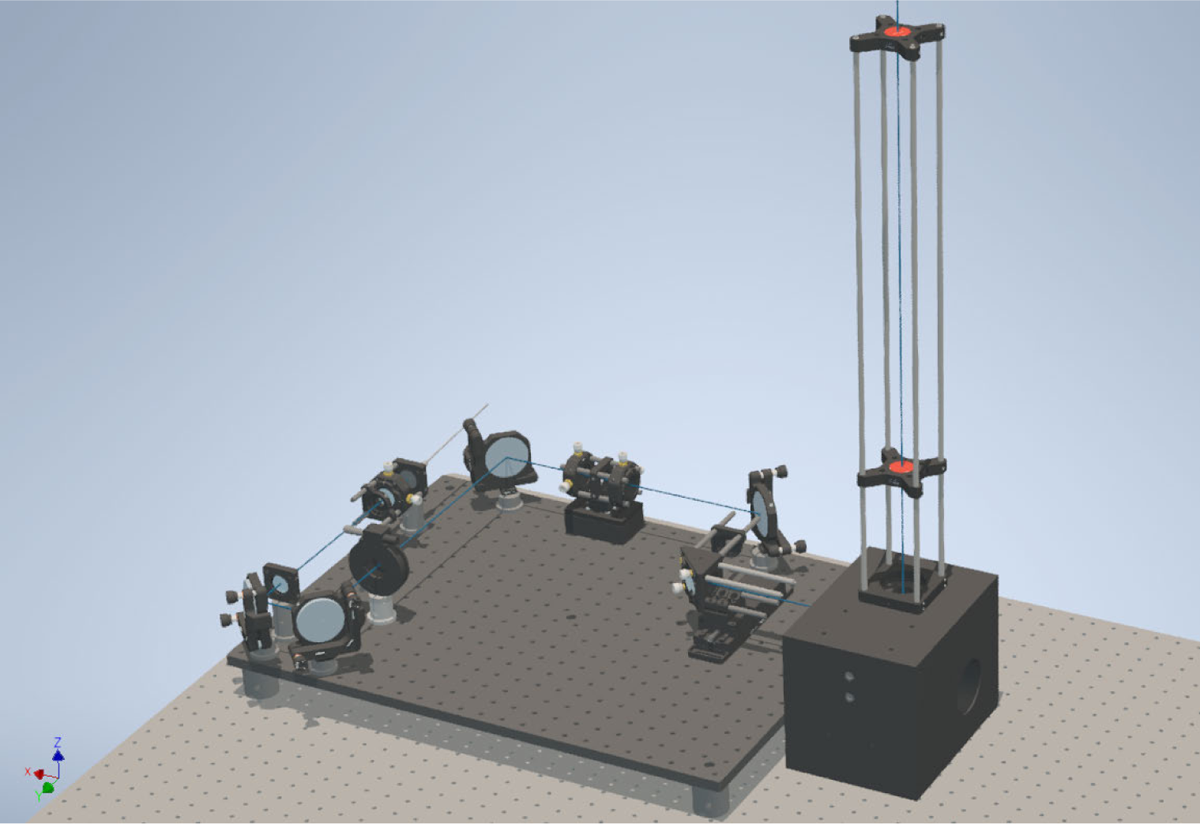
Alignment with respect to the central cube The laser beam needs to exit the cube from the center of the objective hole and perpendicularly to the upper surface of the central cube, after being reflected off the dichroic mirror. The center position and angle are verified by mounting a long cage system directly onto the cube. The cage system features a sliding fluorescent alignment disk in a cage plate. The position of the laser beam on the sliding plate should remain in the center when sliding away from the objective port. If this is the case, mark the point where the laser beam hits the ceiling for later reference. As an alternative to the cage system aligner, a thread with a weight can be strung from the point on the ceiling where the laser beam is visible. The thread should meet the center of the objective port if the alignment is correctly performed, and assuming that the optical bench is leveled. If the laser beam is not centered, or not exiting perpendicular, previous alignment steps must be revisited.

**Figure S13.**
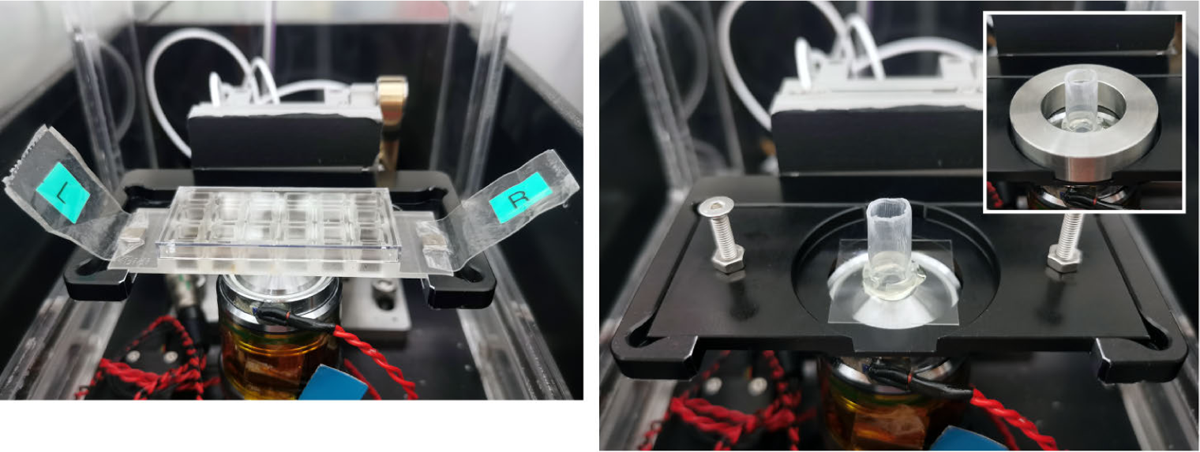
A multi-well imaging slide (left) and a cut Eppendorf tube glued onto a glass slide (right) are mounted on the sample holder and held in place using small magnets or a aluminum ring. Sample stage and sample holders The objective and the sample stage can now be installed. Screw in the objective on its thermal spacer, mount the stage on the central cube following the manufacturers instructions and connect it to the computer. Install the custom-made sample holder, add a drop of immersion oil onto the objective and place a coverslip.

**Figure S14.**
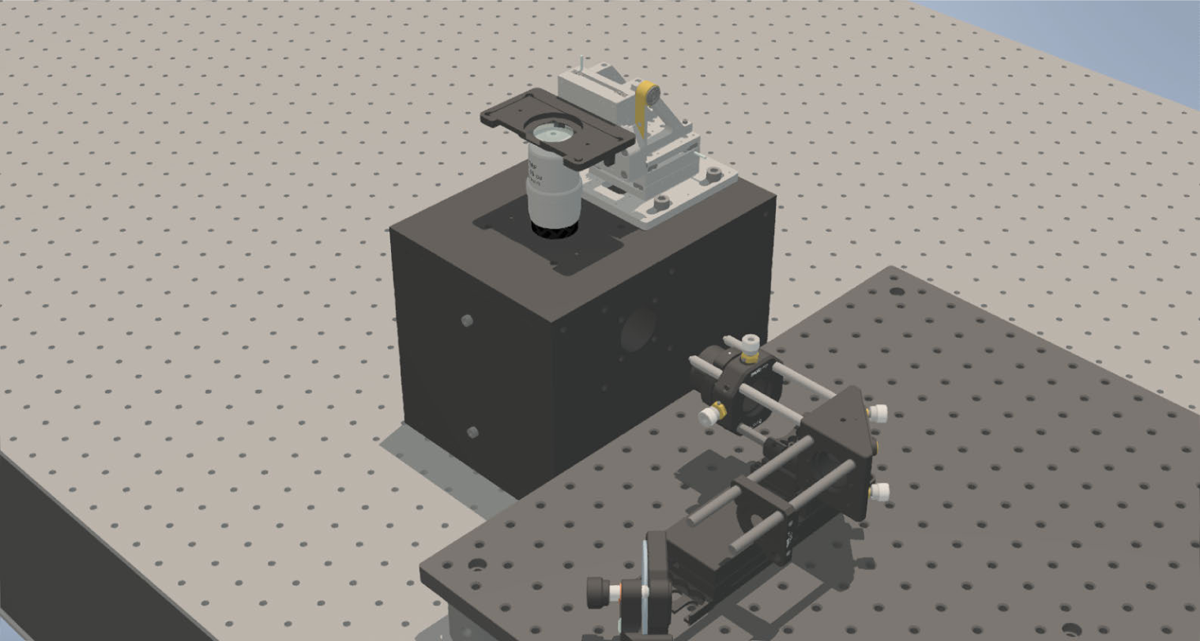
Collimation of the laser beam exiting the objective Install the lens that focuses the laser beam onto the back-focal plane of the objective (TIR lens). It should be placed roughly 115 mm away from the cube onto the TIR stage cage system. A *xy* translating lens mount with a quick release plate is used to be able to change the TIR lenses without realignment, and it also brings the lens as close to the central cube as required, without colliding with the dichroic mirror that couples in the focus lock infrared laser. In future iterations, the distance between the entry port of the central cube and the objective port will be reduced. This will relax the space constraints emerging from the short focal length of the TIR lens, which itself is required to enable the illumination of a big field-of-view. Fine-adjust the axial position such that the laser beam propagates from the objective towards the ceiling with unchanged diameter. Center the lens on the laser beam using the translating lens mount, so that the laser beam hits the marked spot on the ceiling. For correcting angular deviations, carefully use the translating mount of the TIR focusing lens. Note down the reading on the TIR stage as the default position for epifluorescence illumination. With the main excitation pathway installed, you should be able to follow the laser beam from epi- to TIRF-illumination by moving the TIR stage about 4.5 mm away from the epifluorescence setting. After moving the stage another 0.5 mm, the beam will start being clipped by the objective’s aperture. Two remaining parts, the refractive beamshaper and the FRAP flip-in lens, will be installed and aligned once the detection path is set up.

### Detection path

**Figure S15.**
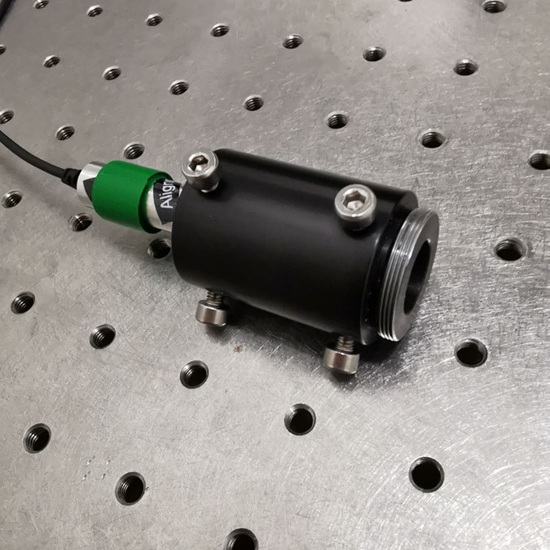
Gustafsson alignment tool The Gustafsson alignment tool is a convenient and effective way to facilitate setting up the detection pathway, and ensuring that the detection pathway is properly aligned, which is essential for accurate imaging of the sample. It consists of a metal cylinder that houses a small diode laser, providing a collimated, visible laser beam. The laser is aligned to the optical axis using set screws, and the assembly is screwed into the objective thread. The wavelength of the collimated laser is chosen to fit in the first emission window (GFP-like) of the emission dichroic mirror. Further information about assembly and alignment is found in Abrahamsson et al. [27].

**Figure S16.**
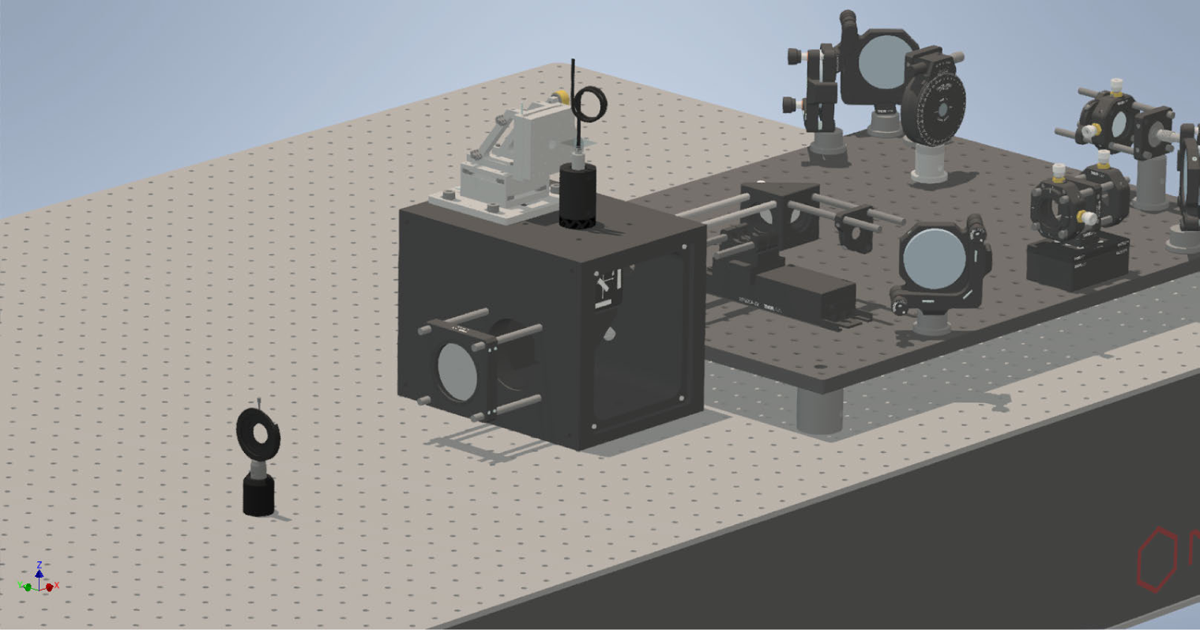
Aligning the right-angle mirror in the cube Exchange the objective with the pre-aligned Gustafsson tool and install cage rods on the detection side of the central cube using M6 thread adapters. Mount two irises, one about 250 mm away from the central cube and one at a long distance (*>* 600 mm) in the row of breadboard holes, centered with respect to the exit port of the cube. As before, use iris posts with the required length to bring the iris centers at the height of the optical axis (65 mm). Verify that the exiting laser beam passes through the first iris, and use the right-angle mirror in the cube to adjust the angle, making the laser pass through the second iris as well. Mark the spot where the laser shines onto the wall for later reference.

**Figure S17.**
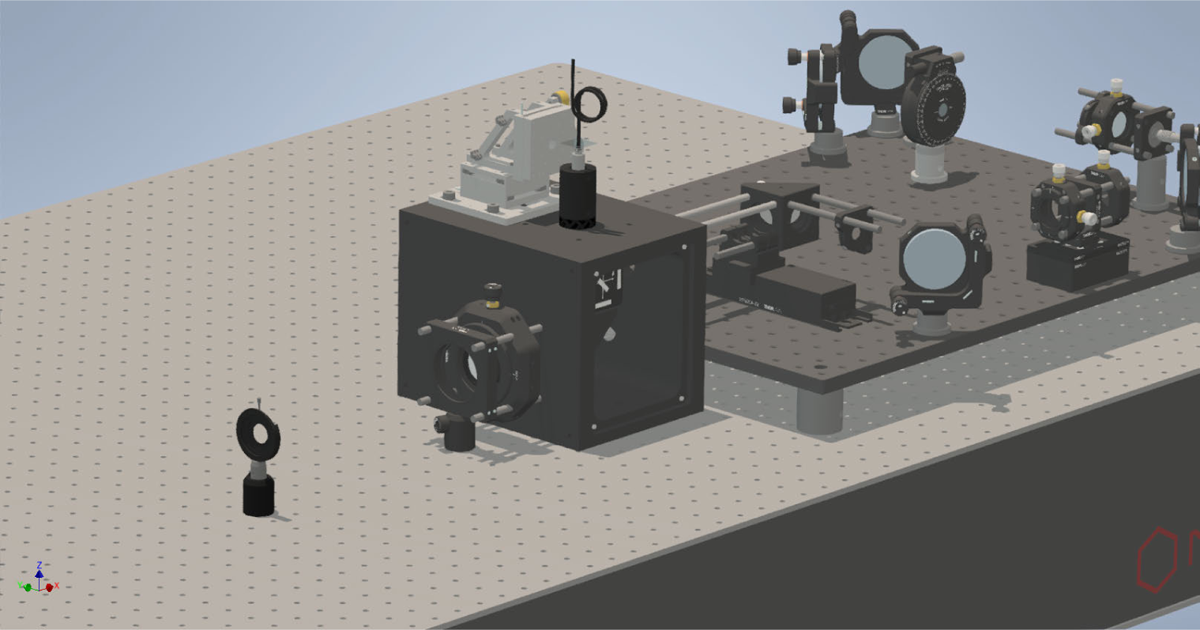
Installing the tube lens Install the tube lens on the translating lens mount placed on the cage rods and slide it into the cube, to minimize the distance between the objective and the tube lens. Make sure that the infinity-side of the tube lens is facing the objective. The cage rods are supported by a cage plate mounted to the optical bench. Align the tube lens using the translating mount by centering the beam on the first iris, which should be roughly in the focus of the laser beam.

**Figure S18.**
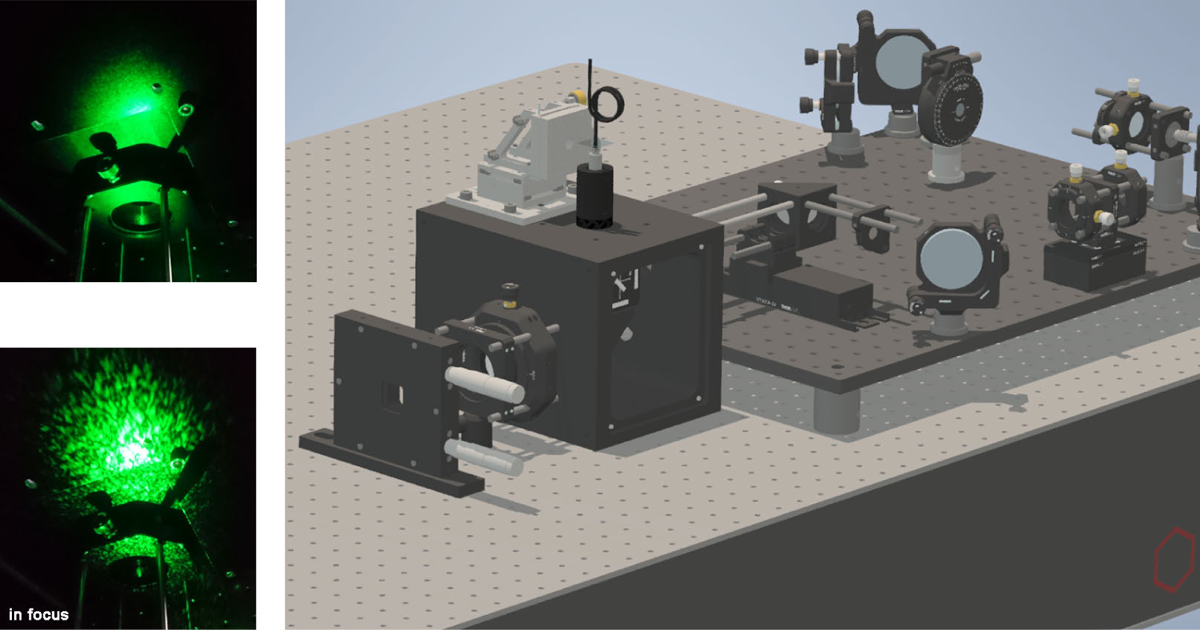
Installing the slit aperture *This may be skipped until after the placement of the camera, since it is very easy to get a sharp image of the razor blades on the camera. A line profile plot can be used to get it as sharp as possible.* The slit aperture restricts the field-of-view such that the final projected images from the three different color channels don’t overlap on the camera chip. It consists of two razorblades mounted on Vernier micrometer stages. The slit aperture is placed in the image plane, formed by the tube lens. In order to determine the exact position of the image plane formed by the tube lens, we make use of the collimated laser beam provided by the Gustafsson tool, and an object with diffusive rough surface, such as the non-shiny part of aluminum foil, or the backside of optical mirrors. Observe the speckles created by the laser beam shining onto the rough surface, in a distance of about 200 mm after the tube lens (the iris used for the previous step may need to be removed). The focus of the laser beam, which corresponds to the image plane formed by the tube lens, is where the speckles are the biggest [40]. Mark this position on the optical bench. Place the slit aperture at the marked spot, close the aperture equally from both sides leaving only a small opening. Make sure to center the aperture on the laser beam. Then, secure the aperture and open it fully.

**Figure S19.**
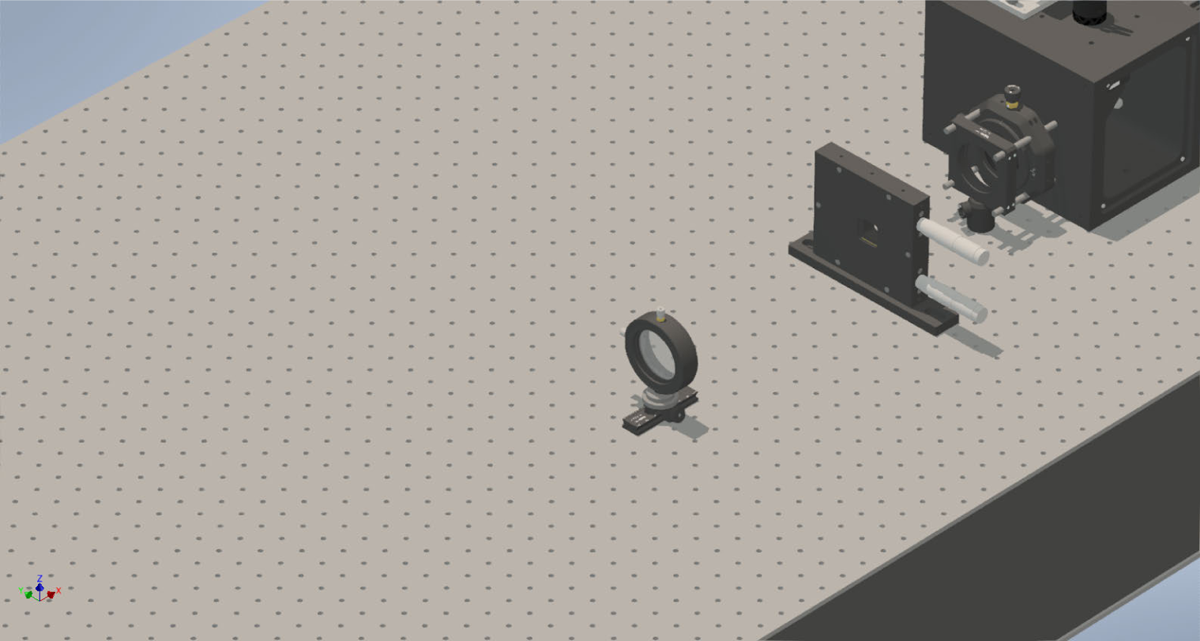
4f-system: first lens Mount the first lens of the 4f-system on a rail roughly one focal length away from the slit aperture (300 mm). Center the lens by viewing its back-reflection on an iris placed in between the 4f-lens and the aperture, and additionally verify that the laser beam hits the spot that was marked before the tube lens was placed. Align the lens axially such that the laser beam is collimated. Preferably, use a shear interferometer to confirm, or view the beam diameter at different distances away from the lens.

**Figure S20.**
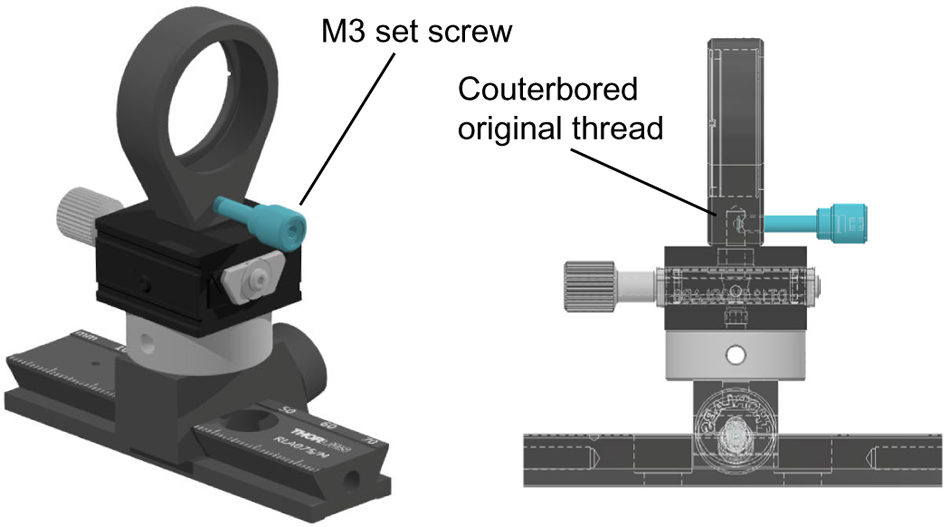
Lens mount modification A modification to the LMR1 lens mount (Thorlabs) is necessary to enable its correct alignment. The connection of the lens mount and dovetail stage is made with a grub screw and the angle of the lens mount with respect to the translation mount can not be adjusted. Therefore, the original mounting hole of the lens mount is counterbored with a 3.9 mm drill, and a new threaded hole (drill with 2.5 mm, M3 threading) is prepared perpendicular to it. This way, the lens mount spins freely on the dovetail mount and can be aligned with its optical axis parallel to the the translation mount below. Once aligned, a M3 set screw fixes the lens mount in place.

**Figure S21.**
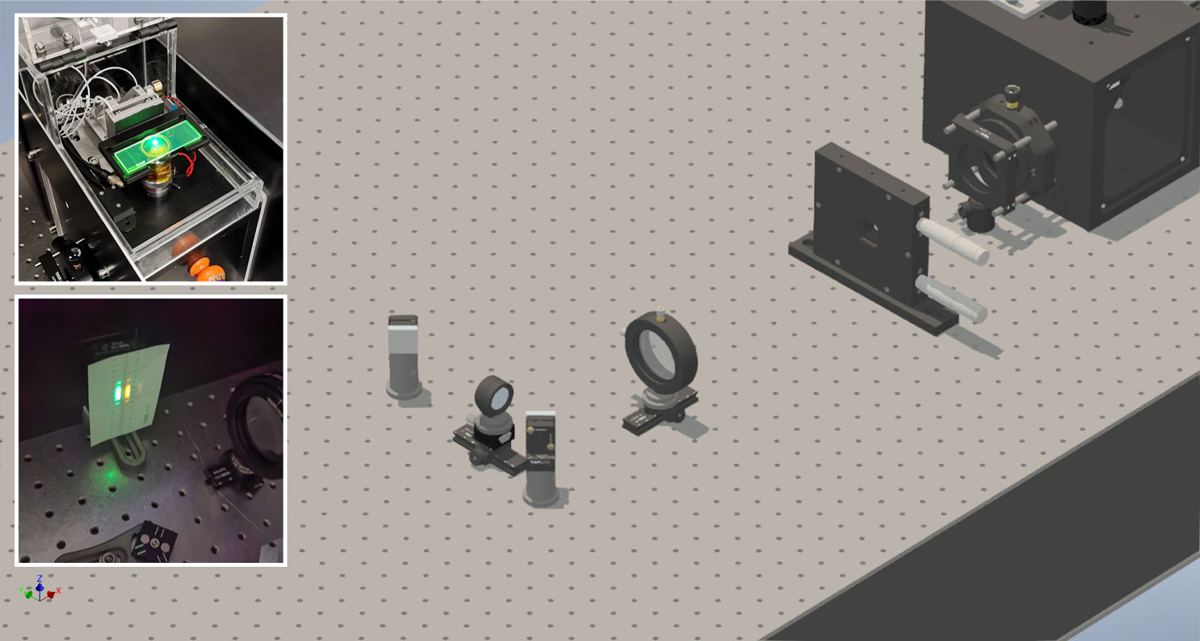
Image splitter pathway: left channel Mount the three rails for the lenses with a distance of 37.5 mm (1.5 breadboard hole rows) to each other. The outer rails can be screwed directly into the optical table, the inner rail should be fixed parallel in between two rows of breadboard holes using rail clamps. Assemble the rail carriages, dovetail stages and (modified) lens mounts. Then, place the leftmost mirror such that the laser beam hits the mirror centered and continues to run centered along to the rail. For the initial placement of the mirror, it helps to put the objective back in place and mount an acrylic fluorescent microscope slide. When illuminated with a high laser power, the fluorescence emission is clearly visible and can be used to facilitate proper placement of the optical elements on the emitted light. Relying only on the small diameter of the Gustafsson laser beam can lead to placements that show vignetting or clipping of the fluorescence emission. Mount the lens and fine-adjust the mirror position and angle using the back-reflection of the lens, since the lens lateral position is fixed by its mount and can not be adjusted. Verify that the back-reflection stays centered when sliding the lens mount along the rail. Then, place the second mirror which directs the emission light towards the camera. The exact position will be adjusted in the next step.

**Figure S22.**
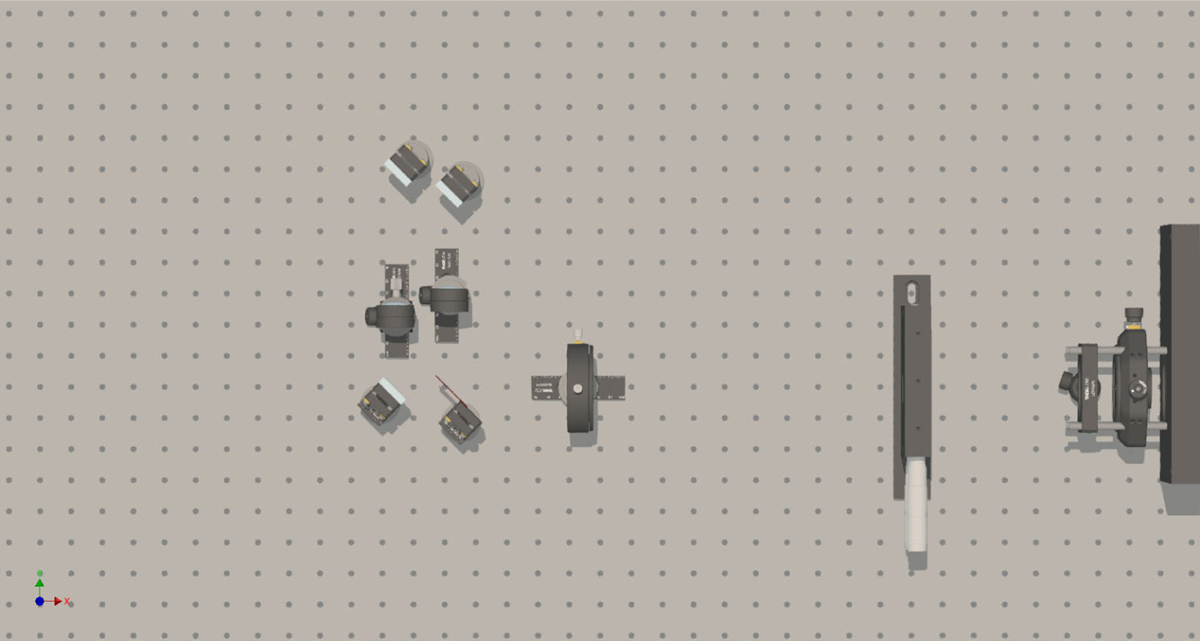
Image splitter pathway: middle channel Mount the dichroic mirror, the lens and the second mirror, using the Gustafsson tool and a fluorescent slide as described in the previous step. Since the middle channel is used as the reference channel for the two other channels, the lens does not require as precise of an axial alignment and is therefore not mounted on a dovetail stage. The mirror just before the camera of the middle channel should be aligned at a 45^◦^ angle with respect to the breadboard holes, so that the alignment laser beam runs parallel along a row of holes. Use the acrylic fluorescent slide to ensure that the mirror of the left channel does not cast a shadow in the path of the middle channel.

**Figure S23.**
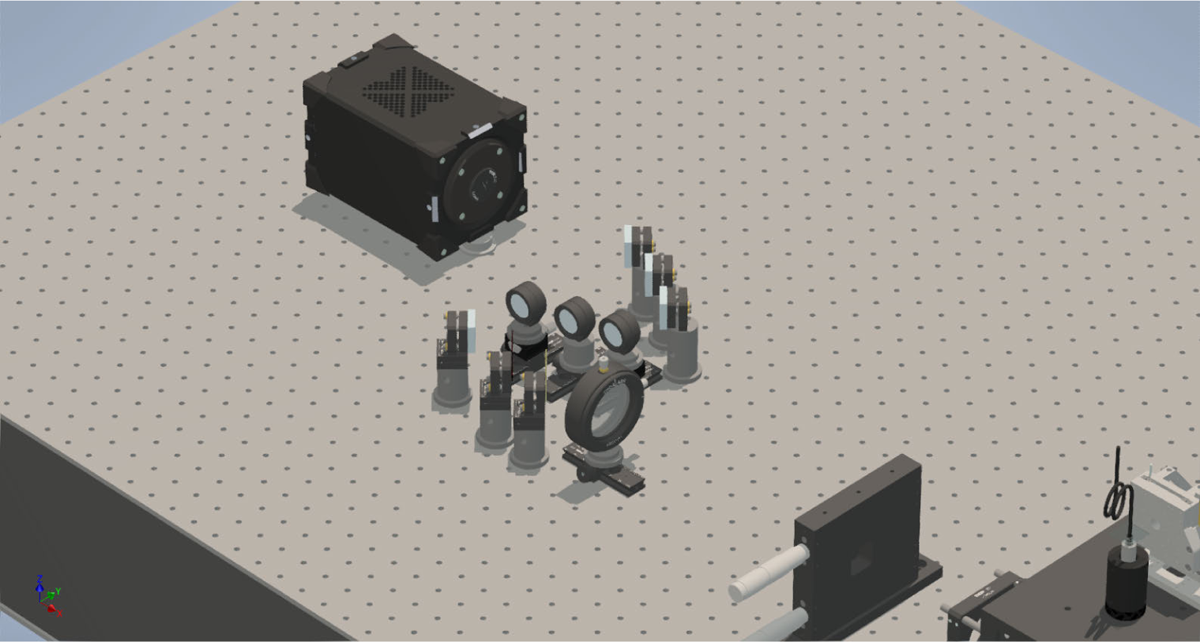
Placing the camera The wavelength of the Gustafsson alignment tool is chosen such that it is reflected by both dichroic mirrors in the triple-color detection path, and thus can be used for the entire alignment. Since the middle channel is the reference channel, it will be used to align the camera, before the rightmost channel is installed. For the simultaneous multicolor measurements, the sensor is typically cropped to a third of its vertical dimension. Therefore, make sure that the camera is oriented in the correct way (chip line-by-line readout direction parallel to the cropping axis) so that cropping the chip will reduce the chip readout time. Also, leave space for the magnetic mount and the mirror for the single-channel detection path. The camera is mounted on custom pedestal posts (10.5 mm thickness) and clamps, elevating the center of the chip to the beam height of 65 mm. With the lasers switched off, exchange the protective cap of the camera with a metal alignment disk to prevent damage of the chip by laser irradiation. With the help of the alignment disk, place the camera so that the laser beam of the middle path is centered on the alignment disk. The focus position can be determined approximately using the previously described speckle method.

**Figure S24.**
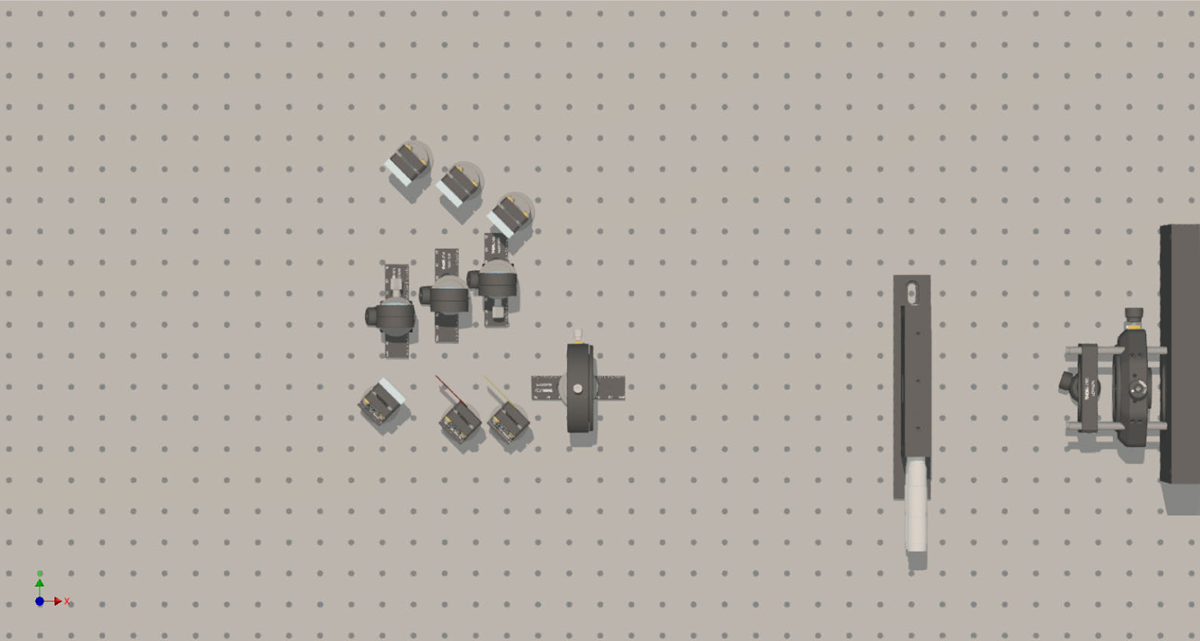
Image splitter pathway: right channel Finally, install the optics for the rightmost channel, as described for the previous two channels.

**Figure S25.**
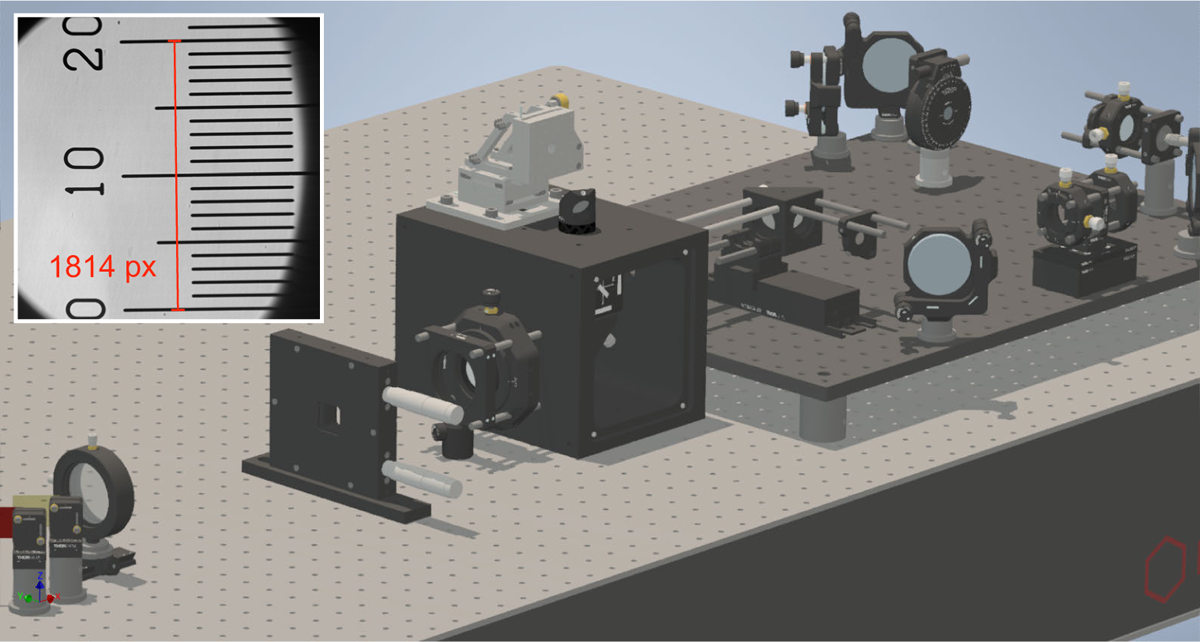
Axial alignment of the camera To determine the correct axial position of the camera, replace the Gustafsson tool with a 45^◦^ mirror on the objective port and block the left and right channels. Image an object as far away as possible. Move the camera carefully axially until the image is roughly in focus, then mount the camera securely. Then, move the second 4f-lens of the middle channel on the optical rail, to bring the image precisely in focus. By moving the first 4f-lens, one could also bring the image into focus on the camera, but it would change the final magnification. After screwing the objective back in, the theoretically expected 60x magnification (*i.e.* 1px = 108 nm) should be verified by imaging a test target slide (see inset: 0.01 µm grid. 200 µm projected at a 60x magnification onto 6.5 µm pixels would translate to a distance of 1846 pixels. Our alignment results in a distance of 1814 pixels, corresponding to a 58.95x magnification in our setup.

**Figure S26.**
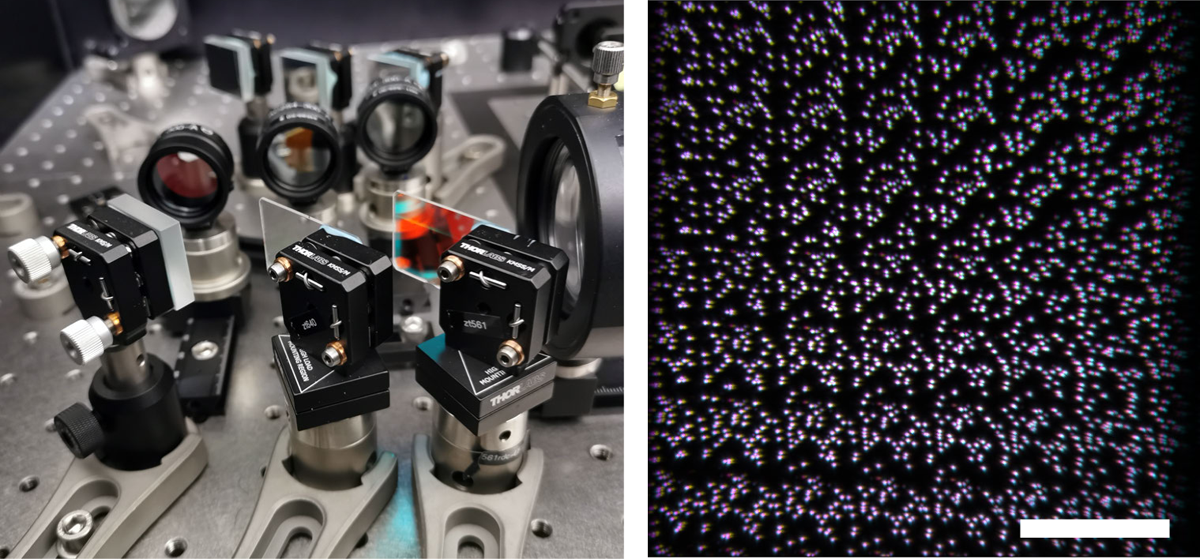
Left: Close-up of the triple-color splitter pathway. In the front, two dichroic mirrors on magnetic mounts and one dielectric mirror split the incoming fluorescence emission into three spectrally separated channels. The magnetic mount allows for easy changing of the dichroic mirrors, to allow multicolor imaging of different combinations of fluorophores, without requiring re-alignment. Each channel features a lens mount precisely positioned along the optical axis, with an additional bandpass filter screwed onto the lens mount. In the far background, the three dielectric mirrors can be seen which guide the fluorescence image onto the camera chip (left, not shown). **Right:** Superposition of multicolor fluorescent beads imaged with triple-color detection. Scale bar: 20 µm.

### Alignment of the color channels

Mount a slide with fluorescently labeled beads and bring it in focus. Close the slit aperture such that the three channels have equal and maximum width without overlapping (*i.e.*, each channel occupies a third of the full width). Place an almost entirely closed iris after the tube lens and center it on the optical axis. Furthermore, determine the center of the camera chip on the live stream image of the camera (*x*: 1024px, *y*: 1024px), and place the mouse cursor there as a marker. The center of the image formed on the camera through the middle channel should align with the cursors position. Opening and closing the iris helps to figure out the center of the image. With the middle channel’s last mirror, bring the center of the image to the marked cursor position. Then, align the other two mirrors of the other channels in the same way, moving the cursors position correspondingly ((*x*: 341px, *y*: 1024px) and (*x*: 1705px, *y*: 1024px)). The next step is the alignment of the focal position for all three color channels. In the previous axial alignment step, a quasi-infinite distance object was imaged to make sure that the camera is placed in the focal point of the second 4f-lens of the middle channel, so that the focused image of the sample is projected on the camera chip without further magnification. Therefore, the middle channel is used as a reference to align the other two channels. Adjust the axial position of the lenses of the left and right channel so that the beads appear in focus in every channel. It helps to move the sample repeatedly up and down a few hundred nanometers, sweeping through the focus position. Either use a second person to move the sample stage up and down, or a script that loops through the positions (see ***Figure SM1***).

Once all channels are co-aligned with respect to their focus position, use the mirrors of the left and right color channel to precisely align the images laterally. Make sure that the bead sample stays in focus during the entire lateral alignment, and if necessary, correct the stage position. The color channels will be co-registered computationally with subpixel precision during the data analysis steps, but aligning the color channels optically helps to maximize the shared usable field of view and is convenient when viewing the raw data before the analysis steps (see *Figure S47*).

Our K2 microscope software has the option to superposition the channels with RGB colors live, which makes it easy to align all three color channels with respect to each other. Alternatively, take a screenshot of the middle channel and transparently overlay it with the other two channels, using *e.g.* the freeware Ghostit [41]. Typically, we achieve residual shifts of less than 0.5 µm by manual alignment (see *Figure 11*d).

**Figure S27.**
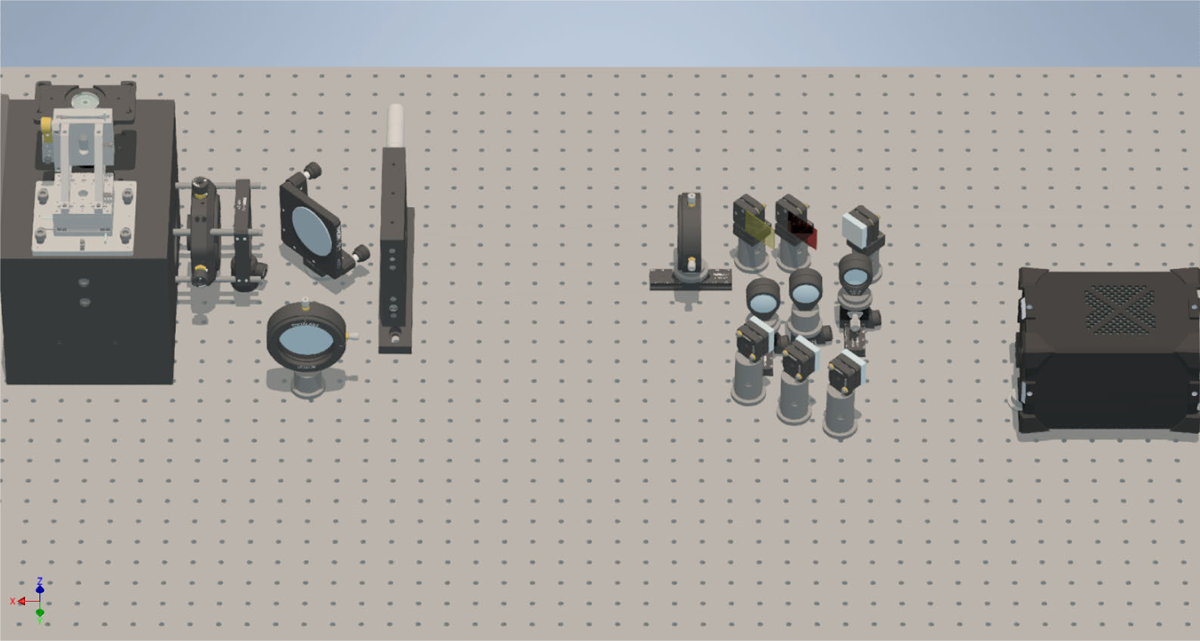
Installing the first mirror and lens of the single-channel path The three-channel detection path can be bypassed using two mirrors placed on magnetic kinematic mounts. They are installed after the tube lens and before the camera, and use a simple 4f-system as an optical relay. The single-channel path is most conveniently installed alongside the three-channel path, using the Gustafsson alignment tool, but in principle it can also be installed at a later point. Assemble the kinematic bases on pedestal mounts and the kinematic tops on the mirrors. Two additional kinematic bases should be placed somewhere on the optical bench where the mirrors can be stored when not used. Place the first mirror mount just after the tube lens cage assembly. Using the alignment laser, orient the mirror such that the laser beam meets the center of an iris mounted at the desired height of the optical axis and runs along a row of breadboard holes. Use the speckle method to determine the focus position, and place the first lens at a distance of one focal length away from the focal plane. Align the lens using the back-reflection method and the previously placed iris, and confirm collimation.

**Figure S28.**
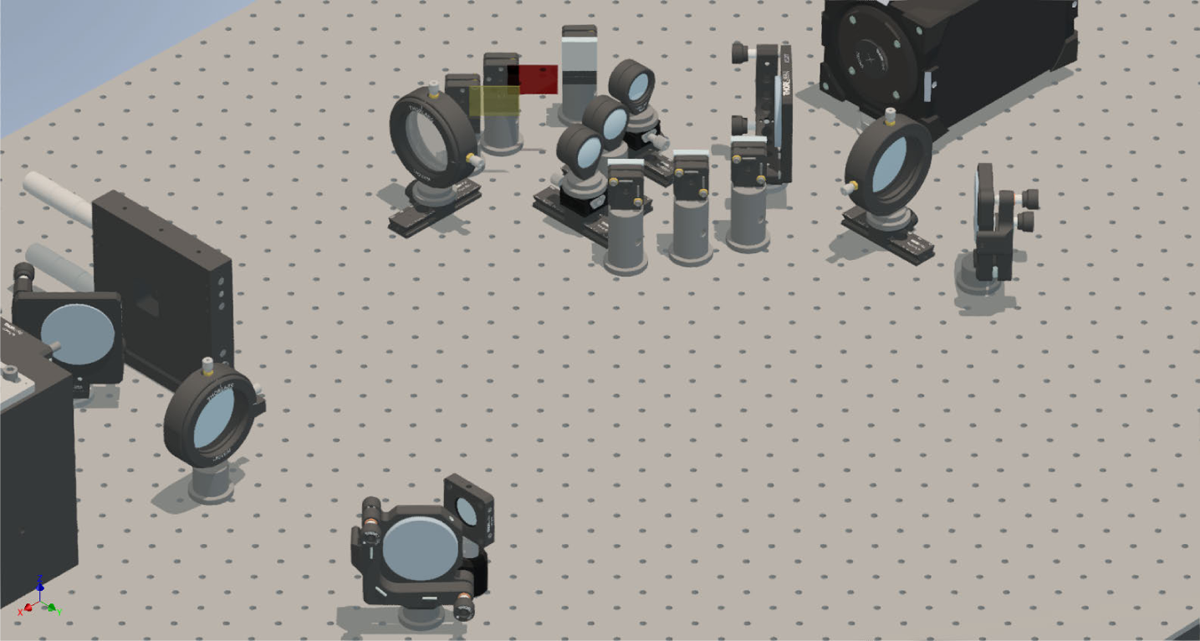
Installing the second mirror, filter and third mirror Place a second mirror and align it with the help of irises mounted along one row of breadboard holes on the optical bench. Any additional emission filters should be placed in the 4f-system as close to the fourier plane as possible, which is located at a distance of one focal length from the first 4f lens. Place the next mirror and align it with the help of irises mounted along one row of breadboard holes on the optical bench. Mount an alignment disk onto the C-mount of the camera, place the final mirror in front of the camera and align it so that the laser beam hits the center of the mirror and reflects it along a row of breadboard holds towards the center of the alignment disk. Then, mount the lens in between the last two mirrors on a rail, roughly one focal length away from the camera. Align the lens using the back-reflection method and verify that the laser beam still hits the center of the alignment disk.

**Figure S29.**
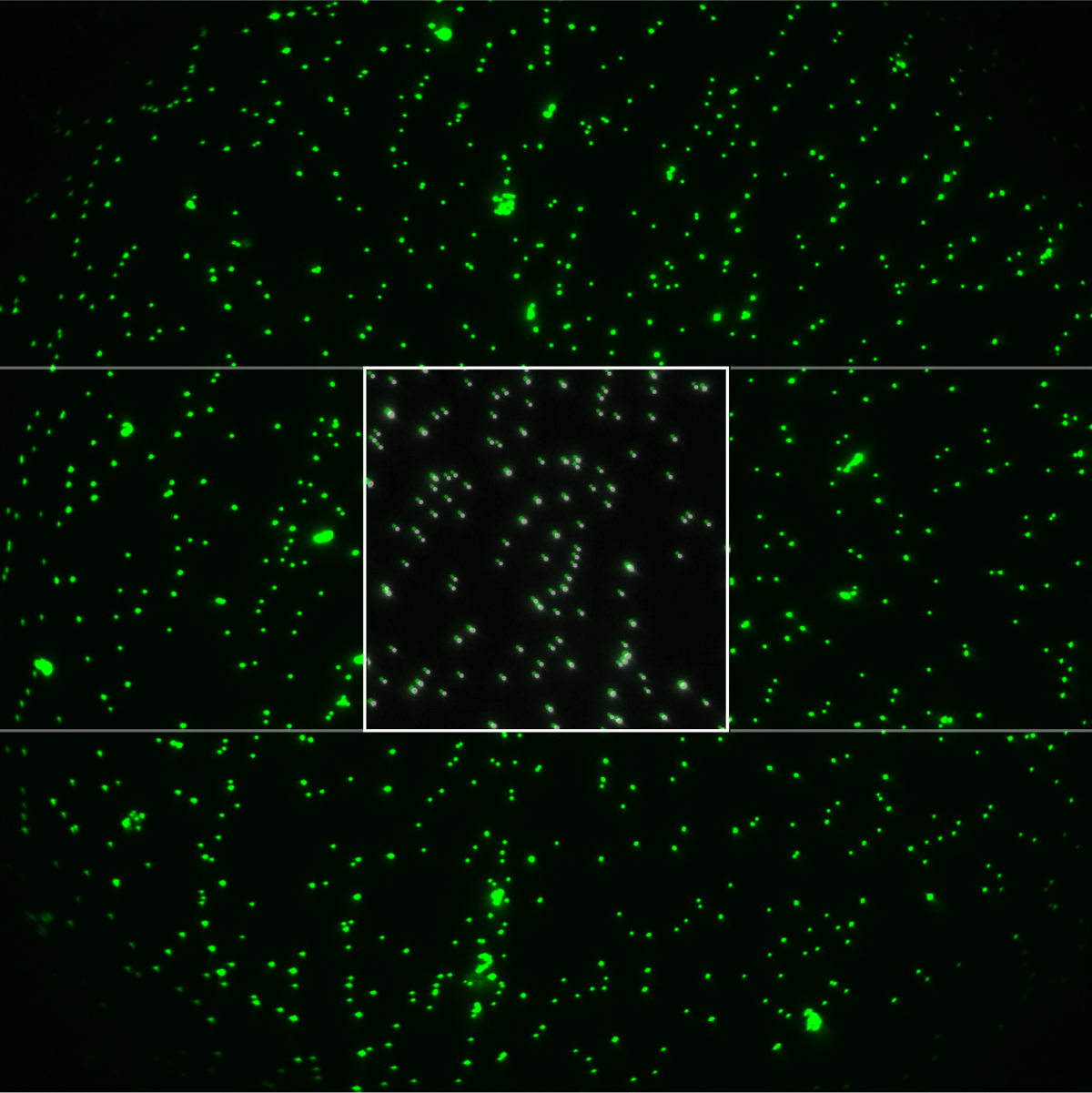
Bead sample imaged using the single-color pathway (green look-up-table). The white square in the middle denotes the overlay of the beads imaged with the single-color pathway, and the beads imaged in the central channel of the triple-color pathway (gray look-up-table). The left and right channels of the triple-color pathway are blocked.

### Co-aligning the single-color with respect to the three-color pathway

For the fine-adjustment of the axial position of the lens, first remove the mirrors on the magnetic mounts, mount the objective and a fluorescent bead sample, and focus on the beads using the already aligned three-channel pathway. Save an image as a reference for the alignment of the single-channel path. Then, place back the single-color pathway mirrors and move the final lens of the single-color pathway axially on the rail until the bead sample appears in focus again. Use the freeware Ghostit [41] to overlay the reference picture, and align the final mirror of the single-color pathway until the central region of the beads colocalize with the reference image of the beads from three-channel path.

### qgFocus path

**Figure S30.**
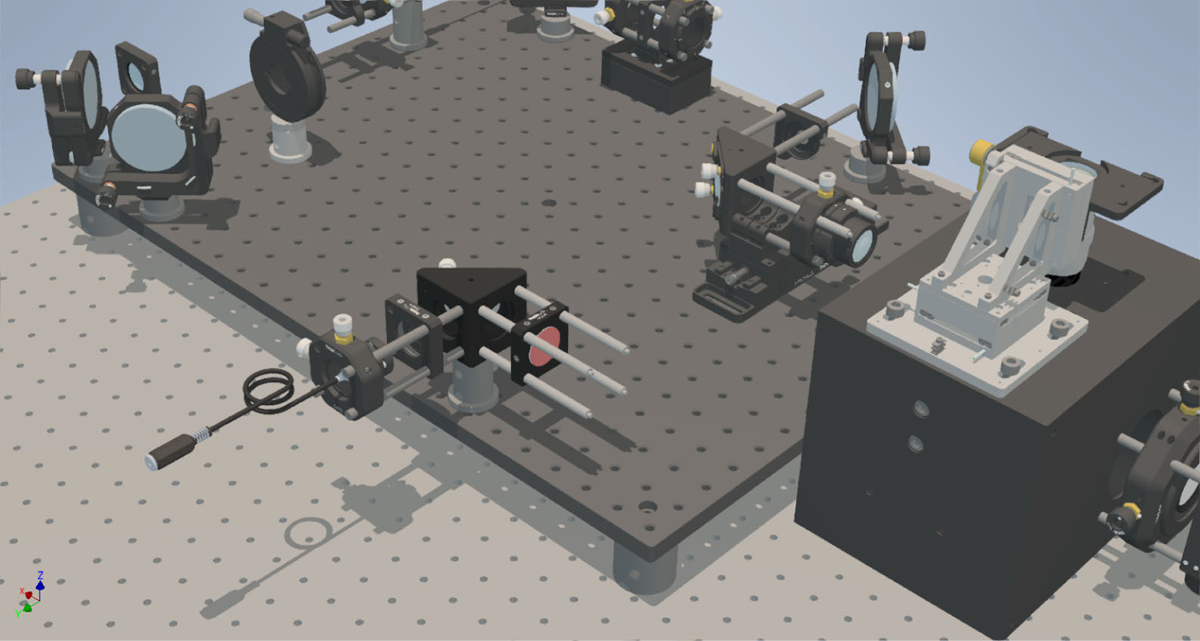
Setting up the qgFocus laser Start with assembling the first part of the cage system by screwing cage rods onto a right-angle mirror mount. Slide a zero-aperture iris on the cage rods and secure the infrared laser in a tip/tilt mount, which itself is mounted in a *x-y*-translation mount. Attach cage rods also on the other side of the mirror mount and mount the entire assembly to the breadboard. Close the iris almost completely and center the infrared laser on the iris using the *x-y* translation mount by viewing it on an infrared viewing card after the almost fully closed iris. Angular alignment is done using the tip/tilt mount and the right-angle mirror while viewing the laser beam on an infrared reticle in a cage mount placed after the right-angle mirror.

**Figure S31.**
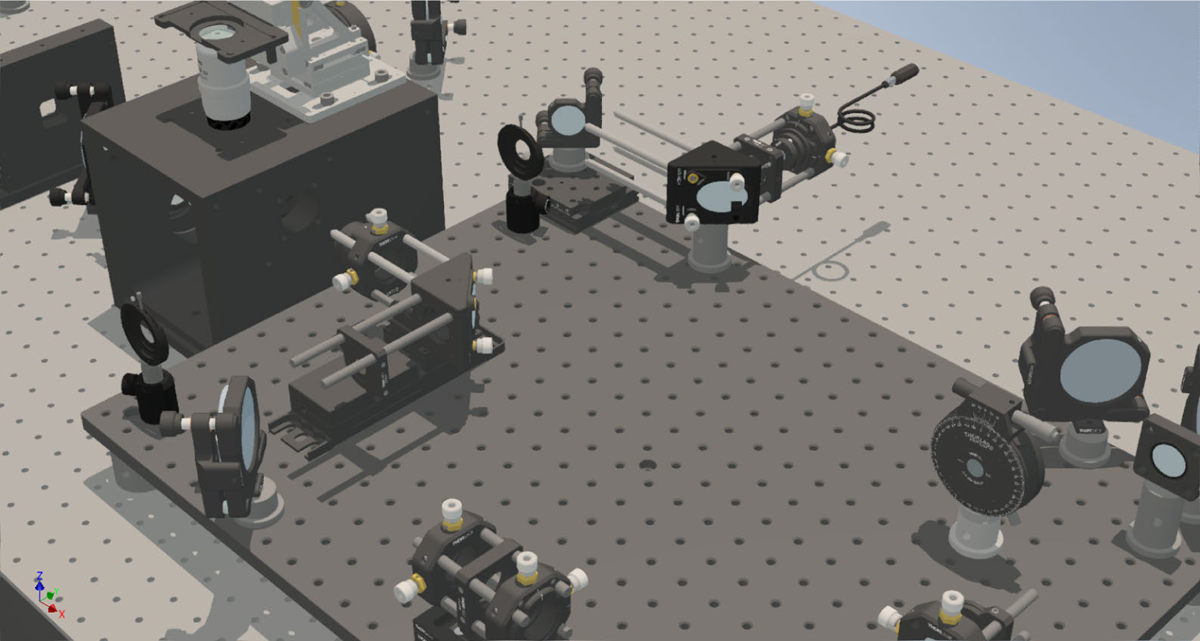
Mounting the dovetail stage Mount the dovetail stage on the breadboard and place a mirror on the stage such that the laser beam roughly hits the center of the mirror on the translation stage. Mount two irises, one far and one distant from the stage, in one row of breadboard holes. Verify that the center of the irises match the beam height of 65 mm. Adjust the dovetail stage and the mirror on the dovetail stage so that the laser beam passes both irises.

**Figure S32.**
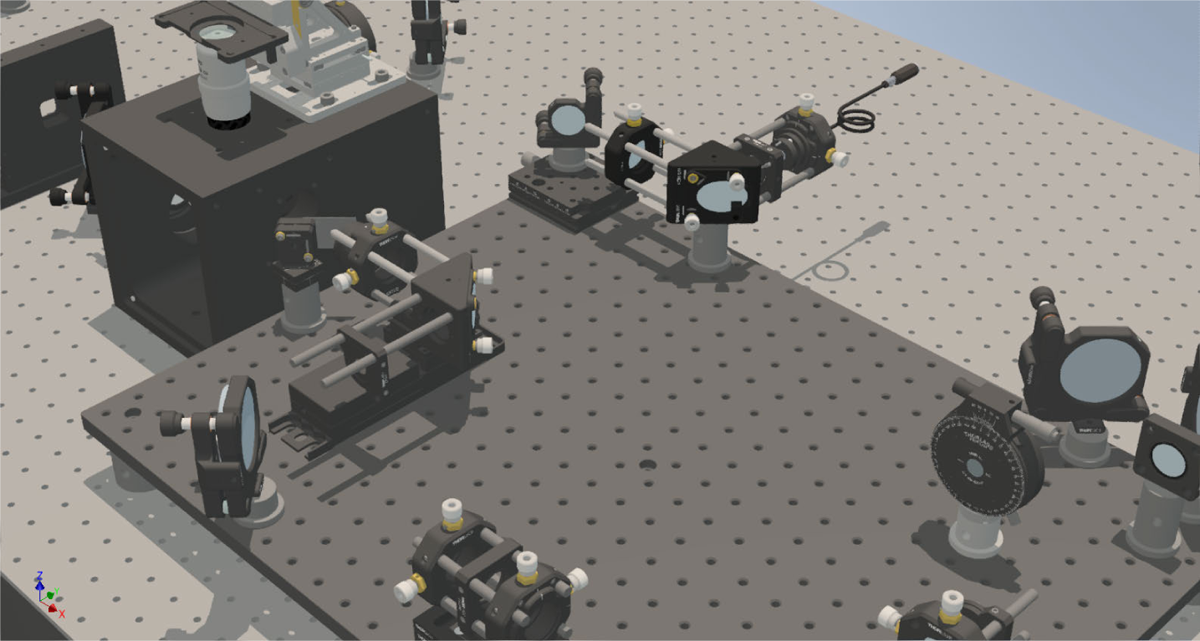
Coupling the infrared laser into the excitation path Assemble the dichroic coupling mirror by mounting the mirror mount onto the magnetic base, and glueing the dichroic mirror onto the mirror mount with two-component silicone glue. Place the infrared coupling dichroic mirror in between the central cube and the TIR focusing lens. Align the dichroic mirror by following the back-reflection of the excitation lasers, which should roughly travel along one row of breadboard holes at unchanged height. Furthermore, make sure that the excitation lasers are not clipped throughout the whole TIR to epifluorescence movement range of the TIR angle stage. Move the excitation lasers into epifluorescence mode, and, using the dovetail stage for translating the infrared laser beam, ensure that the infrared beam coincides with the excitation laser’s spot on the coupling dichroic mirror.

**Figure S33.**
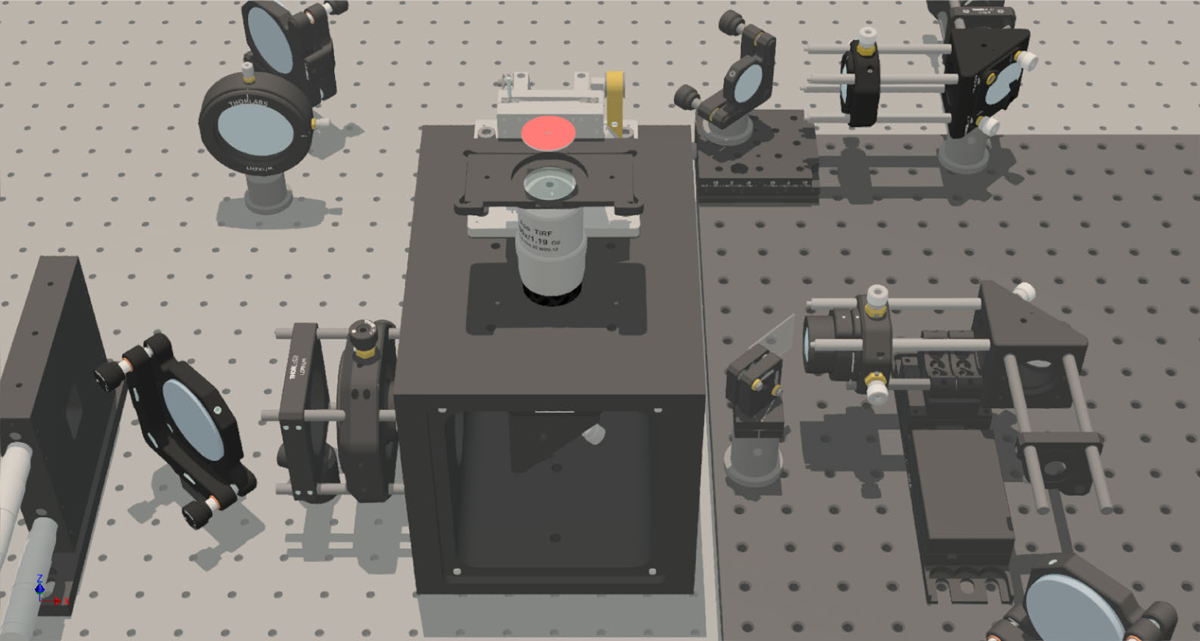
Collimation of the infrared laser beam after the objective *Warning: The following steps need to be done very carefully with the appropriate laser goggles, since the collimated laser beam can potentially hit and harm the eye.* Mount the TIR focusing lens in a *xy*-translation cage mount and slide it onto the infrared laser cage system. Use the translation mount to align the lens so that the excitation and the infrared laser beams remain coincident on the coupling dichroic mirror. Mount a glass slide on the objective with immersion oil and choose the sample stage *Z*-position where the sample would approximately be in focus (this can be determined quickly by placing a sample with beads or dye in solution, switching the excitation lasers to TIRF mode and noting down the stage position where the sample is in focus). Use an infrared viewing disk to follow the excitation laser beam exiting the objective. For checking the co-alignment of the (epifluorescence) excitation laser beam and the infrared laser beam, a convenient approach is to use a very dim excitation beam - higher intensities will quickly saturate the infrared reticle. Intermittently block the excitation beam, then realign the infrared laser using the coupling dichroic mirror, and go back to comparing it to the excitation beam again at different positions above the objective. The axial alignment of the lens focusing the infrared beam onto the back-focal plane should be such that the infrared laser beam is collimated after the objective. The long focal length of this lens allows for some tolerance in the exact axial position. In practice, an infrared beam that is still detectable with the infrared viewing card far away from the objective (*e.g.* the ceiling) is sufficiently collimated.

**Figure S34.**
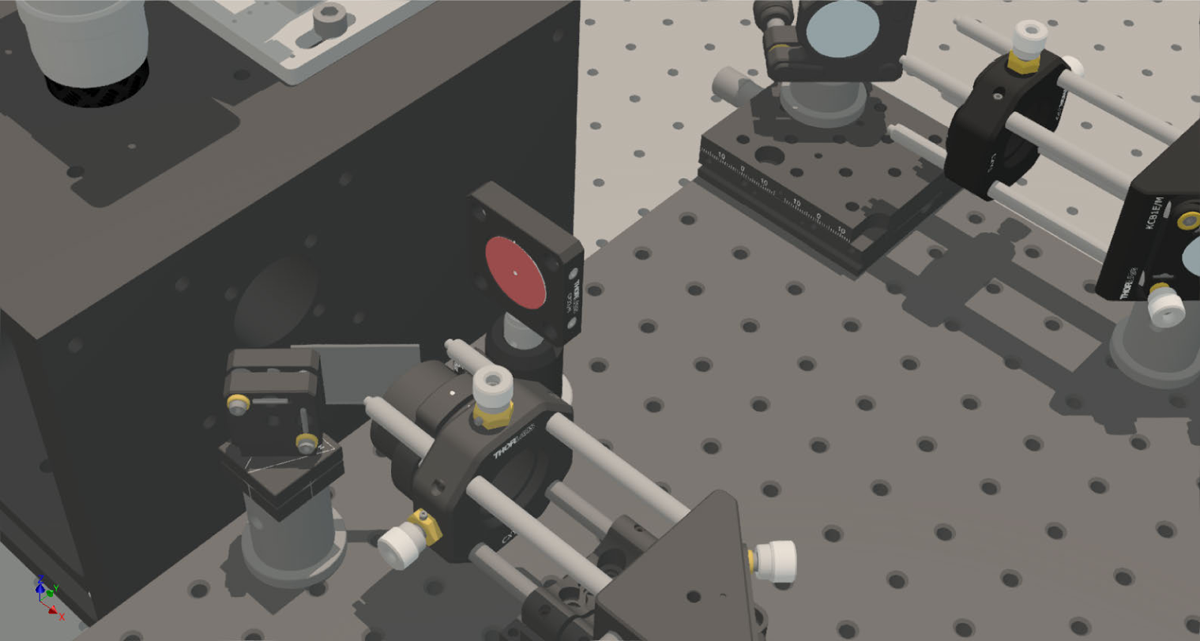
Directing the infrared laser beam into total internal reflection Use the translation stage to shift the infrared laser beam by about 4.5 to 5 mm in negative *x*-direction (*i.e.* towards the direction of the detection path). The transition towards shallower angles can be observed with a viewing card close to the objective. Place an infrared viewing disk with a center hole between the dovetail stage and the coupling dichroic mirror with the laser beam passing through the hole. The back-reflected beam should be visible on the disk, and it’s *x-y* movement resulting from moving the sample in *Z* can be observed (see supplementary movie SM5). This can be done for different stage positions (the reticle has to be moved to avoid obstructing the incoming laser beam), and eventually the intensity of the back-reflected beam will increase noticeably upon total internal reflection. Once total internal reflection of the infrared laser beam is established, add a drop of water onto the microscopy slide. Starting out with a glass-air interface facilitates the initial alignment, but further translation may be necessary since the critical angle for a glass-water interface is higher than for glass-air.

**Figure S35.**
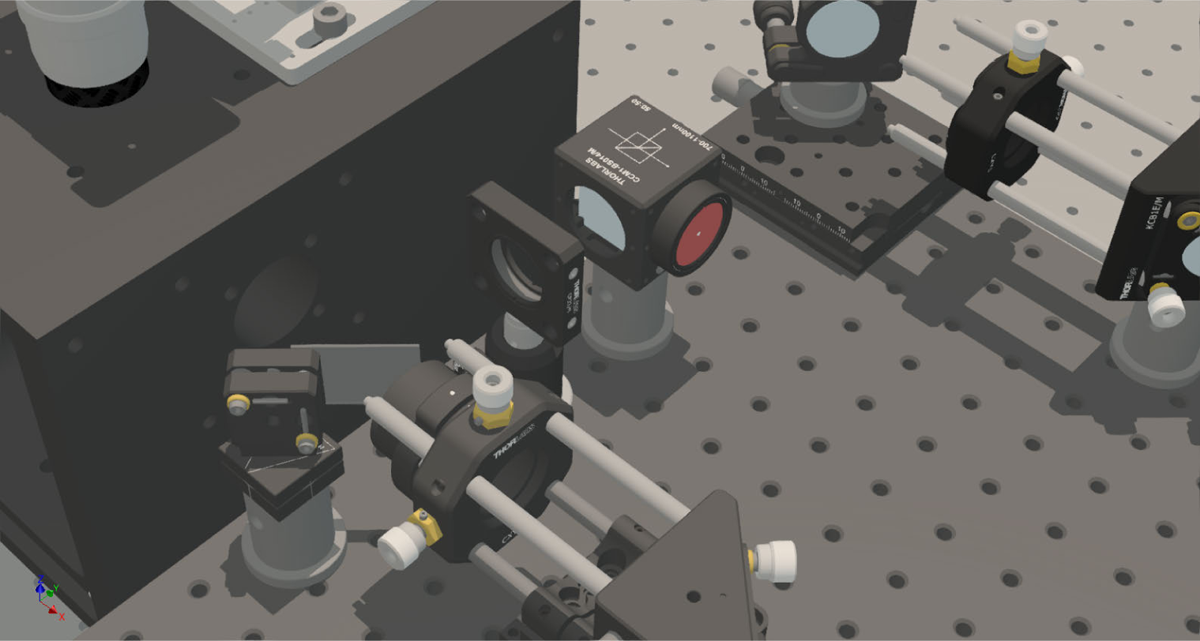
Assembling the qgFocus detection path Place a sample of fluorescent beads or dye in solution, switch on the excitation lasers in TIRF mode and find the focus position. Verify that the back-reflection of the infrared laser is still visible on the infrared viewing disk. Then, mount the infrared viewing disk on the beamsplitter cube and place the beamsplitter cube such that the back-reflection of the infrared laser hits the cube’s center, and is reflected towards positive *x*-direction. Placing the beamsplitter cube centered on the back-reflection ensures that the infrared laser beam is not clipped prematurely by the beamsplitter aperture when moving through big *Z*-ranges with the sample.

**Figure S36.**
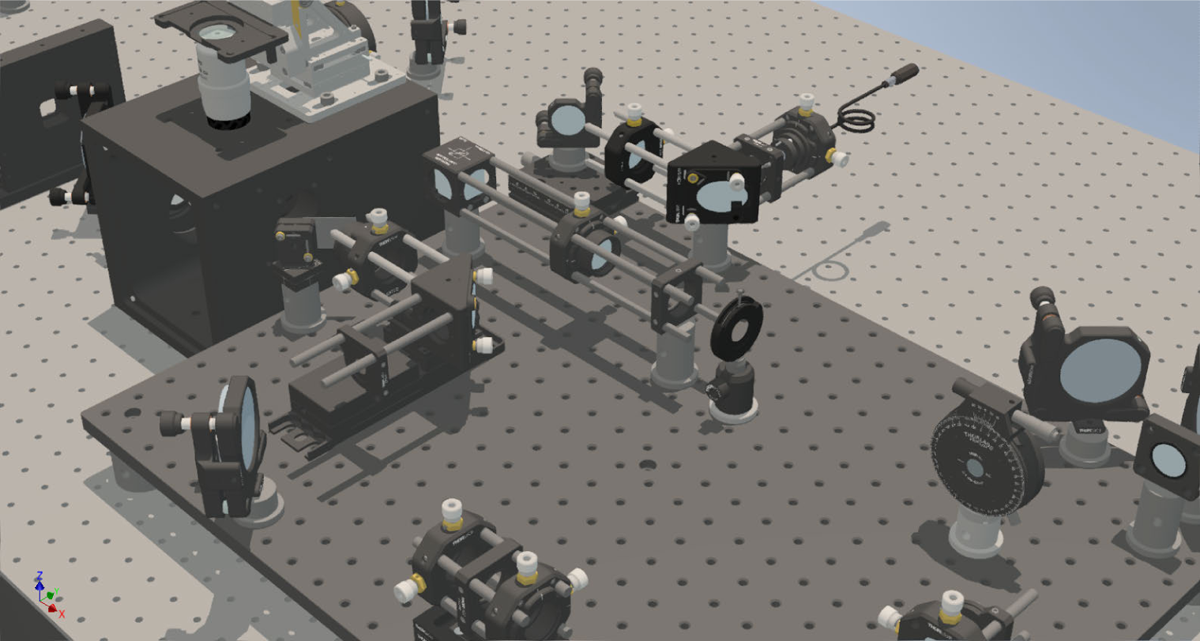
Beamsplitter cage system and focusing lens Use an iris to mark the position of the reflected laser beam after the beam splitter, mount the cage rods onto the beam splitting cube and slide on a translating lens mount with the lens that focuses the infrared laser beam onto the CCD array. Align the lens such that the laser beam remains centered on the iris when moving the lens axially.

**Figure S37.**
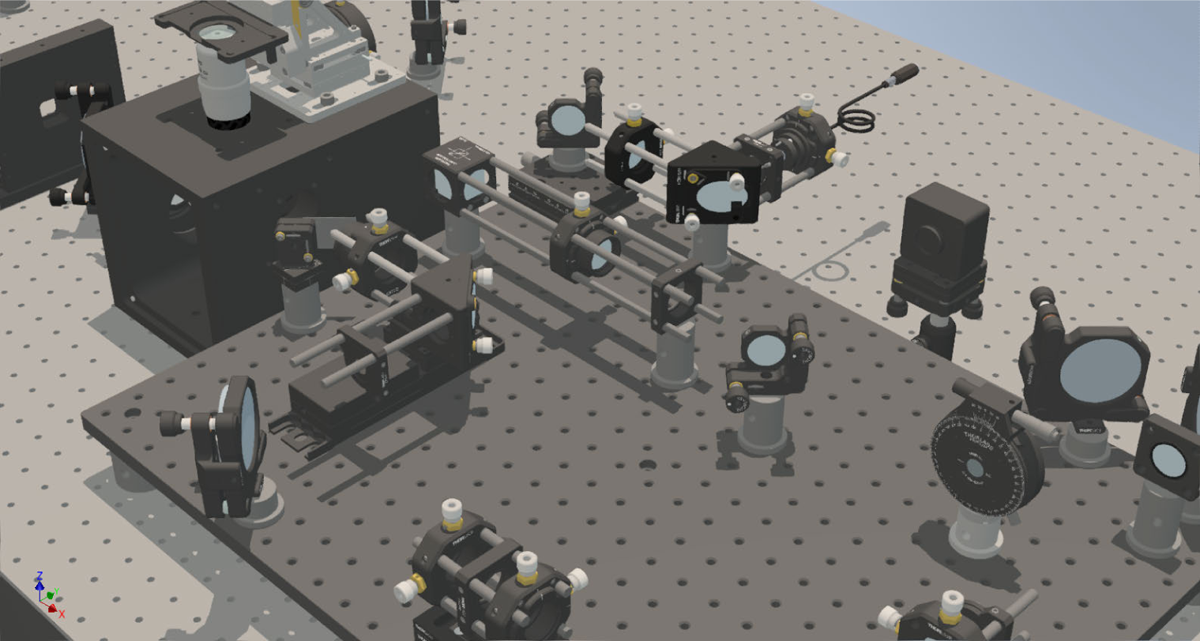
Installing the CCD array Mount a mirror after the cage system to simplify aiming the laser beam onto the CCD array. Place the CCD array about 200 mm after the lens and use the mirror to guide the infrared laser beam onto the CCD array. View the beam profile on the CCD array on the computer and try to get a sharp peak by adjusting the last mirror. If the CCD array is saturated by the infrared laser, it may be necessary to prepone installing the neutral density filter mentioned in the next step. Adjust the distance between the CCD array and the lens such that the central peak spans at least 10 pixels.

**Figure S38.**
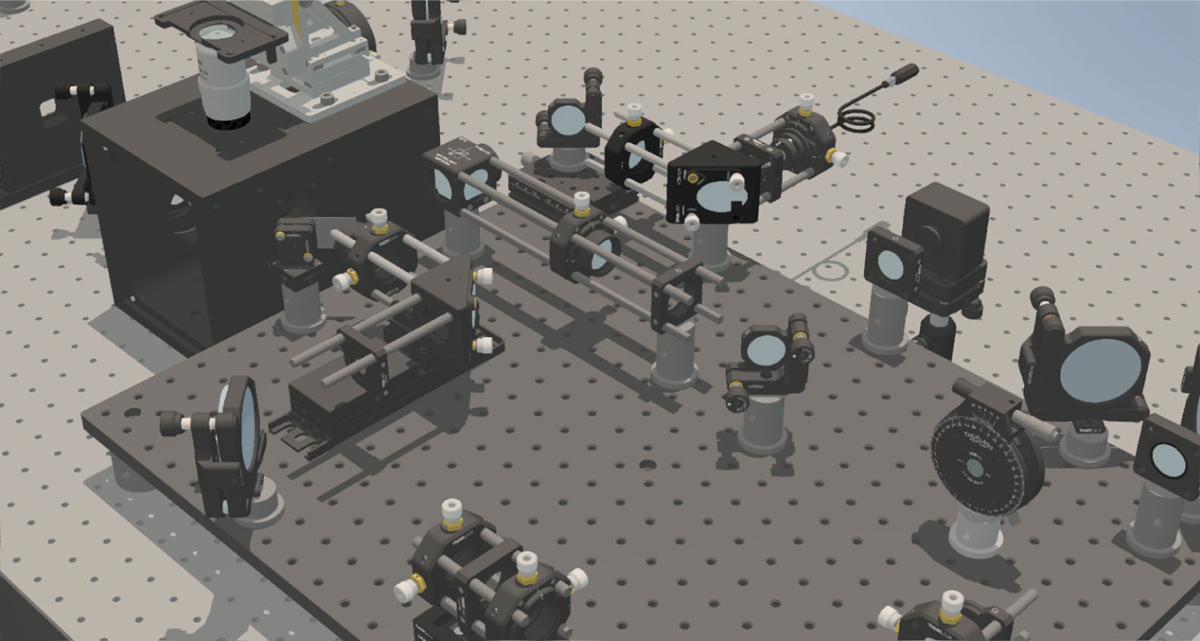
Installing bandpass and neutral density filters Install a long-pass filter just before the CCD array to suppress background light. Finally, mount a neutral density filter on the beam splitter to reduce the infrared laser intensity, preventing it from interfering with measurements. The exposure time of the CCD array may need to be adjusted, to compensate for the lower intensity.

### Additional elements

**Figure S39.**
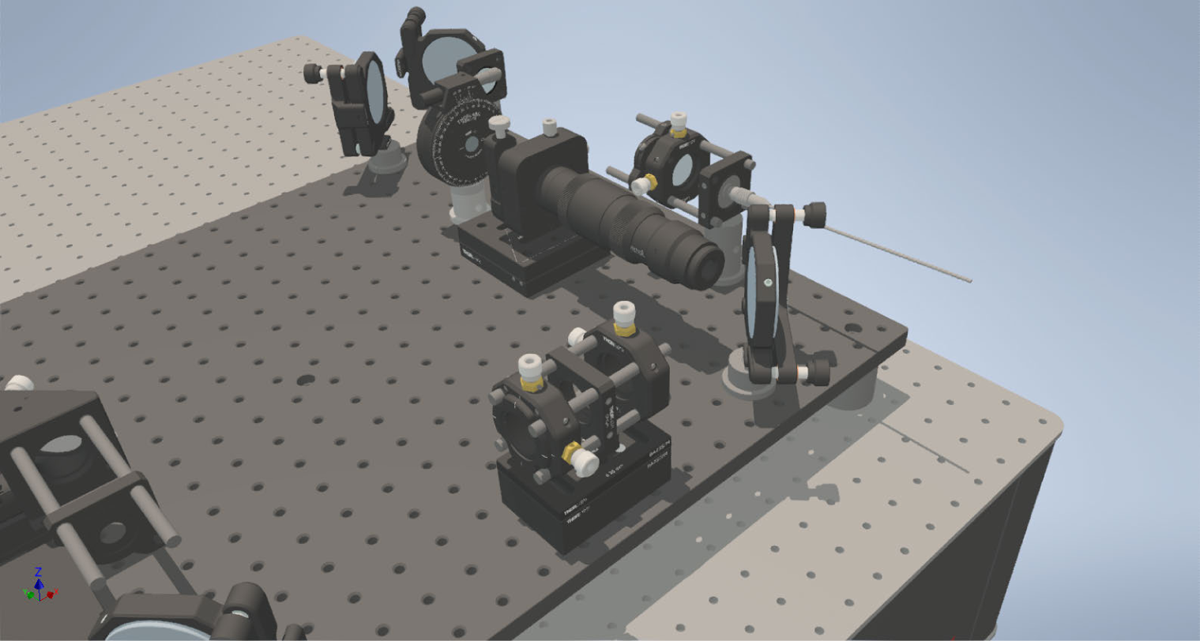
Alignment of the refractive beamshaper Mark the position of the excitation laser beam further downstream of the supposed beamshaper placement for future reference, using an iris or a viewing disk. Assemble the kinematic base on the breadboard, and the kinematic top on the beamshaper mount. Mount the aluminum alignment piece on the beamshaper mount and place it such that the laser beam moves through the device without obstruction. Follow the manufacturer’s manual for alignment. In brief, place the biggest orifice on the alignment tool and use tilt/tip and *x-y* translation to center it onto the optical axis. Repeat with decreasing orifice size and verify that the laser beam stays on the marked spot with the aluminum rod inserted. Then, exchange the aluminum rod for the actual beamshaper. Image a sample of dye in solution and carefully correct for any remaining inhomogeneities of the illumination field.

**Figure S40.**
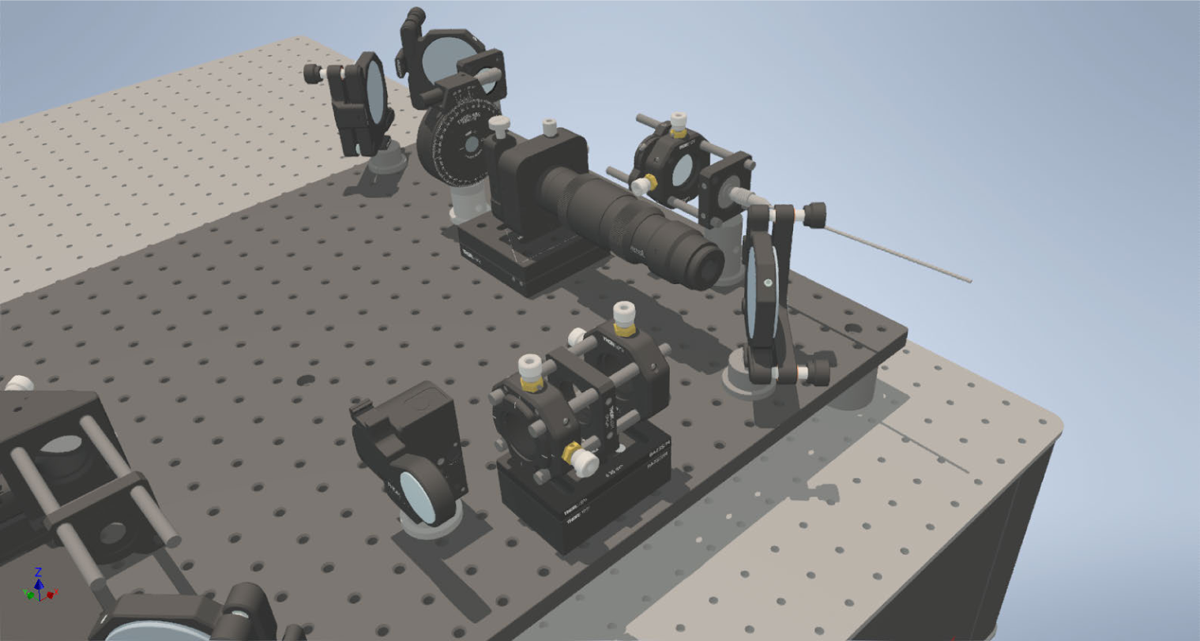
FRAP lens Place the lens in the flip-in mount, and install it on the breadboard using a 12.5 mm pedestal post with 2.5 mm spacers to reach 65 mm beam height. Connect the flip-in mount to the power supply and to the computer. While imaging a sample with dye in solution, flip the lens in and move the mount such that the bleaching spot is at the desired position, uniform, and has the desired size (see Figure 15). Secure the components on the breadboard and flip the lens out again.

**Figure S41.**
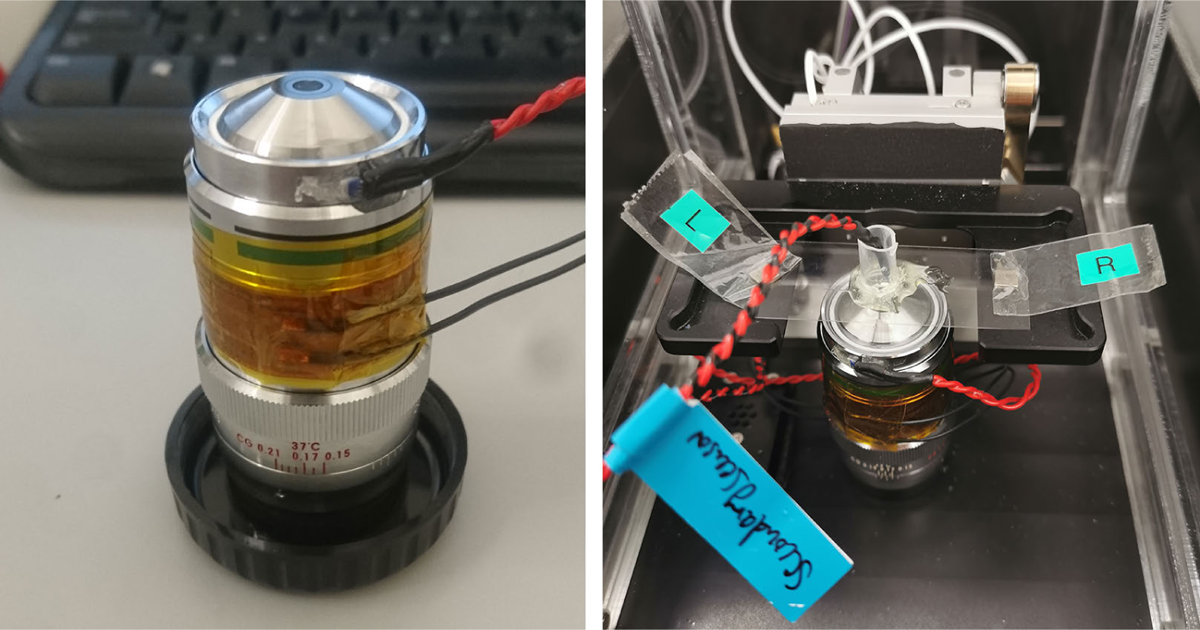
Objective heating Wrap the flexible heater tightly around the objective, such that the correction ring can still move freely, and secure the heater with adhesive Kapton foil. Then, wire up the temperature sensor, prepare two-component heat-resistant epoxy glue and glue the sensor onto the side of the objective, where it does not come into contact with immersion oil (left). A second temperature sensor should be prepared to measure the temperature in the sample (right), to determine the temperature offset between the objective sensor and the sample sensor (in our setup: 5 K at 42 ^◦^C objective temperature). Connect the temperature sensor and the flexible heater to a power source with a PID temperature controller.

**Figure S42.**
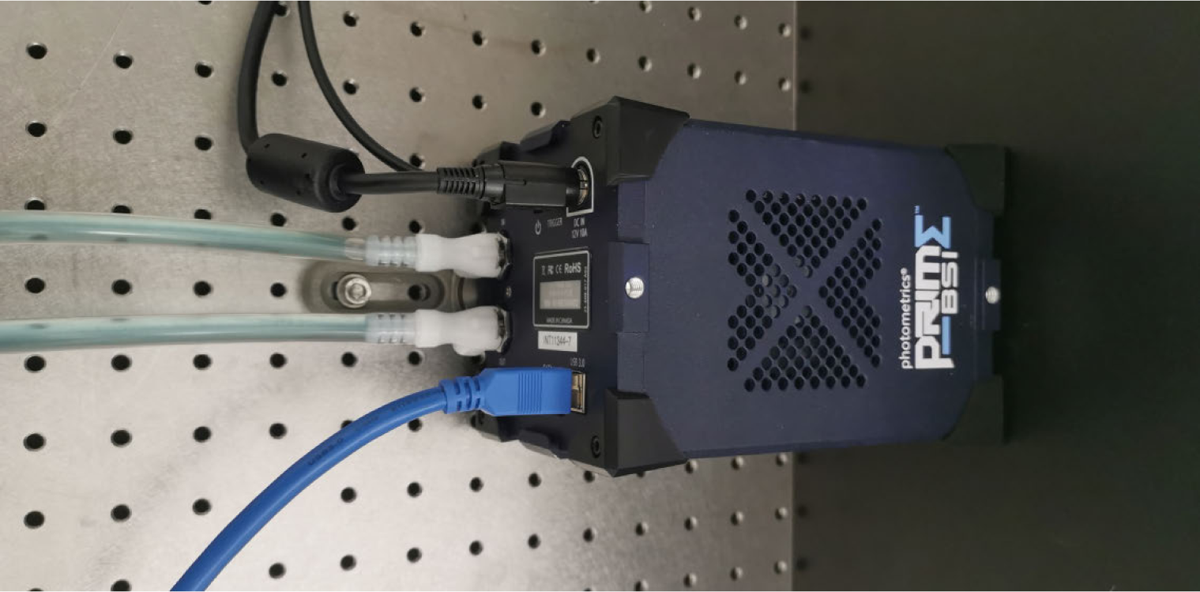
Camera liquid cooling The camera is connected to a liquid cooling system (Eiswand 360 Solo, Alphacool) to minimize vibrations induced due to the cooling fan.

**Figure S43.**
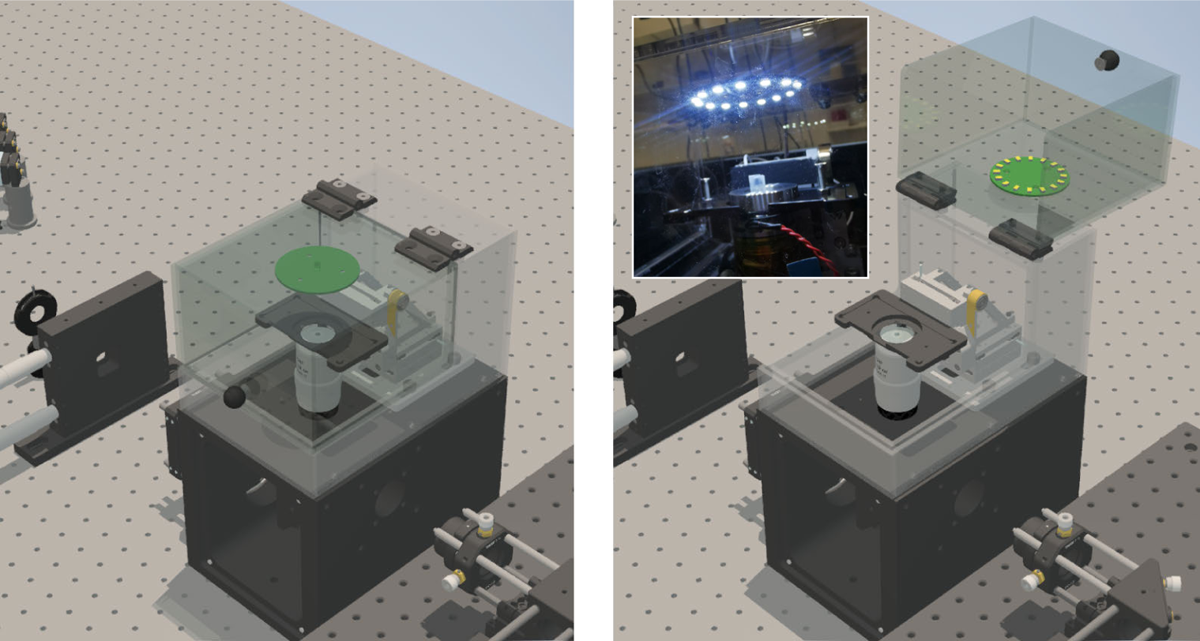
Plexiglass cover and brightfield illumination Install the plexiglass cover and connect the LED ring to the power source. The power source is integrated in a custom box together with the PID temperature controller, but can alternatively also remain a stand-alone part or be powered by a computer USB port. Inset: switched on brightfield LED ring.

**Figure S44.**
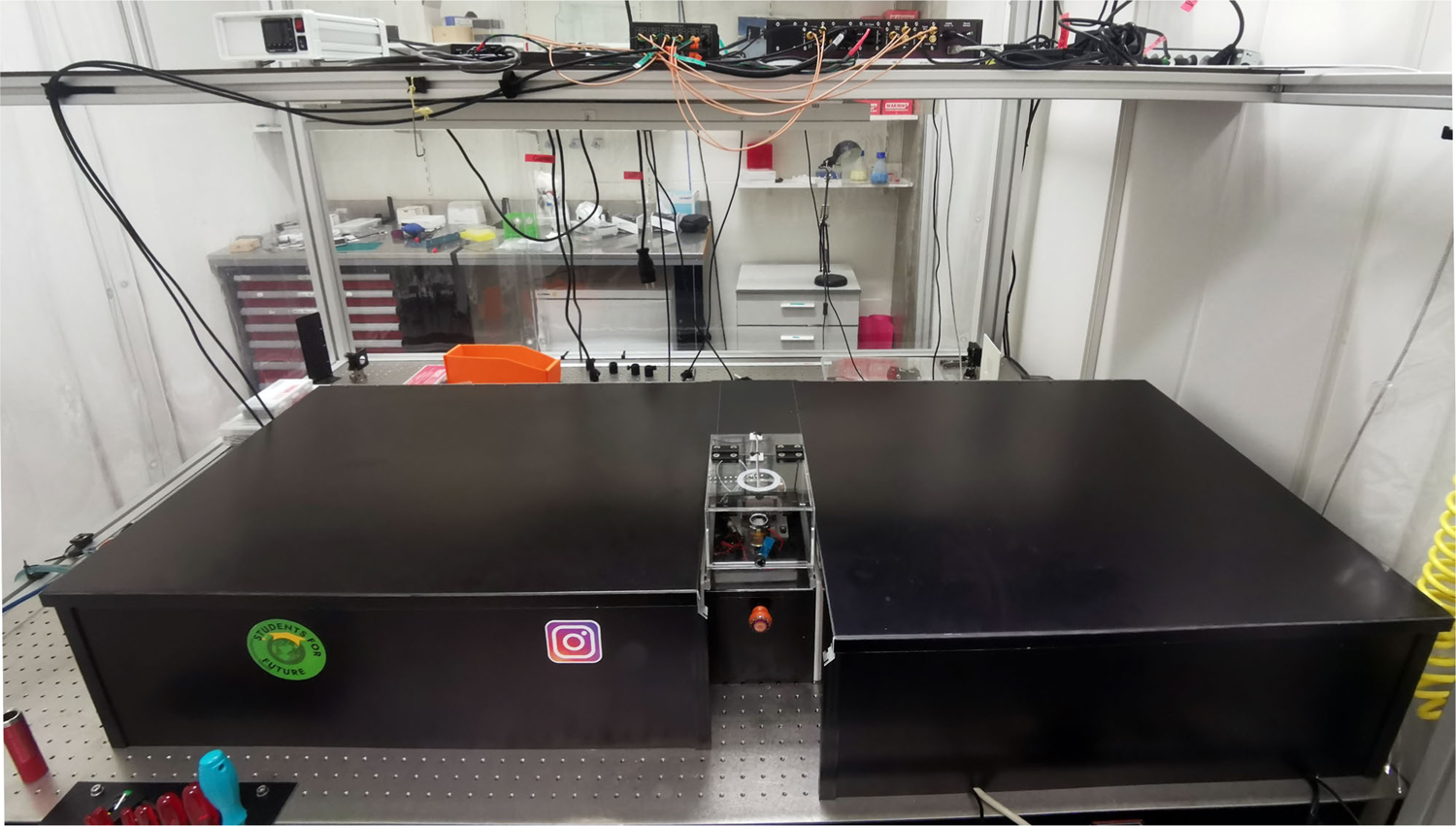
Light-tight aluminum enclosure Light-tight enclosure boxes were constructed from 2 mm thick anodized aluminum and 25 mm square construction rails. Besides minimizing background light levels stemming from the computer screen and the sample stage controller display nearby the setup, it also reduces laser safety issues and dust buildup on the optical components.

**Figure S45.**
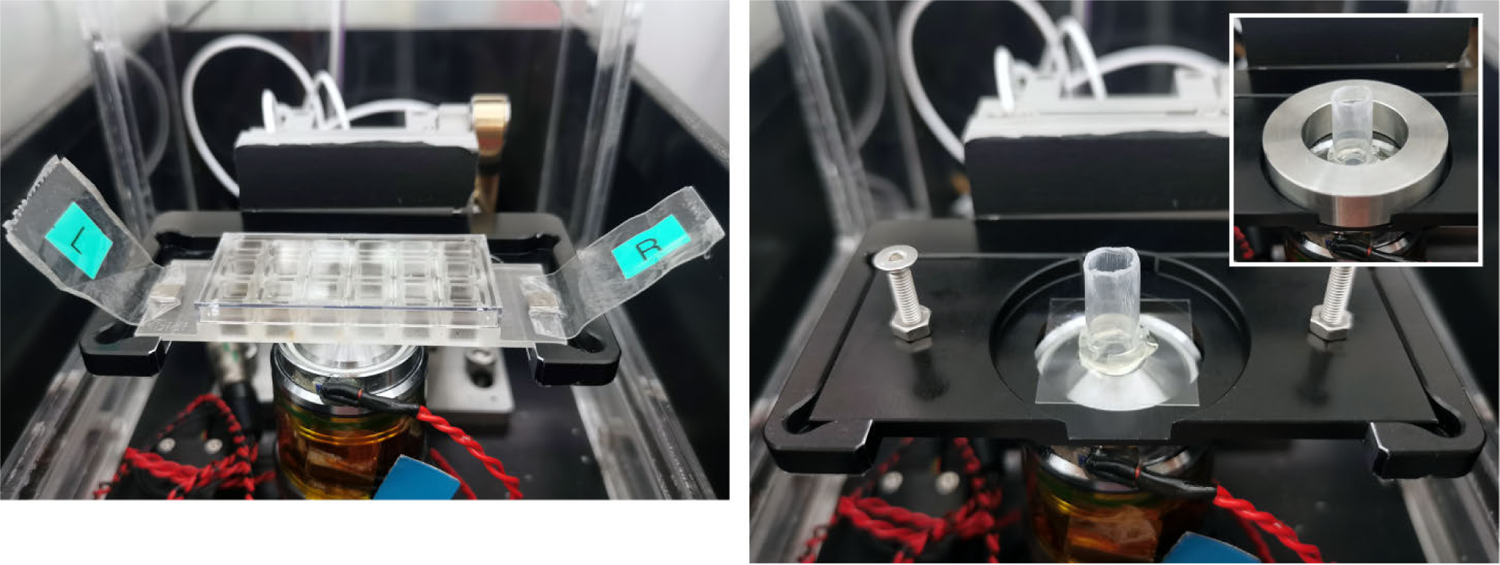
Sample holder Samples with different dimensions are accommodated by a two-part sample holder. Secure mounting is ensured either via magnets (rectangular samples, left) or via a aluminum ring placed on top of the sample (square samples, right).

### Additional K2 software panels

**Figure S46.**
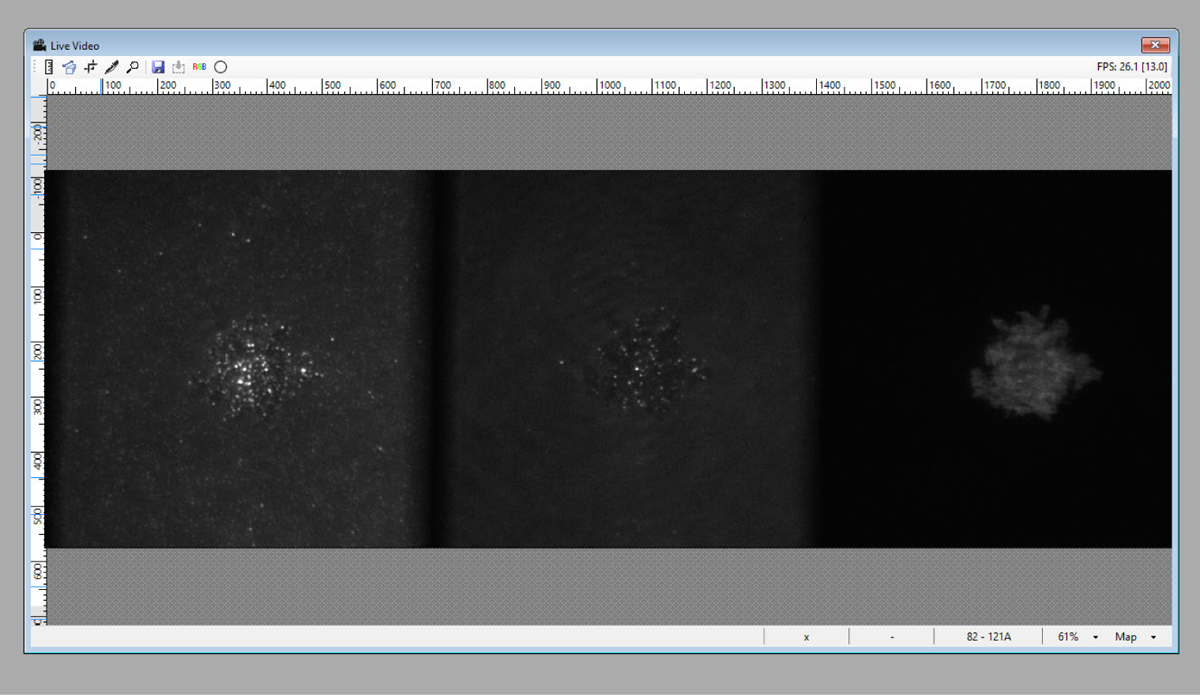
K2 GUI window: Live Video Displays camera live stream with contrast settings either auto-scaled or user-defined. A range of colormaps, a line profile measuring tool and an option to display the FRAP bleaching circle, are available. When operated in triple-color imaging mode, a RGB-viewing mode can be used to view the three channels in superposition (Figure S47).

**Figure S47.**
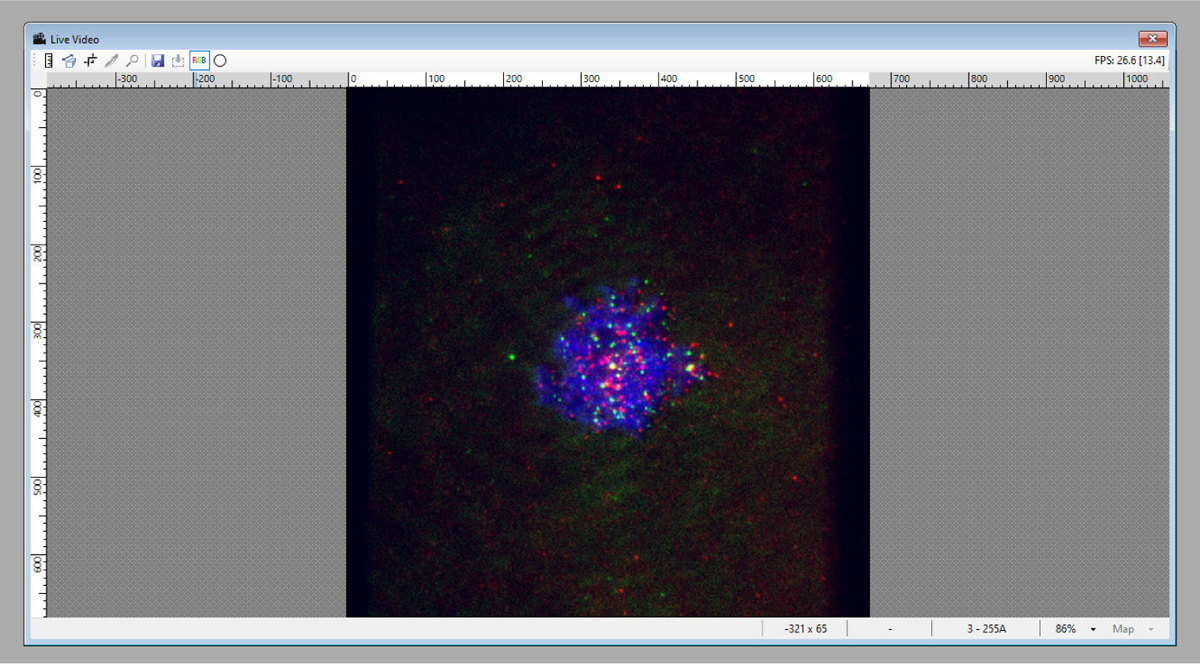
K2 GUI window: Live video - RGB mode Superposition of the three channels in triple-color imaging mode. Contrast settings and spatial offsets are adjustable per channel.

**Figure S48.**
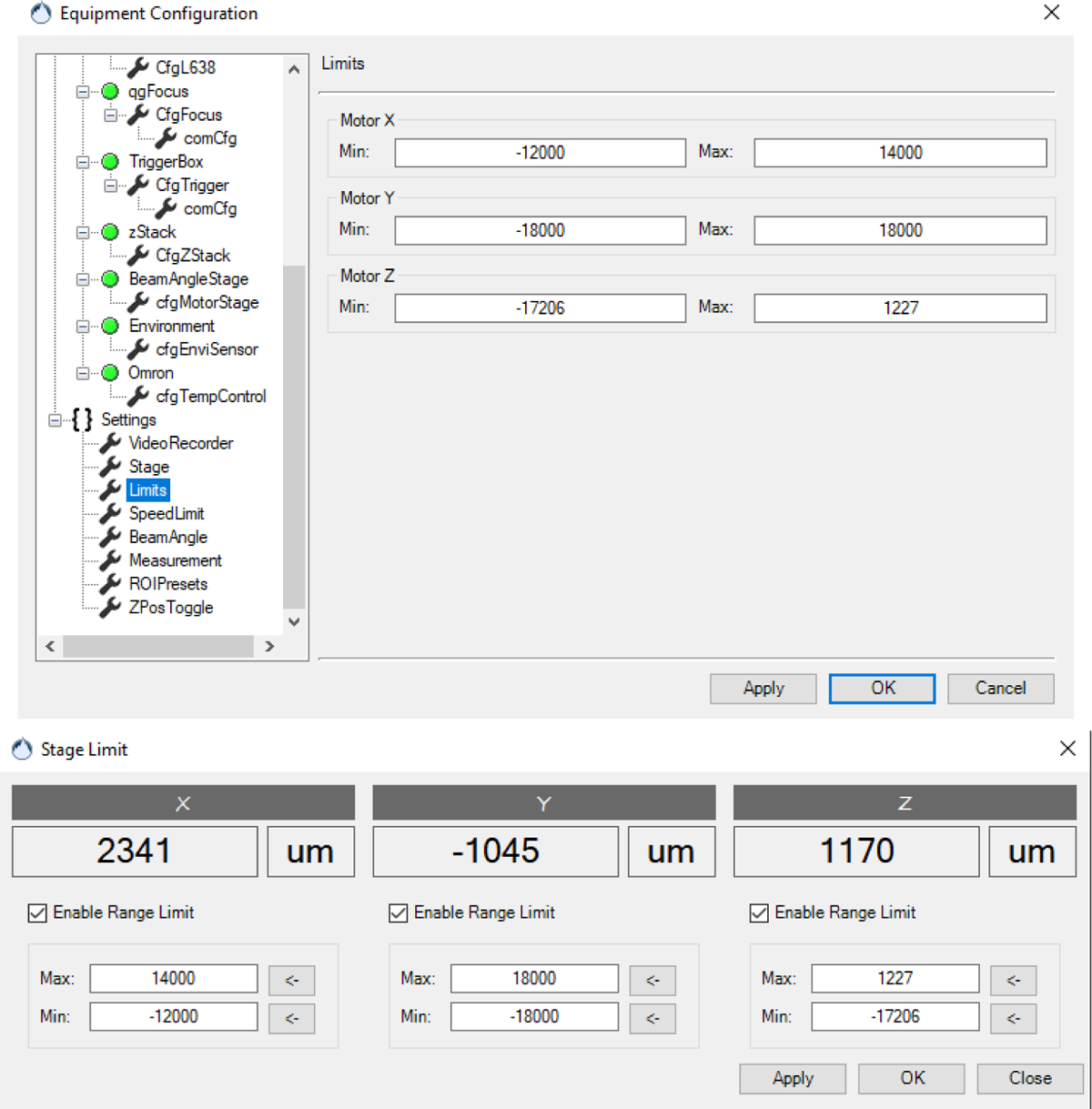
K2 equipment configuration: Stage settings Step distances corresponding to ‘Fine’, ‘Medium’ and ‘Coarse’ and stage position limits are set, preventing the stage from colliding with the objective in case of a user maloperation. However, since the piezo stick-slip stage generates a maximum force of 20 N, physical damages to the objectives are anyway less likely compared to conventional motorized stages.

**Figure S49.**
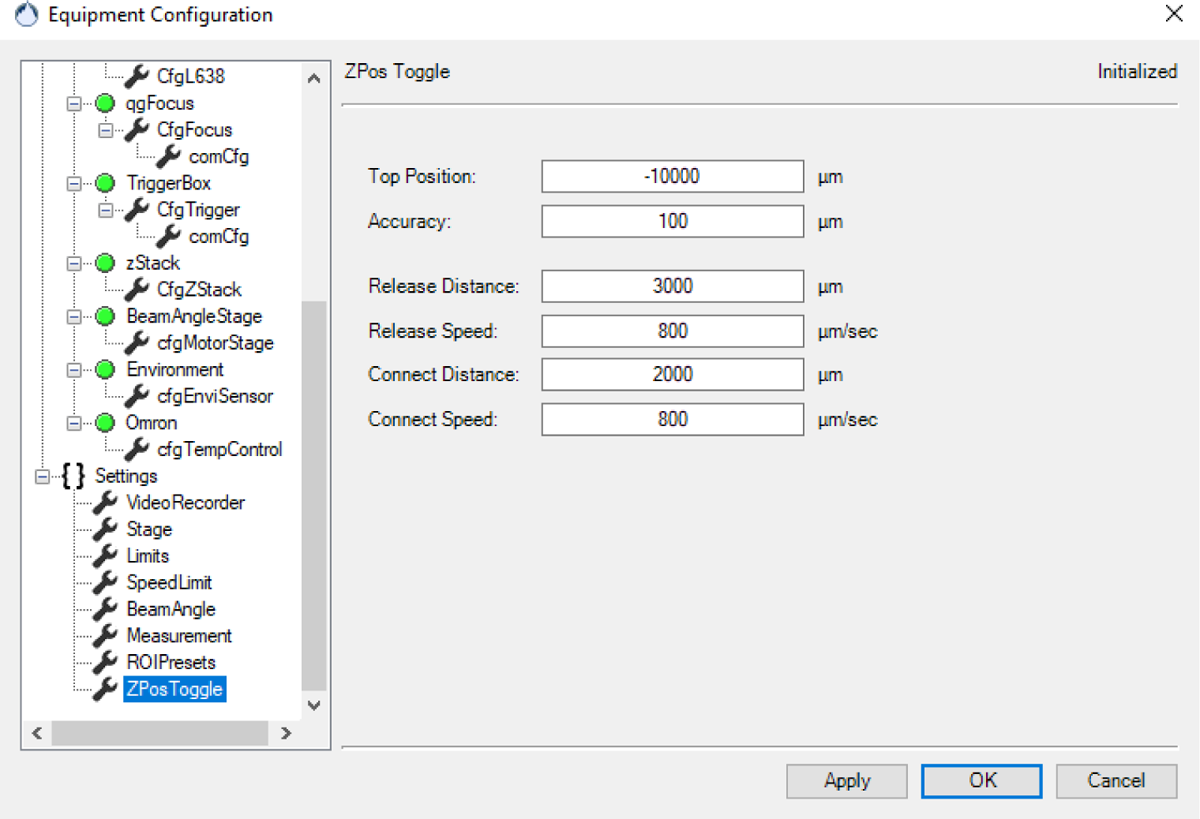
K2 equipment configuration: Z-position toggle settings The Z-position toggle facilitates the exchange of samples, and sample handling during multiposition acquisitions, by moving the sample away from the objective to a user-defined Top Position. During the initial distance (Release Distance) when removing the sample from the objective, and the final distance when approaching the objective (Connect Distance), the sample stage moves slowly at a given Release Speed and Connect Speed. This avoids the introduction of air bubbles into the immersion oil, and gives the immersion oil time to draw back to the objective, therefore reducing the amount of user interaction required to reapply immersion oil.

**Figure S50.**
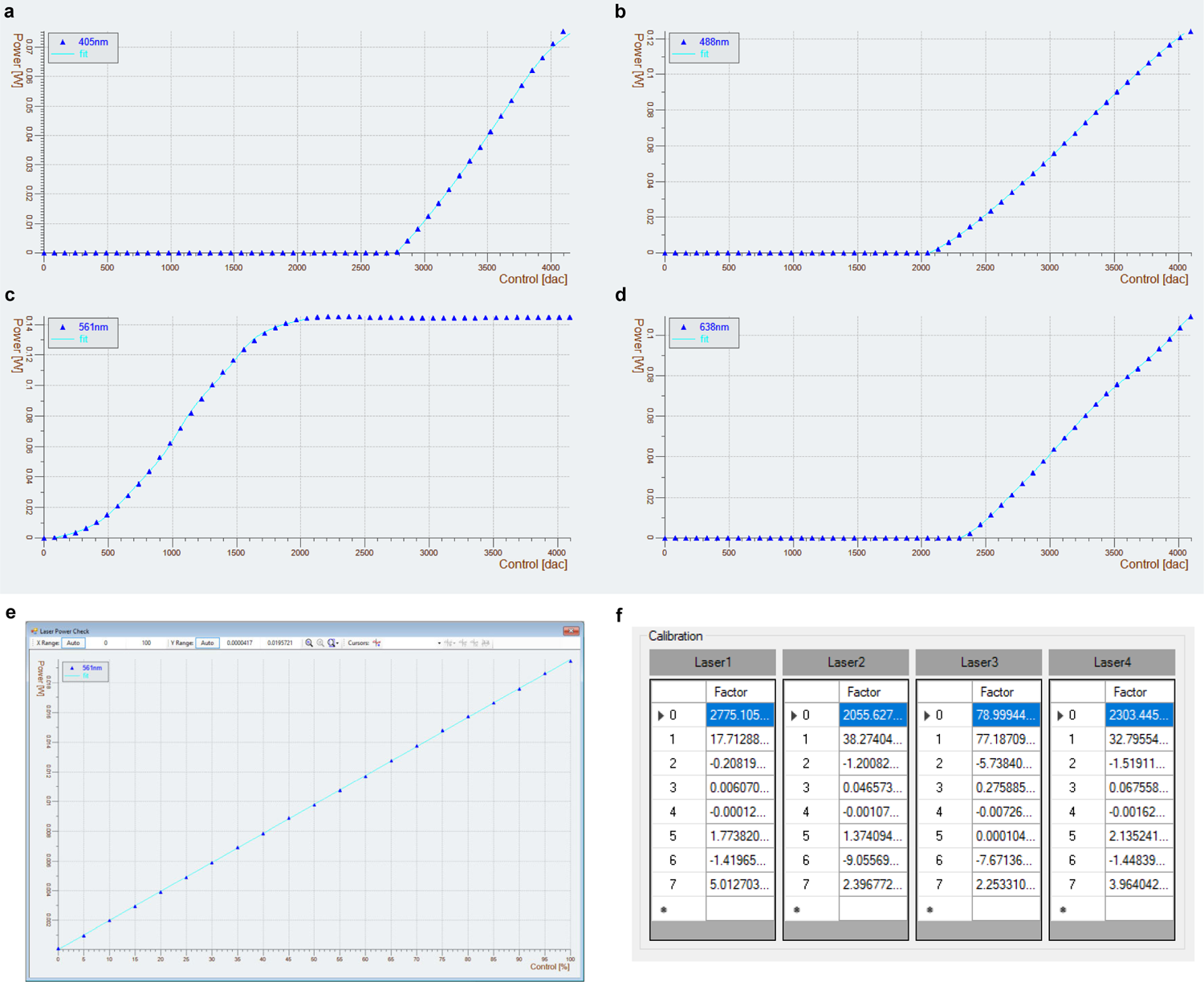
Laser power calibration procedure. a-d) A linear power-to-percentage relation helps optimize the illumination settings during fluorescence experiments and allows the user to have full control and knowledge over the actual laser power used. We developed an automated routine using the K2 software script interface: First, we measure the laser output power after the objective with a microscope slide power sensor (S170C, Thorlabs) while ramping up the output voltage of the laser trigger box. e) Then, a monotonous polynomial is fit to the data and used as a look-up-table to set the laser trigger output voltage. This ensures that the actual laser power corresponds exactly to the user-requested percentage of the maximum laser power, providing accurate and reliable results for fluorescence experiments. f) The polynomial coefficients are stored in the K2 equipment configuration.

**Figure S51.**
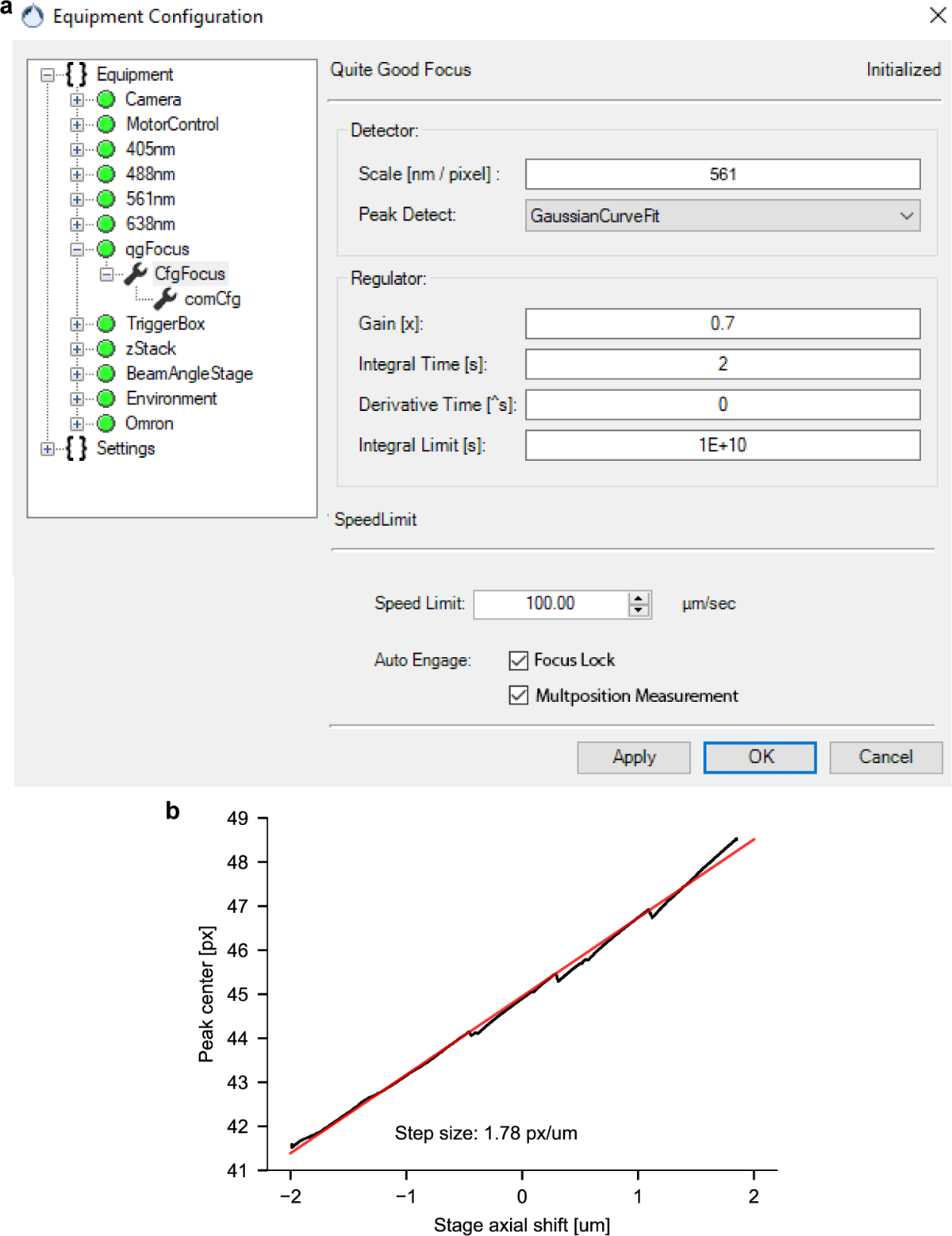
qgFocus configuration: a) Setting appropriate parameters for the focus stabilization feedback loop is essential to ensure a fast correction with little overshoot or oscillations. Other peak detection options than the Gaussian curve fitting, such as a simple maximum detector and a weighted average detector, are available. In our system, the Gaussian curve fit proved to be a stable option. PID feedback loop parameters were determined experimentally. A speed limit for the sample stage can be set and enforced during focus lock engagement, and/or an automated multiposition measurements, to avoid introducing Z-displacements that can’t be followed quickly enough by the qgFocus. b) The scaling factor to translate between the displacement of the laser beam on the sensor, and the Z-drift of the sample, is determined by extracting the slope of a linear fit to the positions of the center of the infrared laser while performing a Z-stack over an axial range of 4 µm. The jumps in the line are due to undersampling of the back-reflected laser beam - during the operation of the qgFocus, this did not appear to cause problems but could likely be fixed by changing the distance between the sensor and the lens in the qgFocus detection pathway.

**Figure S52.**
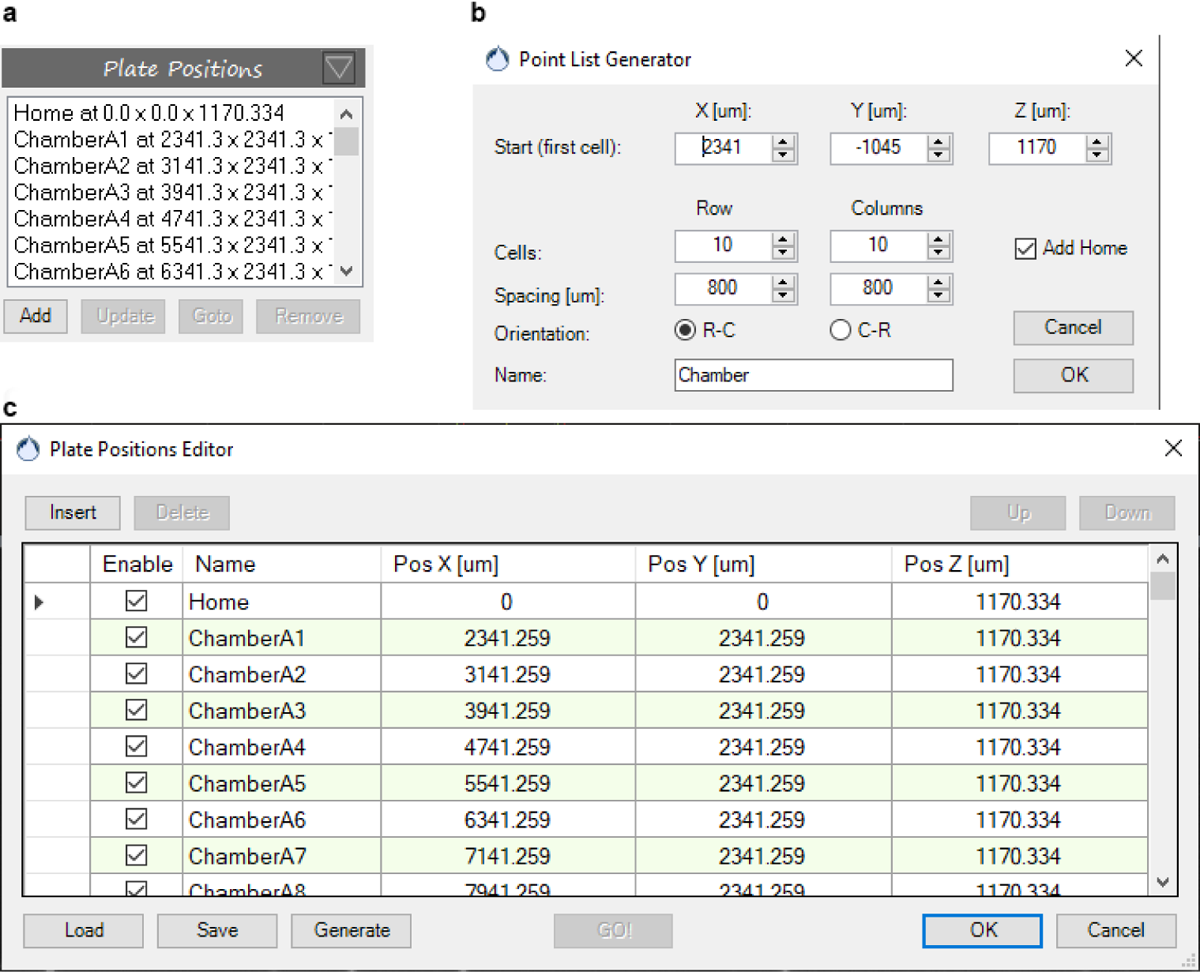
Point list generator: Plate positions (a) are created either manually by adding the current position of the stage or by loading a text file with coordinates and names. For automated raster scanning or the imaging of multi-well chambers, a point list generator (b) helps generating the plate positions (c).

**Figure S53.**
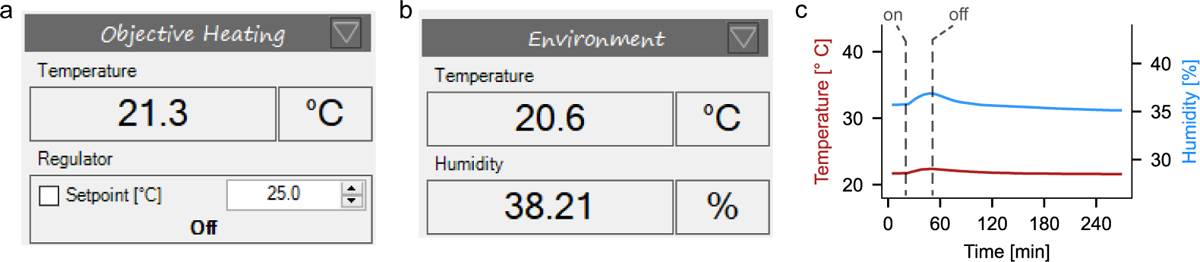
Environmental parameters and objective heating: **a**) In the objective heating panel, the objective temperature measured by a separate temperature sensor is displayed and the setpoint for the objective heating is specified. **b**) An additional temperature and humidity sensor is placed on the surface of the main cube. The temperature and humidity readings are displayed in the Environment panel and logged during experiments. **c**) The environmental sensor measures an increase in temperature during objective heating to 42 ^◦^C, likely due to radiative and convective heat transfer from the objective, and residual heat conduction through the thermal insulation spacer. The increase in humidity is explained by evaporation from the heated open sample chamber.

**Figure S54.**
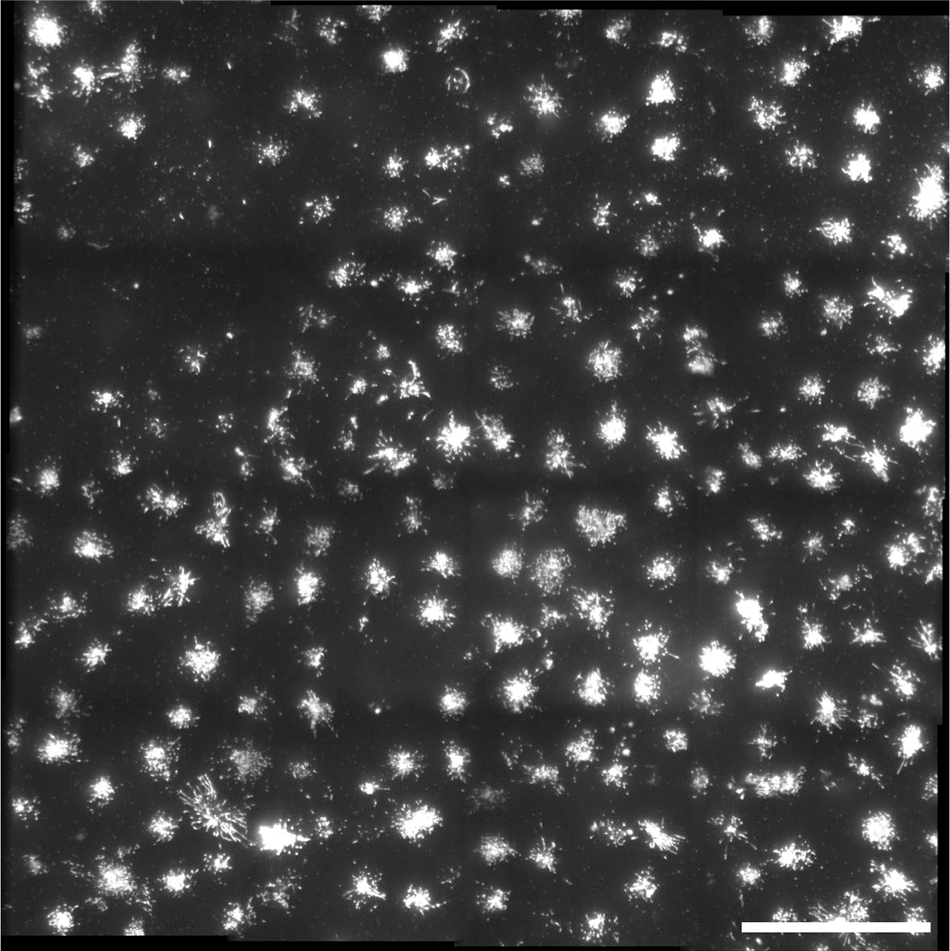
Multiposition measurements Four-by-four multiposition tile-scan of adherent cells seeded on fibronectin-coated coverslides, labeled with CellMask Deep Red Plasma membrane stain. The spatial overlap of the tile scan was chosen to be 15%. Stitching was done using the Stitching plugin [42] in Fiji [43]. Scale bar: 50 µm.

**Figure S55.**
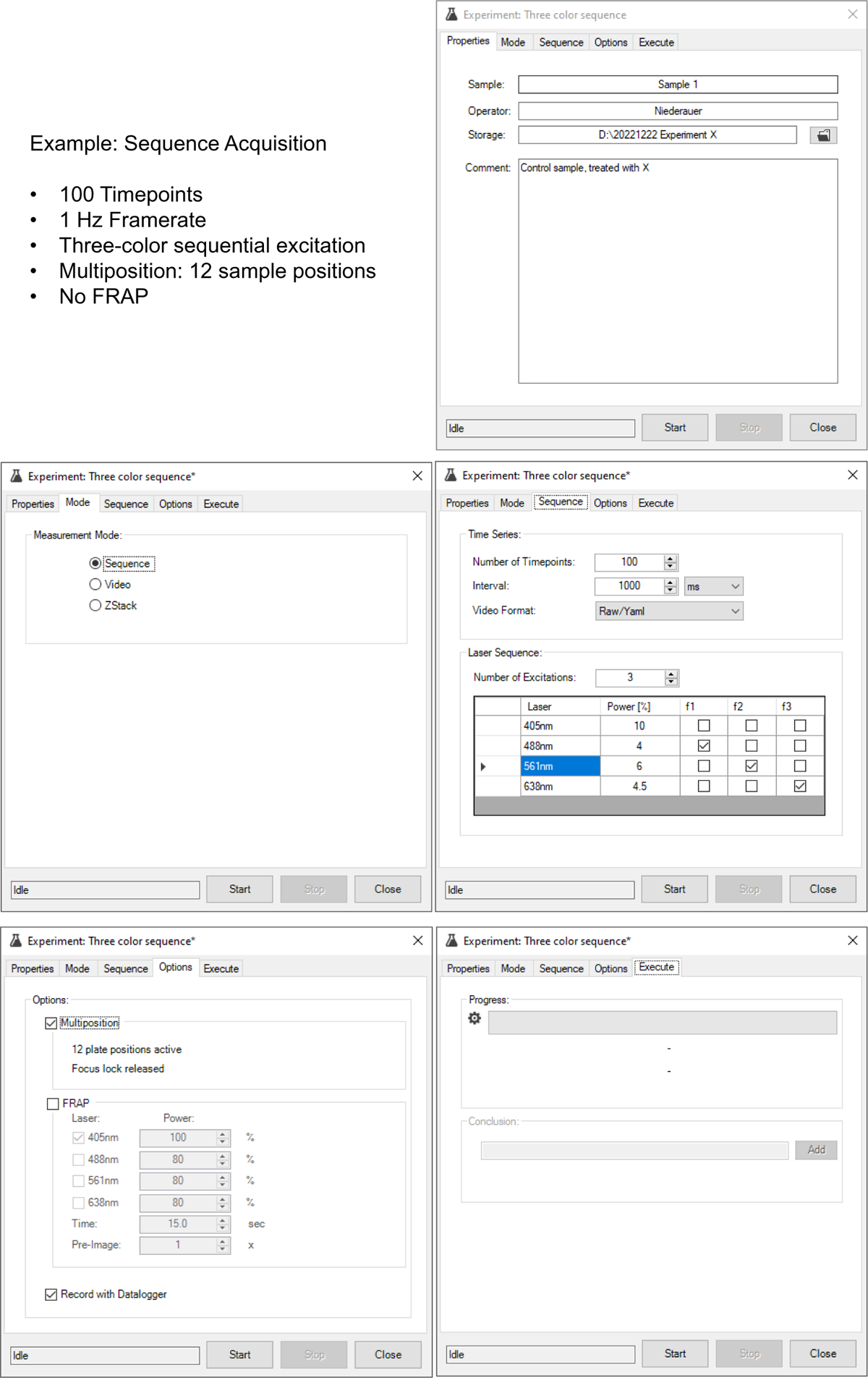
Experiment configuration: Sequence Acquisition Pre-configured three-color sequence acquisition of 100 timepoints with a 1 s interval. For each timepoint, three frames are acquired with 488 nm, 561 nm, or 638 nm laser excitation. The acquisition is repeated at 12 plate positions.

**Figure S56.**
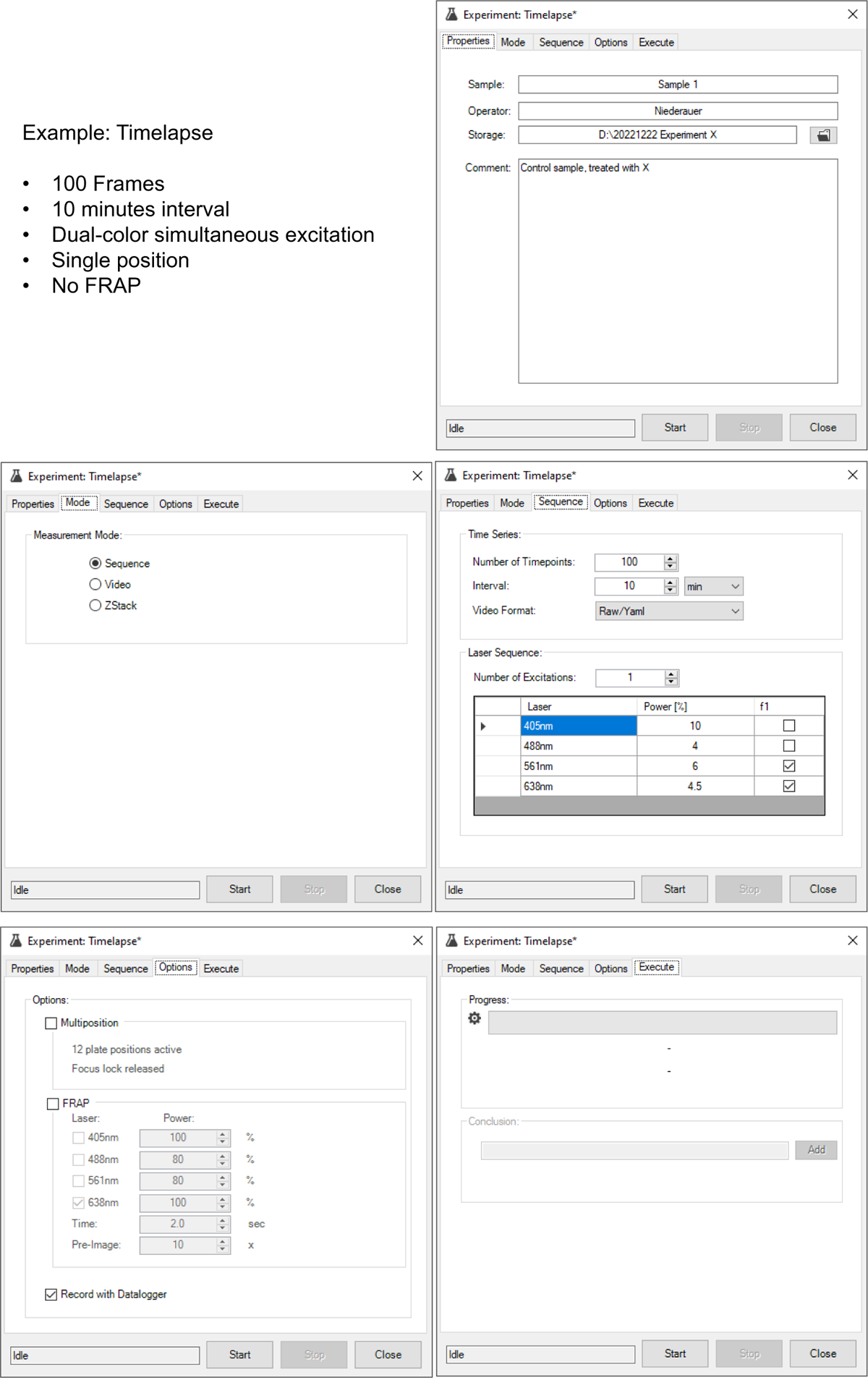
Experiment configuration: Timelapse acquisition Pre-configured timelapse acquisition of 100 timepoints with an interval of 10 min. At each timepoint, a single frame is acquired with 561 nm and 638 nm laser excitation.

**Figure S57.**
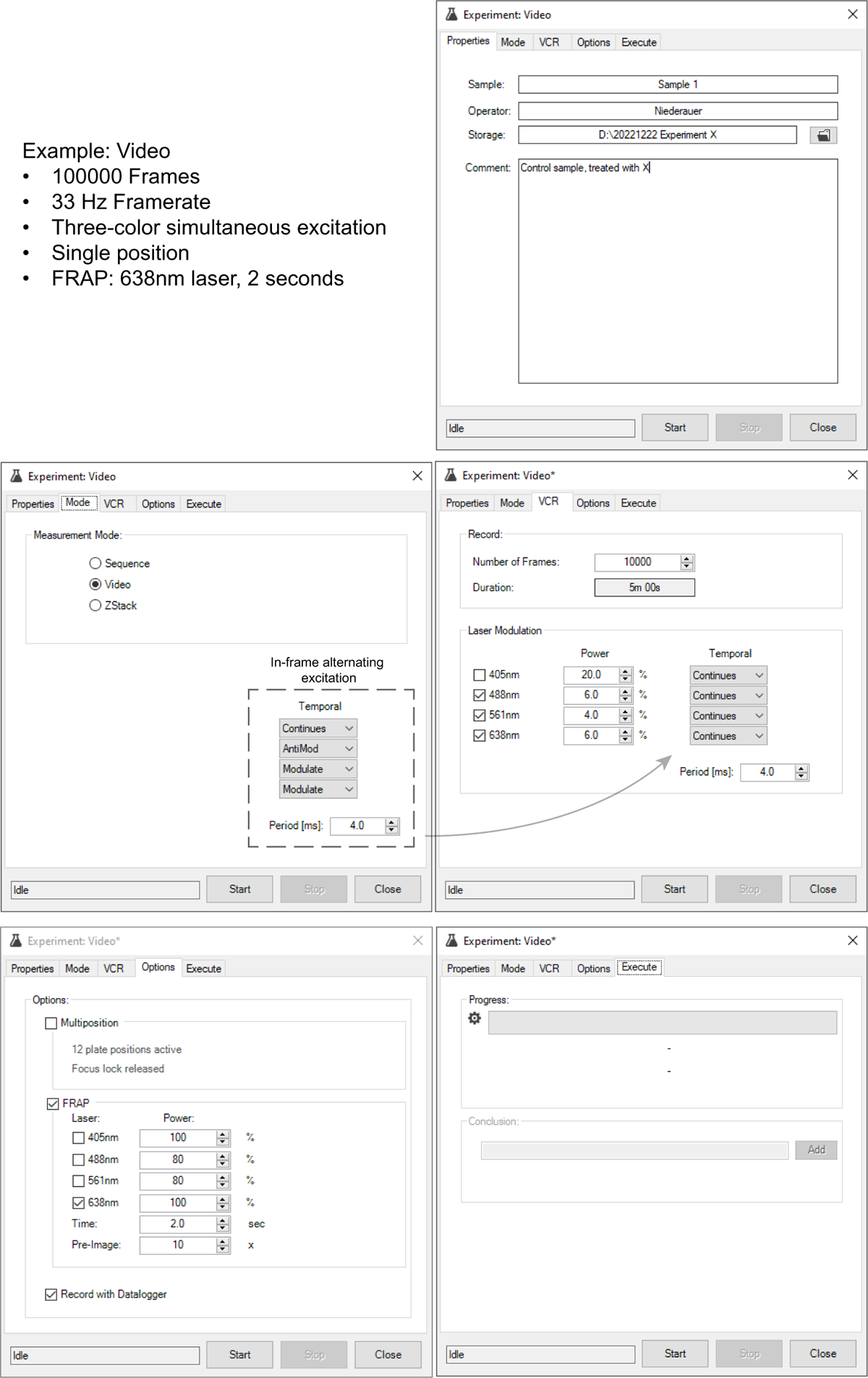
Experiment configuration: Video acquisition Pre-configured three-color video acquisition of 10,000 frames with 33 Hz framerate and an initial FRAP measurement. In the dashed box, the configuration for in-frame alternating excitation is shown, to reduce 488 nm laser-induced bleaching of dyes excited simultaneously by the 561 nm or 638 nm lasers.

**Figure S58.**
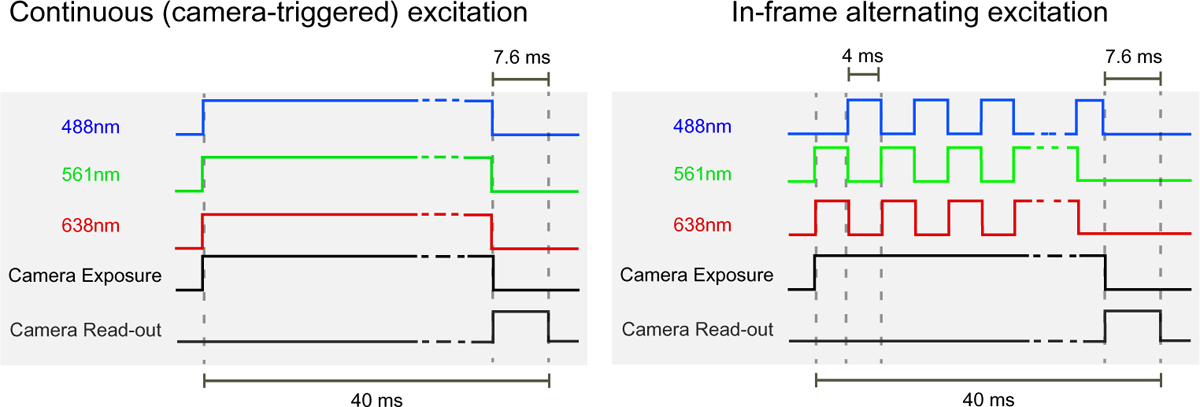
Left: Example of one frame cycle of a standard three-color laser excitation scheme, with laser excitation triggered by the camera exposure at a framerate of 25 fps. **Right:** In-frame alternating laser excitation: Laser pulsing is triggered by the camera exposure, with the 488 nm laser pulsing anti-cyclic compared to the 561 nm and 638 nm lasers. The pulse duration is set to 4 ms. For fast moving emitters, longer pulse durations would lead to recording the emission at increasingly divergent positions for the different color channels.

### Supporting Movies

**Figure SM1.**
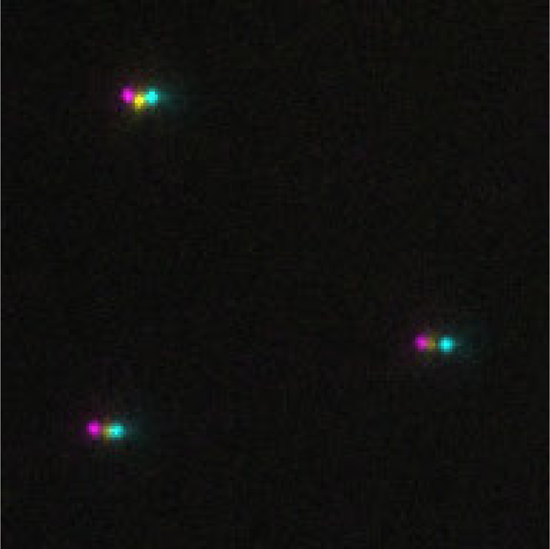
Three-channel superposition of multicolor fluorescently labeled beads imaged with the triple-color pathway, before precise *x* − *y* alignment. The sample stage is continuously swept along the *Z*-axis, to facilitate the axial co-alignment of the individual lenses in the three separated color channels.

**Figure SM2.**
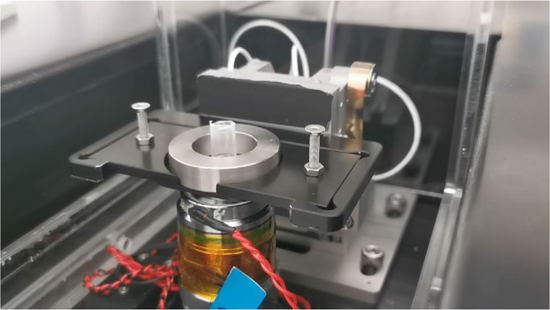
Sample ejection and mounting using the *Z*-toggle. To avoid introducing air bubbles, and decrease the amount of oil that needs to be supplied during multiposition measurements, the sample is slowed down when leaving and approaching contact with the immersion oil.

**Figure SM3.**
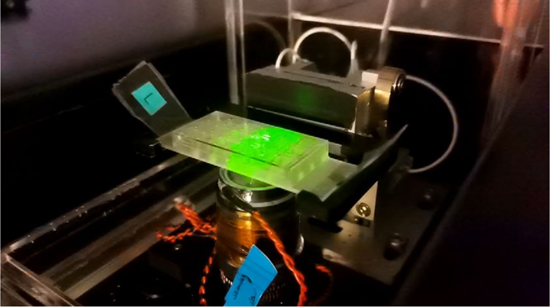
Sequential imaging of the wells of a 18-well microscopy slide using the multiposition acquisition option in the K2 software. Replay speed is 20x.

**Figure SM4.**
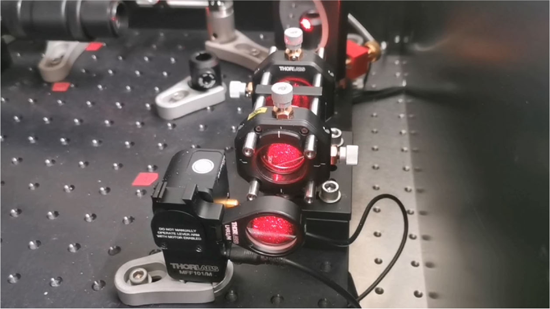
FRAP measurement: First, a series of pre-FRAP images is taken, then the flip-in lens moves into the optical path. After the bleaching step of 2 s, the lens returns to it’s default position and the imaging continues.

**Figure SM5.**
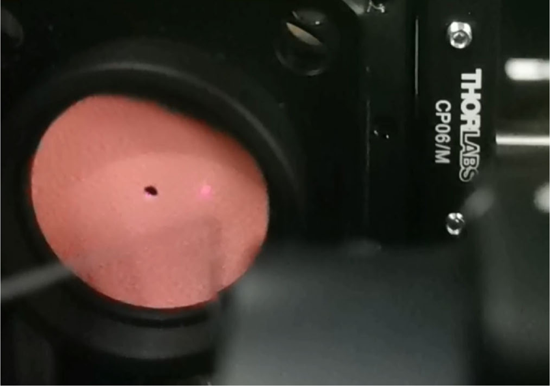
Back-reflection of the focus stabilization infrared laser beam, visualized on a infrared viewing disk. The sample is continuously sweeping up and down the *Z*-axis, which translates into a lateral movement of the back-reflection on the viewing disk.

## Notes

### Competing Interest Statement

The authors have declared no competing interest.

### Summary of Updates

Added Supplementary Information and Video Files.

