## Supplementary figures and images for "The K2: Open-source simultaneous triple-color TIRF microscope for live-cell and single-molecule imaging"

### Supplemental Video 1

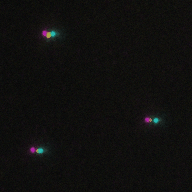
